# Computational Fluid Particle Dynamics-Informed Machine Learning Prototype for a User-Centered Smart Inhaler Enabling Uniform Drug Delivery to Small Airways

**DOI:** 10.64898/2026.03.16.712264

**Authors:** Ziyang Zhang, Hang Yi, Arun Varghese Kolanjiyil, Chenang Liu, Yu Feng

**Author notes:** Corresponding Author: Address: 354 Engineering North, Stillwater, OK 74078, USA. Corresponding Author: Address: 420 Engineering North, Stillwater, OK 74078, USA. Co-first Author.

## Abstract

Small airways are the primary sites of airflow obstruction in chronic obstructive pulmonary disease. Effective delivery of aerosolized drug particles to these regions is crucial to maximize treatment efficacy while minimizing side effects. However, conventional inhalation therapy approaches (i.e., full-mouth particle release and inhalation (FMD)) typically result in insufficient drug deposition in the small airways and an uneven distribution across the five lung lobes. To address such deficiencies, the goals of this study are triple folds: (1) to develop a fast and accurate framework to secure target drug delivery (TDD) nozzle diameter and location based on the conventional computational fluid particle dynamics (CFPD)-FMD simulations, (2) to develop a CFPD-informed machine learning (ML) inverse-design framework that predicts optimal inhaler nozzle parameters based on patient-specific breathing patterns and drug properties, and (3) to demonstrate the feasibility of embedding this framework into a user-centered smart inhaler prototype to improve uniform TTD to the small airways across all five lung lobes. Specifically, a subject-specific mouth-to-generation-10 human respiratory system was employed, and 108 high-fidelity CFPD-FMD simulations were performed under varied physiological and design parameters, including tidal volume, particle diameter, release location, and release timing. Particle release maps generated from those CFPD-FMD simulations *via* backtracking identified optimal nozzle diameters and locations that promote uniform multi-lobe drug delivery while limiting off-target deposition. Accordingly, a dataset was compiled with inputs (i.e., flow rate, particle size, release z-coordinate, release time) and targets (i.e., nozzle center x- and y-coordinates, nozzle diameter). These inputs and targets form the CFPD-TDD dataset, on which 16 ML models were trained to learn inverse mapping from patient- and drug-specific inputs to optimal nozzle design parameters. Performance was evaluated using mean squared error (MSE) and mean absolute error (MAE) overall and per target feature. Parametric analysis using CFPD-FMD simulations was conducted to determine how patient-specific and drug-specific factors affect pulmonary air-particle transport dynamics and to explain why achieving CFPD-TDD in small airways with CFPD-FMD strategies remains challenging. Furthermore, the ML evaluation in this feasibility study demonstrated robust learning of the inverse mapping from patient-specific inputs to optimal nozzle parameters. Four top-performing models showed consistently low MSE/MAE across cases, and an ensemble (i.e., mixed model (MixModel)) combining their strengths was formulated. Independent CFPD-TDD simulations beyond the training and testing datasets were used as the ground truth to validate ML-predicted nozzle configurations. Compared with conventional CFPD-FMD strategies, ML-guided nozzle designs significantly improved inter-lobar deposition uniformity and reduced off-target deposition in the upper airways, demonstrating the feasibility of ML-enabled TDD to the small airways. Overall, this study establishes a CFPD-informed ML inverse-design framework as a viable algorithmic foundation for user-centered smart inhalers, enabling adaptive, patient-specific TDD to the small airways with improved deposition uniformity across all five lung lobes. By integrating first-principle-based CFPD with ML, this work provides a methodological pathway toward next-generation smart inhalers for more effective treatment of small airway diseases.

## 1. Introduction

The term “small airway disease” was introduced in 1968, being marked as a primary contributor to increased airway resistance in individuals with obstructive lung disease (Hogg et al., 1968). It is reported that over 75% of small airway diseases are categorized as chronic obstructive pulmonary disease (COPD) (Crisafulli et al., 2016). Indeed, COPD is a significant and increasingly prevalent global health issue. By 2050, the number of COPD cases is expected to reach a total of 592 million, which marks a 23.3% increase compared to the statistics in 2020 (Boers et al., 2023). Moreover, COPD is becoming a more frequent reason for hospital stays and work absences, leading to substantial healthcare expenses, and the costs now surpass those linked to asthma by over three times. In the United States, the financial burden of COPD is expected to rise significantly over the next two decades, with annual costs nearing $40 billion, indicating an expense of around $800 billion (Soriano et al., 2020). Therefore, patients with small airway disease, particularly those with COPD, bring more severe global health challenges, which contribute to economic strain and increase daily-life complications. This underscores the importance of efficient pulmonary disease treatment that can deliver medications to small airways.

The small airways, often referred to as peripheral airways, are air passages less than 2 mm in hydraulic diameter, comprising bronchioles, terminal bronchioles, respiratory bronchioles, alveolar ducts, and alveolar sacs (Usmani et al., 2021). Conventional approaches for evaluating airway function, including forced expiratory volume in one second (FEV1) and peak expiratory flow (PEF), are affected by the level of airway resistance (Aggarwal et al., 2006). A study found that the risk of death and disability from COPD is highly linked with a decline in lung function over time, with a loss in FEV1 exceeding 50 ml per year, which is significantly higher than the typical annual decline of 20 ml (Fletcher & Peto, 1977). Another study concluded that a decrease in PEF value by 49 L/min (27% from baseline) was significantly associated with an increased risk of COPD exacerbations and hospitalization (Cen & Weng, 2022). This gradual decline in FEV1 and PEF worsens the disease over time, leading to a progressive increase in shortness of breath during physical activity and eventually progressing to respiratory failure.

In the management of COPD exacerbations, inhalation therapy plays a crucial role, as international guidelines recommend the use of bronchodilators and anti-inflammatory medications at all stages of the disease (Halpin et al., 2021). The concept of inhalation therapy was introduced by Philip Stern in 1764, who argued that delivering medicine directly to the lungs could only be achieved *via* the windpipe (Stern, 1767). Later, in 1778, English physician John Mudge coined the term “inhaler” in his book “*A Radical and Expeditious Cure for a Recent Catarrhous Cough*”, where he introduced an inhaler crafted from a pewter tankard to deliver opium vapor for cough relief (Mudge, 1779). Since then, the rise of scientific rigor and regulation in the pharmaceutical industry has preserved foundational principles, though most early inhaler devices have evolved or disappeared (Sanders, 2007). An ideal inhaler should deliver a consistent, precise dose of drug particles to the diseased area (i.e., designated lung sites) of the respiratory system. It must be user-friendly to minimize errors, while remaining affordable and eco-friendly. Additionally, its design should ensure compatibility with different medications to maximize drug effectiveness and patient adherence (Sorino et al., 2020). Today, the most used inhalers fall into four categories: nebulizers, dry powder inhalers (DPIs), pressurized metered-dose inhalers (pMDIs), and soft mist inhalers (SMIs). Despite various advancements in these four types of inhaler technology and drug formulation, these devices are often designed independently of the specific medications they deliver (Cazzola et al., 2020). Studies also showed that nearly 60% of patients struggle with adherence, frequently due to unintentional inhaler handling errors that compromise treatment effectiveness (Krigsman et al., 2007; Molimard et al., 2003). The situation becomes worse when healthcare providers attribute poor clinical outcomes to the drug’s pharmacological properties while overlooking the possibility of improper inhaler use. This often leads to unnecessary dosage increases, resulting in significant toxicity and more severe side effects. Consequently, the market lacks an inhaler that meets all the criteria for an ideal device.

Nevertheless, many researchers have focused on developing new drugs that can be delivered effectively via inhalers to alter the progression of COPD. Inhalation is considered a highly advantageous route of administration, offering significant advantages over oral, transdermal, or intravenous methods, such as faster onset of action (Mash et al., 1996), lower doses with reduced side effects (Labiris & Dolovich, 2003), comparatively better drug concentrations in the lungs (Patton & Byron, 2007), and non-invasive administration (Veloso et al., 2018). Despite the advantages of inhalation therapy, treating small airway diseases such as COPD is still challenging. In COPD patients, chronic inflammation in the peripheral lungs thickens small airway walls through squamous metaplasia and goblet cell hyperplasia, while inflammatory exudates obstruct the lumen, ultimately narrowing the airways and reducing ventilation (Hogg et al., 2004). The exacerbation of COPD symptoms mentioned above can hinder the efficient deposition of inhaled therapeutic particles in the small airways, as airway obstruction alters airflow patterns and can further impair drug delivery (Usmani, 2014). Additionally, the anatomical structure of the airways plays a critical role in determining both the efficiency of drug delivery and the deposition site of inhaled medications, whether primarily in the larger conducting airways or the peripheral regions of the lungs (Verma et al., 2015). In fact, most traditional inhalation therapies achieve only 10-40% lung deposition, with a large portion of the drug deposited in the upper airways (Yang et al., 2014). The unnecessary drug deposition in the upper airways leads to suboptimal therapeutic outcomes and undesirable side effects due to off-target deposition on healthy tissues (O’Callaghan, 1994). Therefore, improving the precision of drug delivery is essential to maximize therapeutic particle deposition in small airways while minimizing unwanted deposition in healthy tissues.

Over the past few decades, targeted drug delivery (TDD) within the airways has shown remarkable potential to enhance therapeutic efficacy. TDD approaches offer significant advantages over conventional inhalation therapies by minimizing drug deposition in healthy tissues and reducing the risk of side effects, while specifically targeting diseased airway regions (Bae & Park, 2011). Additionally, TDD enables the sustained release of therapeutic agents, providing prolonged exposure to target tissues with a key advantage in managing chronic lung diseases (Tayab & Hochhaus, 2005). Numerous approaches were employed to achieve TDD, e.g., (1) magnetic targeting, which uses external magnetic fields to direct drugs attached to paramagnetic carriers (Martin & Finlay, 2008), (2) nano-carrier-based drug, in which nanoparticles serve as a transporting agent for drug particles (Mishra et al., 2010), and (3) a controlled air-particle stream method by determining the suitable particle release position *via* the so-called “backtracking” strategy (Kleinstreuer & Zhang, 2003; Kleinstreuer et al., 2008). The key factors for achieving TDD *via* the pulmonary route are the properties of the drug, including its size, density, and composition (Finlay, 2001; Kleinstreuer et al., 2014). However, it is also crucial to understand and control aerosol dynamics in the respiratory tract to ensure effective drug delivery to specific lung regions, beyond the scope of drug formulation engineering. Specifically, subject-specific complex and dynamic lung environments make it difficult to deliver drugs uniformly and precisely. Direct *in vivo* experiments for TDD in the lungs are limited in their ability to visualize the transport process, are costly and time-consuming, and are always constrained by numerous ethical concerns. As an alternative, the experimentally validated computational fluid particle dynamics (CFPD)-based *in silico* study offers a non-invasive, efficient, and high-resolution approach for studying key variables that can influence airflow dynamics and particle transport (Islam & Feng, 2023; Islam et al., 2024; Kleinstreuer & Zhang, 2003).

Specifically, CFPD has been pivotal in understanding the pulmonary fluid-particle dynamics of drug aerosol transport and in advancing respiratory drug delivery (Walenga et al., 2019). CFPD is first principles-based and can accurately predict pulmonary air-particle flow dynamics in subject-specific 3D airway geometries, with advanced capabilities for simulating particle-air interactions, liquid-gas phase changes, and particle-particle interactions. Recent advancements in CFPD have expanded its use across numerous engineering disciplines, including the fundamental understanding of the complex mechanisms involved in respiratory drug delivery. Longest et al. (Longest et al., 2019) conducted an extensive review of the application of CFPD techniques to advance pulmonary drug delivery, covering areas such as aerosol transport and deposition, inhaler performance evaluation, and the development of effective drug delivery strategies. In the context of enhancing pulmonary targeted drug delivery, CFPD researchers employed various strategies. The concept of “controlled particle release” for direct drug delivery was first demonstrated by Kleinstreuer and Zhang (Kleinstreuer & Zhang, 2003), who developed and validated a CFPD method for targeting tumors by optimizing radial inhaler positioning based on particle release maps.

Later, Childress and Kleinstreuer (Childress & Kleinstreuer, 2014) redefined the development of particle release maps using CFPD simulations, which serve as descriptive tools for identifying optimized drug delivery locations at the inlet. Specifically, the particle “backtracking” method identifies release positions at the mouth corresponding to specific lung deposition sites, enabling precise drug delivery to targeted lung regions. For example, Feng et al. (Feng, Chen, et al., 2018) demonstrated that lobe-specific drug delivery could be improved by up to 90% using a particle release map-based targeted delivery strategy compared with conventional inhalation methods, in which drug particle release position and velocity were optimized. Another recent CFPD study used a similar TDD concept to effectively deliver aerosolized chemotherapeutic particles to small airway tumors via endotracheal catheters and demonstrated significantly improved drug delivery efficiency at selected generation 10 (G10) terminals (Islam & Feng, 2023).

Despite these advances, achieving effective TDD in small airways remains a significant challenge that has not been systematically explored. Delivering a sufficiently high and evenly distributed drug dose across the five lung lobes is difficult, particularly because the “backtracking” method using CFPD is case-specific and computationally intensive. Any change in airway geometry, patient-specific breathing pattern, or drug-specific particle characteristics (e.g., size or density) requires new CFPD simulations. Consequently, there is an urgent need for a fast, adaptive, and user-centered optimization framework that can translate the rich physics-based insights of CFPD into real-time inhaler guidance for individualized therapy.

To address the need mentioned above, the present study proposes a CFPD-informed machine learning (ML) framework that serves as the foundation for a user-centered smart inhaler prototype (Islam et al., 2024) designed to enable uniform and efficient drug delivery to the small airways. The ML models are trained on first principles-based CFPD data to predict optimal drug-release nozzle configurations, including in-plane center coordinates and nozzle diameters, based on patient-specific breathing patterns (e.g., inhalation flow rate, particle release timing), inhaled drug-specific properties (e.g., particle diameter), and inhaler insertion depth (see **Fig. 1**). The goal is twofold, i.e., (1) to maximize drug transport to the peripheral lung regions beyond generation 10 (G10), and (2) to minimize deposition disparities among the five lung lobes, promoting spatially uniform delivered drug particle distribution.

**Figure 1:**
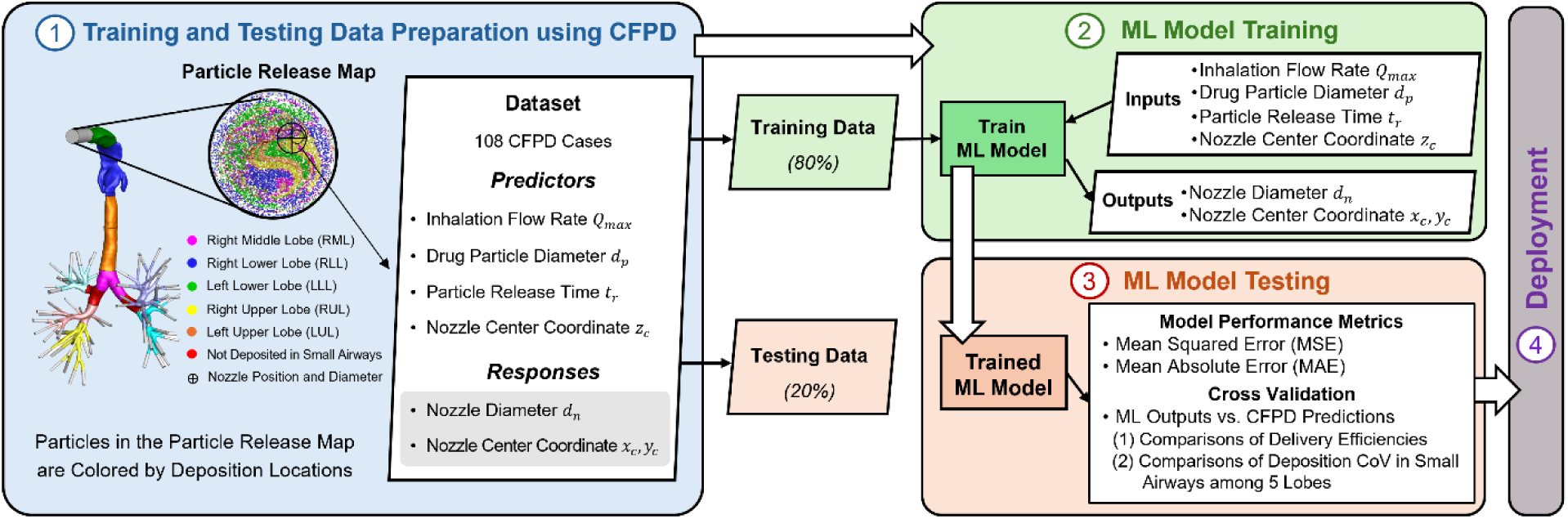
Illustration of the ML algorithm for achieving the targeted drug delivery from data collection to model training and testing.

In this study, a subject-specific airway geometry extending from the mouth to G10 has been employed in simulations using an experimentally validated CFPD model to generate training and testing data. In the envisioned innovative inhaler system integrating an ML algorithm (see **Fig. 2**), the signal acquisition module first collects patient breathing profiles, particle sizes, and inhaler insertion depth, which are then used as input to the ML model. The model determines the optimal nozzle diameter and position, and this information is transmitted to an automated mechanism that adjusts the nozzle and releases the drug during the next inhalation of the patient. **Figure 1** illustrates how the CFPD model generates datasets for ML model training and testing.

**Figure 2:**
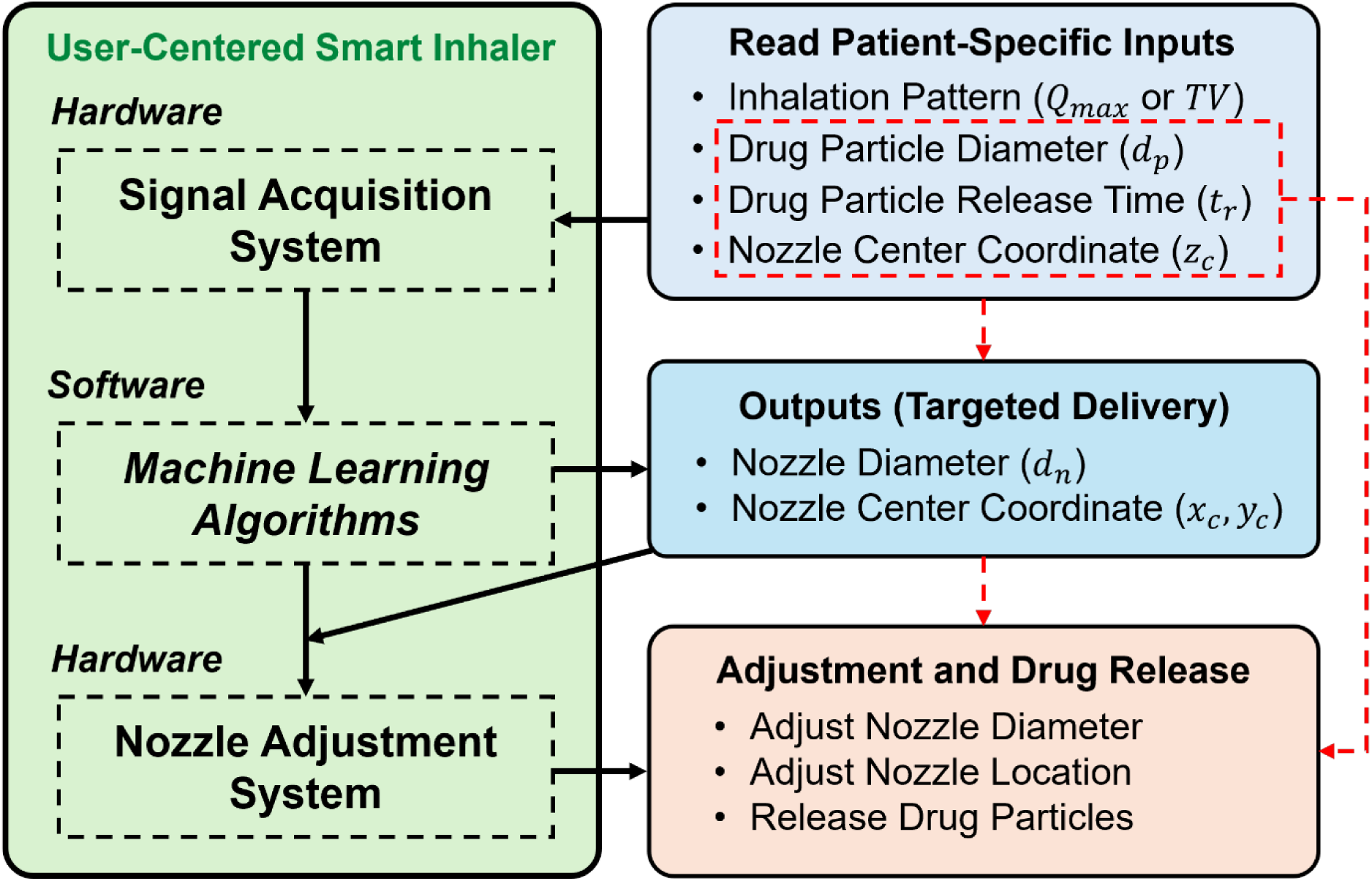
Schematic of the working principle for the user-centered smart inhaler prototype.

Meanwhile, **Fig. 2** illustrates how the smart inhaler integrates patient- and drug-specific data into the ML framework. Specifically, in the smart inhaler design, the outer mouthpiece diameter is kept constant to maintain user comfort and consistent inhalation behavior. The “nozzle diameter and location” predicted by the ML model, therefore, refer to an internal aerosol release orifice rather than a change to the overall mouthpiece size. In practice, this adjustment could be achieved using a small mechanical actuator, such as a sliding aperture or an iris-type mechanism, to modify the effective opening area and shift the aerosol’s release position within the mouthpiece. Because altering the orifice area affects the local jet velocity via conservation of mass, the patient’s inhalation flow rate is explicitly included as an input to both the CFPD simulations and the ML training process. As a result, the predicted nozzle settings reflect the combined influence of geometry, airflow velocity, and particle transport behavior under patient-specific breathing conditions.

The long-term goal is to develop an AI-powered smart inhaler that precisely delivers therapeutic aerosols to diseased regions of the lung, enabling safer, more effective, and patient-specific treatments. This study represents a step toward seamlessly integrating ML with CFPD to support TDD in inhalation therapy, with the potential to revolutionize individualized pulmonary drug delivery, especially for the treatment of small airway diseases such as COPD. It is worth noting that the proposed CFPD-informed ML framework is intentionally designed to be architecture-agnostic for inhalers, rather than developed for any single inhaler category. Instead, the current work focuses on developing a generalizable algorithmic prototype that can be integrated into various inhaler types.

## 2. Methodology

### 2.1 Geometry and Mesh

**Figure 3 (a)** shows a 3D subject-specific airway geometry extending from the mouth to G10 across five lobes, which was used in this study as a representative virtual human respiratory system. The five lobes are left upper lobe (LUL), left lower lobe (LLL), right upper lobe (RUL), right middle lobe (RML), and right lower lobe (RLL), respectively. This geometry was reconstructed from computed tomography (CT) images of a 47-year-old male volunteer with a height of 174 cm and a weight of 78 kg, who had no prior history of respiratory illnesses (Zhang et al., 2012). A circular mouth opening with a diameter 𝐷_𝑚_ of 20 mm was designed to replicate the size of the inhaler mouthpiece. The mouth opening is centered at origin (𝑥, 𝑦, 𝑧) = (0, 0, 0) and is oriented with the normal vector aligned with the positive z-direction. The tracheobronchial (TB) tree has 72 small airway terminals with an average hydraulic diameter of 1.91 mm and a median of 1.87 mm.

**Figure 3:**
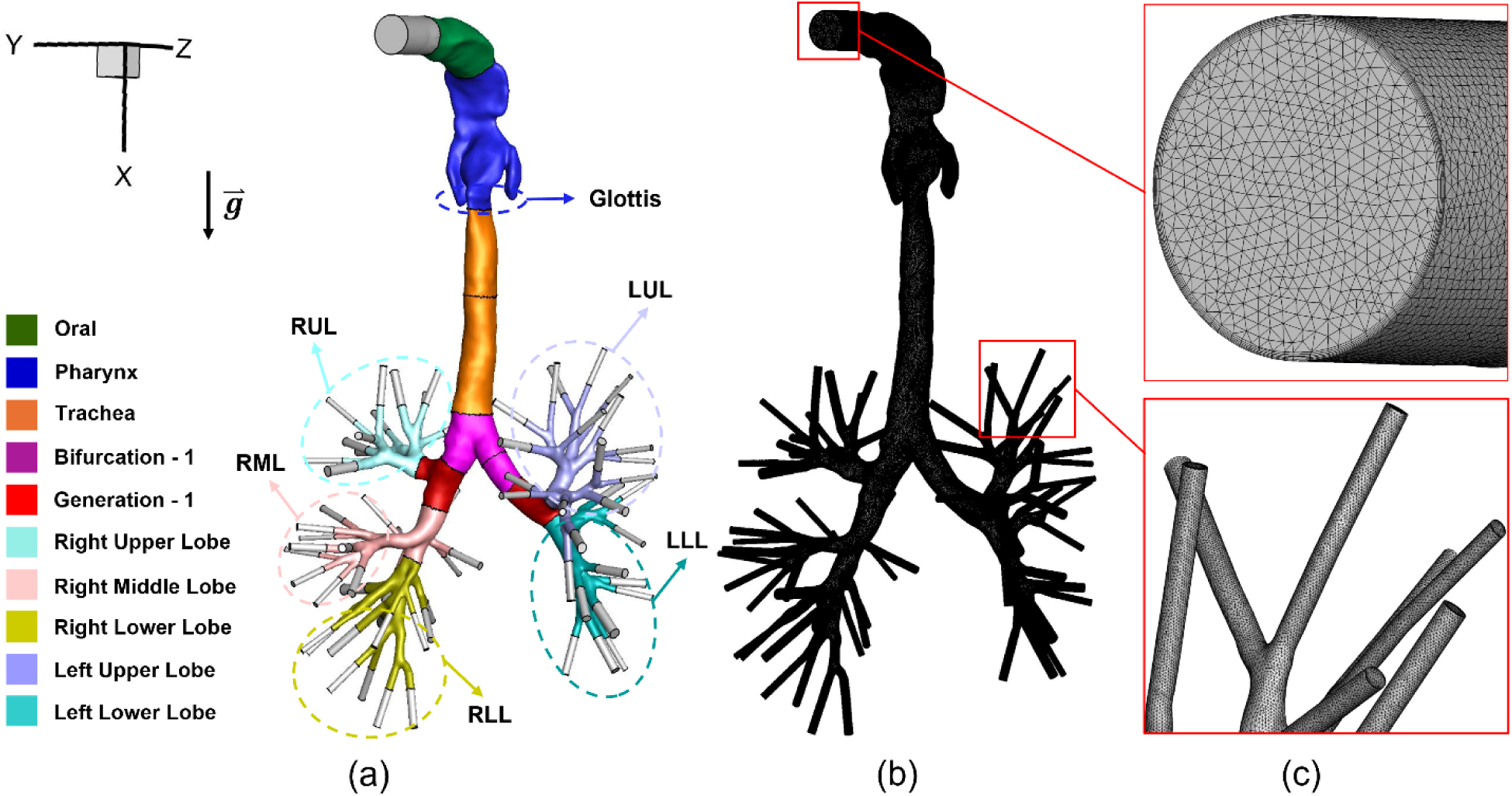
Geometry and mesh details of the 3D mouth to generation-10 (G10) geometry: (a) Schematic of the mouth opening with five lung lobes (D_in_=20 mm), (b) Tetrahedron-based mesh employed in this study, and (c) Mesh details at the mouth opening and small airways with six near-wall prism layers.

Tetrahedron-based meshes were created using Ansys Fluent 2023 R1 Meshing (Ansys Inc., Canonsburg, PA). Six near-wall prism layers were created with refined thickness to ensure 𝑦^+^ < 1 and to effectively capture laminar-to-turbulence transition regions, where 𝑦^+^represents the dimensionless wall distance (Menter et al., 2006). The final mesh (see **Figs. 3 (b) and (c)**) comprises approximately 7.7 million cells, 17.3 million faces, and 2.5 million nodes. The final mesh employed in this study was generated using a similar meshing strategy determined by mesh independence tests documented in existing publications for the same airway geometry (Feng et al., 2017; Feng, Zhao, et al., 2018).

### 2.2 CFPD Model

#### 2.2.1 Governing Equations

In this study, inhalation-exhalation waveforms with three different tidal volumes (𝑇𝑉), i.e., 300, 500, and 750 ml were employed to mimic the inhalation-exhalation patterns in the respiratory system (i.e., mouth-G10), respectively. The determined tidal volumes based on a predicted body weight (PBW) of 6 ml/kg, and the adult weight ranges from 50 to 125 kg was considered in this study (Cheung So & Chang, 2022; Network, 2000). Details can be found in Section 2.2.2. Since Reynolds number 𝑅𝑒≈2400 in the glottis region (see **Fig. 3 (a)**) at the instant of inhalation peak flow rate of 28.27 L/min when the lowest 𝑇𝑉=300 ml, the Transition Shear-Stress Transport (SST) model (Chen et al., 2017; Jiang et al., 2025; Li et al., 2025; Liu et al., 2024; Zhang & Kleinstreuer, 2011) was employed to accurately predict the flow patterns transitioning from laminar to turbulence in the upper airway. Pulmonary airflow was assumed to be incompressible. Accordingly, the continuity equation solved in this study can be given as

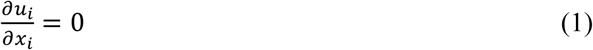

where 𝑢_𝑖_ is the airflow velocity.

The Navier-Stokes (N-S) equation represents the conservation of momentum, as shown below in **Eq. (2)**, where the viscous tensor 𝜏_*ij*_ is defined in **Eq. (3)**.

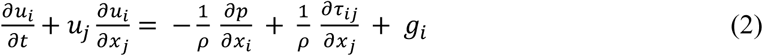

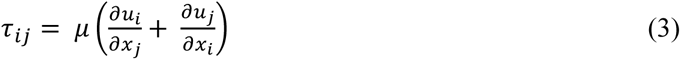

In **Eqs. (2) and (3)**, 𝜌 represents air density, 𝜇 denotes dynamic air viscosity, 𝑝 stands for pressure, and 𝑔_𝑖_ = (9.81, 0, 0) m²/s is the gravitational acceleration.

The inhaled drug particles were assumed to be monodisperse and spherical. The interaction between particles was considered negligible, and a one-way coupled Euler-Lagrange method was employed to predict particle transport from the mouth to G10, assuming the flow is primarily in the dilute regime. Four different sizes of particles were considered for this study, and the particle aerodynamic diameters (𝑑_𝑝_) are 0.5, 1, 2, and 5 µm, respectively. Additionally, the study considered drag force, Saffman lift force, and gravity as the primary forces acting on the particles, while the Brownian motion-induced force was neglected. For predicting particle trajectories, Newton’s 2nd law was employed and can be given as

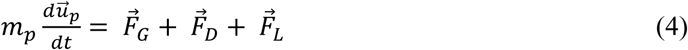

where 𝑚_𝑝_ and 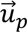 represent the particle mass and velocity, respectively. 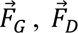 and 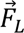 denote the gravitational force, drag force, and Saffman lift force. The drag force 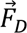 can be defined as

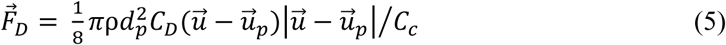

where 𝑑_𝑝_ is the particle aerodynamic diameter, 𝐶_𝑐_ represents the Cunningham correction factor, and 𝐶_𝐷_ refers to the drag coefficient (Chen et al., 2018), which can be given as

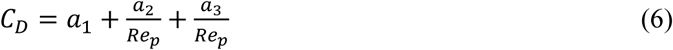

Constants 𝑎_1_, 𝑎_2_, and 𝑎_3_ are dependent on the particle Reynolds number, 𝑅𝑒_𝑝_, which is defined as (Morsi & Alexander, 1972)

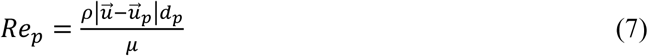

The Cunningham correction factor 𝐶_𝑐_ in **Eq. (5)** can be given by (Allen & Raabe, 1985)

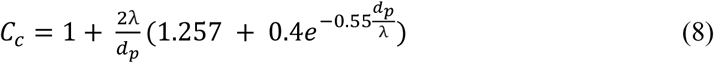

where 𝜆 is the mean free path of air.

The Saffman lift force, 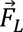 is particularly significant for particles with a larger size (Kallio & Reeks, 1989). The tensor form of the Saffman lift force 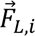_𝑖_ can be given as (Li & Ahmadi, 1992)

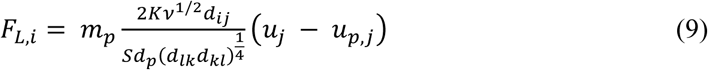

where 𝐾 = 2.594 is the constant coefficient for the Saffman lift force, 𝜈 represents the kinematic viscosity, 𝑆 is the ratio of particle density to fluid density, and 𝑑_*ij*_ is the deformation rate, defined as

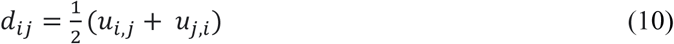

#### 2.2.2 Initial and Boundary Conditions

A CFPD parametric study was conducted to determine how factors (i.e., particle size, release location, and release time) influence targeted drug delivery to small airways in the lung. The CFPD results served as training and testing datasets for the next step, e.g., developing the prototype of the user-centered smart inhaler CFPD-ML framework. In total, 108 CFPD simulations were designed and conducted to produce the necessary data. **Table 1** summarizes the design of the parametric study. Specifically, the CFPD simulations examined how factors like particle diameter (𝑑_𝑝_), peak inhalation flow rate (𝑄_𝑚𝑎𝑥_), particle release time (𝑡_𝑟_), and particle release position (𝑧_𝑐_) influenced the efficiency and evenness of drug delivery to the small airways beyond G10. Therefore, for training and testing, the outcomes of 108 CFPD simulations, along with mapped data, were combined to form data samples, yielding the dataset used for ML model development. The training and testing dataset structures are shown in Fig. 1 and **Appendix A** (see Section 2.3 for more details).

**Table 1:** Parametric designs for CFPD simulations to generate the training and testing dataset.

| Peak Inhalation Flow Rate<br>($Q_{max}$) [L/min] | Particle Diameter<br>($d_p$) [$\mu\text{m}$] | Z-Coordinate of Particle<br>Release ($z_c$) [m] | Particle Release<br>Time ($t_r$) [s] |
| --- | --- | --- | --- |
| 28.27 (TV = 300 ml) | 0.5 | 0.001 | 0 |
| 47.12 (TV = 500 ml) | 1 | 0.02 | 0.25 |
| 70.69 (TV = 750 ml) | 2 | 0.04 | 0.5 |
|  | 5 |  |  |

Specifically, with the same inhalation and exhalation time for all simulations, three transient inhalation-exhalation flow waveforms were designated as idealized sinusoidal, as shown in **Fig. 4**. The breathing cycle time 𝑇 was set to 2 seconds. In the three breathing patterns, the 𝑇𝑉s were 300, 500, and 750 ml, with corresponding 𝑄_𝑚𝑎𝑥_ = 28.27, 47.12, and 70.69 L/min (see **Table 1**). Thus, the transient inhalation-exhalation flow rate, 𝑄_𝑖𝑛_ can be given by 𝑄_𝑖𝑛_

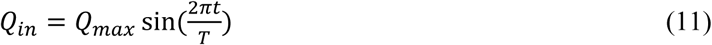

**Figure 4:**
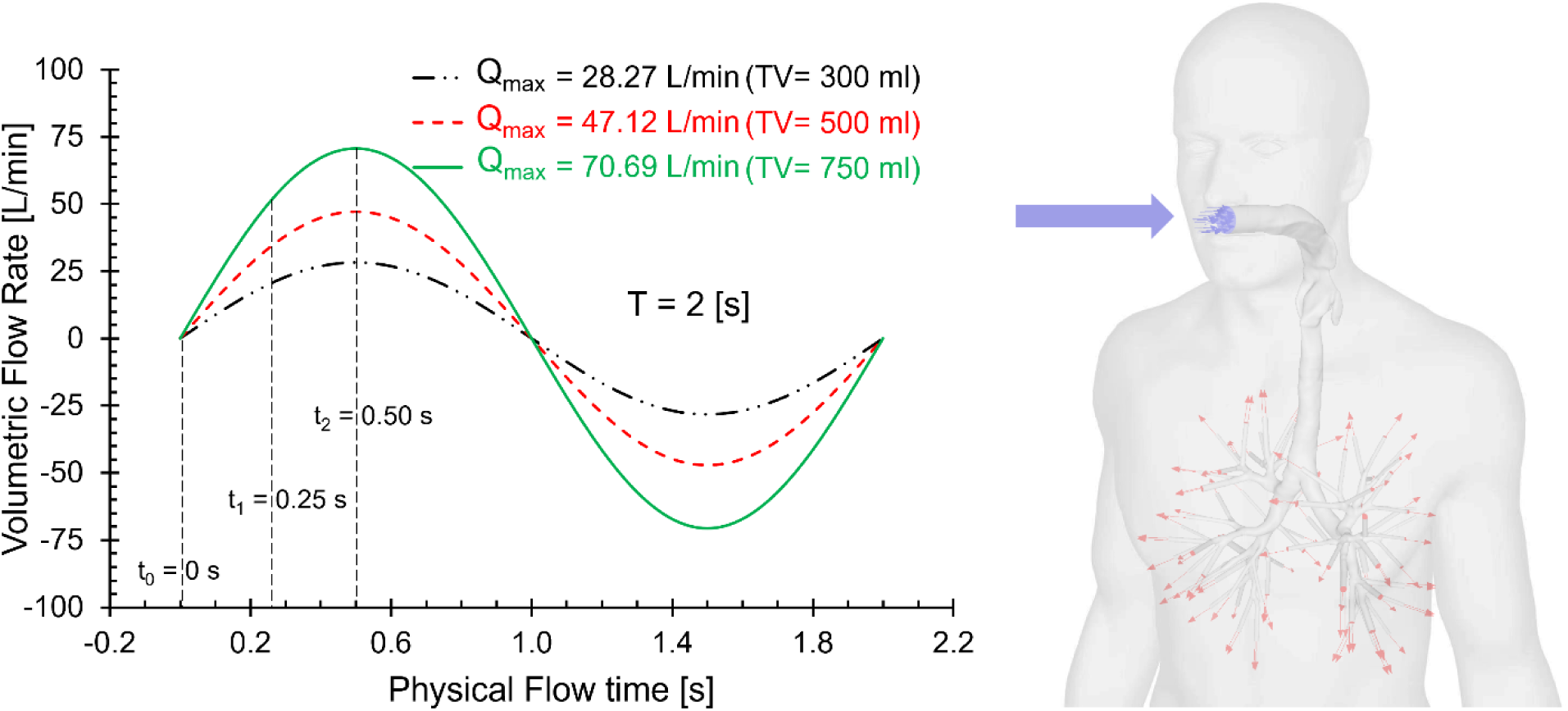
Sinusoidal inhalation-exhalation waveforms with three peak inhalation flow rates.

During the inhalation, a total of 10,036 particles, each with a constant diameter and a density of 1000 kg/m³, were injected as a bolus single time point injection from the mouth in the positive z-direction (see **Fig. 3**). A no-slip boundary condition is used on the airway walls. CFPD simulations in this study model the airways as surfaces covered with mucus layers by applying a “100% trapped wall” boundary condition for particle-airway interaction. This boundary condition results in particle deposition when the distance between the particle center and the airway wall is less than the particle radius. At the mouth opening and the small airway openings, an “escape” boundary condition was applied for particles, whereas the gauge pressure was set to zero at all small airway openings.

#### 2.2.3 Particle Delivery Efficiency

To quantify the delivery efficiency, the regional deposition fractions (DFs) of drug particles in the human respiratory system were employed, which can be defined as

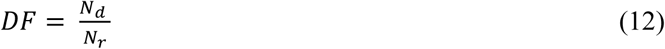

where 𝑁_𝑑_ the number of particles deposited in a specific region or the number of particles exited G10, and 𝑁_𝑟_ is the number of total particles released from the mouthpiece opening. It is worth noting that particles existing from G10 terminals were assumed to be deposited 100% in the downstream lobes, i.e., no particles can reenter the mouth-to-G10 domain after escaping.

### 2.3 Criteria and Workflow for Search Optimal Nozzle Configuration to Achieve TDD

To provide quantitative guidelines for determining the maximum allowable nozzle diameter and optimal nozzle location to achieve personalized TDD, this study developed novel criteria to determine the optimal nozzle diameter and location. Two primary criteria for seeking optimal TDD nozzle diameter and location are: (i) The first and highest-priority criterion is the best delivery uniformity, ensuring that particles are distributed as evenly as possible across the five lung lobes; (ii) The second criterion is maximizing the total number of particles delivered to the TB tree, which ensures effective drug delivery to the peripheral lung, which is the designated lung sites for this study. Based on these two criteria, this study developed a systematic framework to determine the optimal TDD nozzle location and diameter using particle-release maps generated from CFPD-FMD simulations (see **Appendix A**), which are subsequently used as ML training and testing datasets. The detailed framework and workflow for identifying the optimal nozzle diameter and location for TDD are shown in **Fig. 5**. The workflow can be described as follows:

1. *Step 1: Particle Release Map Reconstruction*. First, for each CFPD-FMD simulation (i.e., conventional inhalation therapy *via* full-mouth particle release), particle release information (i.e., particle initial release position at the mouthpiece opening) is obtained using an in-house Python code based on the particle backtracking strategy (Feng, Chen, et al., 2018; Islam & Feng, 2023; Kleinstreuer & Zhang, 2003). Specifically, particle release positions are classified by the deposition location (i.e., LLL, LUL, RLL, RML, or RUL), enabling reconstruction of the spatial distribution at the inhaler mouthpiece opening. Then, five particle release maps are generated to represent the initial injection coordinates (X and 𝑌) at the mouthpiece opening. These maps are obtained using a backtracking strategy based on CFPD-FMD simulation results and are classified by particle deposition region, i.e., RML, RLL, LLL, RUL, and LUL.
2. *Step 2: Definition of the Search Space and Spatial Scanning:* A circular search space with a diameter 𝐷_𝑚_ = 20 mm and centered at (0,0) was defined at the mouthpiece opening. Candidate nozzle diameters were constrained between 5 mm and 20 mm based on manufacturing considerations (Cai et al., 2022). Using spatial resolutions of 0.2 mm for nozzle diameter, all candidate locations within the circular search space corresponding to each designated mapping diameter were systematically evaluated. For each candidate region, the number of particles from each lung lobe contained within the mapped region was recorded.
3. *Step 3: Evaluation of Delivery Uniformity Across Lobes:* To quantify the uniformity of particle delivery across 5 lobes, the coefficient of variation (𝐶𝑜𝑉), a widely used statistical metric for dispersion, was employed. 𝐶𝑜𝑉 is a unitless percentage-based measure, and a lower 𝐶𝑜𝑉 indicates a more uniform particle distribution across the five lung lobes. It is expressed as

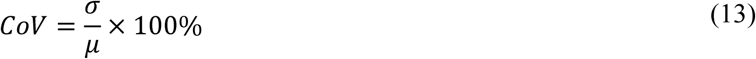

where 𝜎 and 𝜇 are the standard deviation (SD) and the mean value of the particle 𝐷𝐹𝑠 across the five lung lobes.
4. *Step 4: Determination of the Final Optimal Nozzle Configuration:* For each candidate nozzle diameter, the 𝐶𝑜𝑉s of all tested circular regions are compared to identify the optimal TDD nozzle location (i.e., nozzle center coordinates) corresponding to that diameter, as illustrated in **Fig. 5**. Subsequently, the optimal configurations obtained for all candidate diameters ranging from 5 to 20 mm are compared to determine the final TDD nozzle diameter and location. The selection follows two criteria: (i) To minimize 𝐶𝑜𝑉 to achieve the highest uniformity of particle delivery across the five lung lobes, and (ii) to maximize the delivery dose to the TB tree when 𝐶𝑜𝑉 values are comparable. The final TDD configuration corresponds to the location and diameter that yields the smallest 𝐶𝑜𝑉 (see the pink-circled region in **Fig. 5**), and this optimal configuration is subsequently used to generate datasets for ML training and testing.

**Figure 5:**
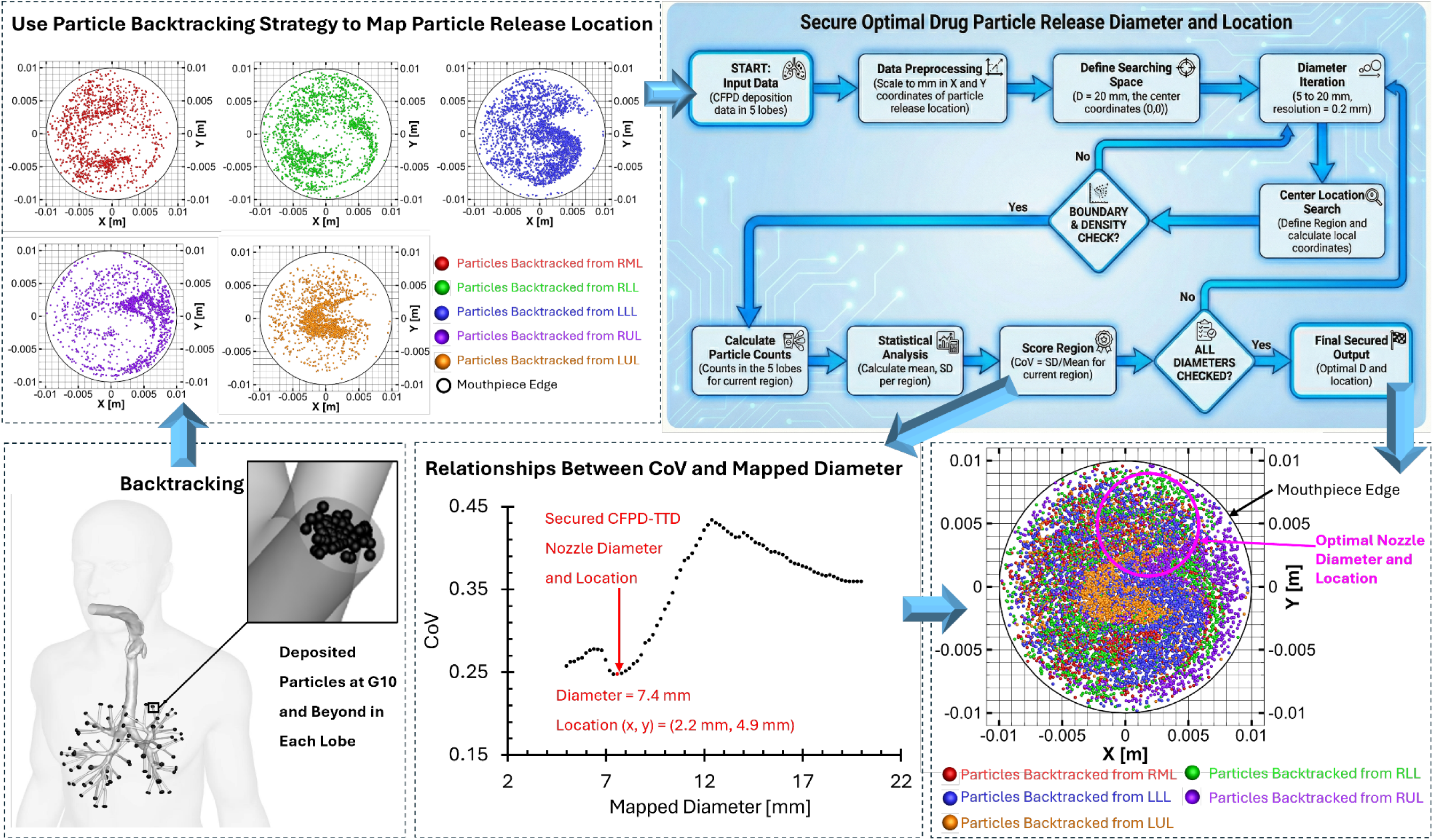
Framework and workflows of securing the optimal CFPD-TDD nozzle location and diameter.

### 2.4 CFPD-Informed ML Model

This study leverages a computational and data-driven framework to develop and explore an ML model that optimizes nozzle parameters for targeted drug delivery to small airways, achieving evenly distributed deposition across five lobes. The ML model serves as a critical component of the inverse design framework. It enables the mapping between patient-specific and drug-specific factors as inputs (i.e., inhalation flow rate 𝑄_𝑚𝑎𝑥_, particle diameter 𝑑_𝑝_, z-coordinate 𝑧_𝑐_ of the nozzle center, and particle release time 𝑡_𝑟_) and optimal nozzle parameters as outputs (i.e., nozzle diameter 𝑑_𝑛_, and nozzle center coordinates 𝑥_𝑐_ and 𝑦_𝑐_). By learning the underlying relationships between the above-mentioned inputs and outputs from the CFPD dataset, it predicts optimal nozzle configurations tailored to personalized breathing patterns and drug properties. The implementation of this framework could provide guidance on selecting the geometric dimensions of the inhaler nozzle and on specific inhaler use conditions to achieve better therapeutic effects. In this study, which 16 ML models were trained to learn inverse mapping from patient- and drug-specific inputs to optimal nozzle design parameters. Details are presented below (also see **Appendix B**).

#### 2.4.1 Data Generation and Preprocessing

108 CFPD simulation results were generated and prepared as the training (80%) and testing (20%) datasets (see **Appendix A**). The data samples highlighted in orange in **Appendix A** are obtained from particle release maps generated from CFPD-FMD simulations, where X-C (i.e., 𝑥_𝑐_), Y-C (i.e., 𝑦_𝑐_) and ND (i.e., 𝑑_𝑛_) are determined using the searching strategy as described in Section 2.3. Each CFPD-FMD simulation varies in key parameters (see **Table 1**). Specifically, the data structure obtained from each CFPD-FMD simulation are shown in **Fig. 1** (also see Section 2.2.2). The optimal parameters, i.e., the TDD nozzle diameter (𝑑_𝑛_) and location (𝑥_𝑐_, 𝑦_𝑐_) ensure maximized delivery efficiency to small airways and uniform delivery across the five lobes (see Section 3.3). All input features (i.e., 𝑄_𝑚𝑎𝑥_, 𝑑_𝑝_, 𝑧_𝑐_, 𝑡_𝑟_) were standardized by being scaled to the standard normal distribution to ensure consistent training performance and efficient gradient descent. Two additional CFPD simulations are used as cross-validation datasets to validate ML models and are provided in Appendix C, with “CFPD-FMD” in the “Models” column.

#### 2.4.2 Model Architecture Selection and Performance Evaluation

To determine the most effective ML approach for optimizing inhaler nozzle parameters in targeted drug delivery, multiple deep learning models were developed and evaluated. These models were designed to predict the optimal 𝑑_𝑛_, 𝑥_𝑐_, and 𝑦_𝑐_ based on patient-specific and drug-specific input parameters (see **Fig. 1** and **Table 1**). Specifically, 16 ML models were implemented and tested. These models are designed to evaluate the effect of model structure and model depth on regression performance. They include 4 standard regression models, 4 regression models with dropout regularization, 4 convolutional neural network (CNN)-based regressors, and 4 transformer-based regressors. The standard regression models vary in the number and width of their hidden layers. Dropout-based regression models introduced stochastic regularization. CNN-based models utilize convolutional layers to capture local interactions among features. Transformer-based regressors investigate the ability of self-attention to model dependencies within the input vector. All models, except Transformer-based models, are trained with ReLU activations and a final linear output layer. The architectures of ML models and their corresponding abbreviations are described as follows.

##### Multilayer Perceptron (MLP)-Based Models

Four multilayer perceptron (MLP) architectures with varying depths and widths were developed to evaluate the influence of model complexity on predictive performance. All models employed ReLU activation functions and a final linear output layer.

- **Baseline Regression Model (RM):** A standard MLP containing three hidden layers with 64, 64, and 32 neurons.
- **Small Regression Model (RM_S):** A shallow network comprising two hidden layers with 32 and 16 neurons, respectively.
- **Medium Regression Model (RM_M):** A deeper architecture with four hidden layers of 128, 128, 64, and 32 neurons.
- **Deep Regression Model (RM_D):** The most complex structure, incorporating five hidden layers with 256, 128, 128, 64, and 32 neurons.

##### Dropout-Enabled MLP Regression Models

To improve generalization and reduce overfitting, dropout layers (*p* = 0.3) were incorporated after each hidden layer. These models retained the same architecture as their non-dropout counterparts.

- **Baseline Regression Model with Dropout (RMD):** Based on RM with dropout.
- **Small Regression Model with Dropout (RMD_S):** Based on RM_S with dropout layers added.
- **Medium Regression Model with Dropout (RMD_M):** Based on RM_M with dropout.
- **Deep Regression Model with Dropout (RMD_D):** Based on RM_D with dropout.

##### Convolutional Neural Network (CNN)–Based Regression Models

Four one-dimensional CNN regressors were constructed to capture localized feature interactions within the input data. The input vector was reshaped into a 1D signal of shape *(Batch_size, 1, 4)*, processed through convolutional layers, and then flattened before being passed through fully connected layers.

- **Baseline CNN Regressor (CNN_B):** A baseline CNN comprising one 1D convolutional layer (16 filters, kernel size = 2) followed by a 32-unit fully connected layer.
- **Small CNN Regressor (CNN_S):** A lightweight model with one 1D convolutional layer (8 filters, kernel size = 2) followed by a 16-unit fully connected layer.
- **Medium CNN Regressor (CNN_M):** A deeper architecture with two 1D convolutional layers (16 and 32 filters, kernel sizes = 2 and 2), followed by a 64-unit fully connected layer.
- **Deep CNN Regressor (CNN_D):** The most complex CNN variant, featuring three 1D convolutional layers (16, 32, and 64 filters, kernel sizes = 2, 2, and 1), dropout (*p* = 0.3), and two fully connected layers (64 and 32 units) before the output layer.

##### Transformer-Based Regression Models

Four transformer-based regressors were developed to explore the capability of self-attention mechanisms to model long-range dependencies among input features. The models differ in embedding dimensionality, number of attention heads, and encoder depth.

- **Baseline Transformer Regressor (TF):** Employs a 64-dimensional embedding layer, two encoder layers with two attention heads, and a 64-unit fully connected layer.
- **Small Transformer Regressor (TF_S)**: Includes a 32-dimensional embedding layer, one encoder layer with a single attention head, and a 32-unit fully connected layer.
- **Medium Transformer Regressor (TF_M):** Features a 64-dimensional embedding layer, two encoder layers with two attention heads, and a dropout rate of 0.1. A LayerNorm layer precedes a 64-unit fully connected layer.
- **Deep Transformer Regressor (TF_D):** The deepest transformer variant, consisting of a 128-dimensional embedding layer, four encoder layers with four attention heads, and dropout (*p* = 0.1). A LayerNorm layer precedes a 128-unit fully connected layer with additional dropout (*p* = 0.2), followed by a final 64-unit output layer.

All ML models listed above were trained using mean squared error (MSE) as the loss function. To evaluate the prediction accuracy and generalization capability of each model, both MSE and mean absolute error (MAE) were calculated on the testing set (see the cases with names in bold in **Appendix A**) and the validation set (see the evaluation metrics on the testing and validation sets in **Appendix B**). Both feature-wise and average MSEs and MAEs are calculated according to equations listed below.

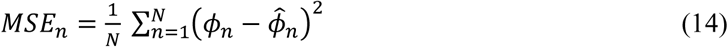

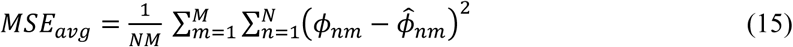

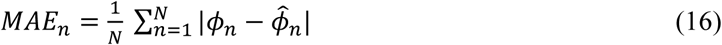

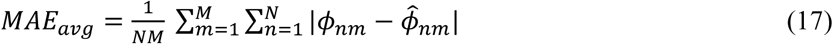

In **Eqs. (14)-(17)**, 𝜙_𝑛_ and 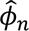 denote the ground-truth CFPD value and the corresponding ML-predicted value for the n-th sample, respectively. The subscript 𝑚 represents the feature index. Accordingly, 𝑦_𝑛𝑚_ and 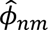 correspond to the true and predicted values of the m-th feature for the n-th sample. 𝑁 is the total number of samples in the datasets, and 𝑀 is the total number of output features.

It is worth noting that the goal of testing multiple architectures was to identify a model that minimizes prediction error, prioritizing the best uniformity of deposition across all five lung lobes, and enhancing deposition fraction beyond G10 compared with conventional full-mouth particle inhalation. 3 ML models with the lowest MSE and a mixed model (i.e., MixModel), which is an ensemble of the 3 models with the lowest feature-wise MSE (see **Appendix C**), in addition to those listed in **Appendix A**. These additional CFPD simulations were conducted to validate the predictive reliability of selected ML models and to assess their real-world capability to guide TDD optimization. Once validated, the selected ML models can be integrated into an innovative smart inhaler system, enabling automatic, patient-specific, and drug-specific adjustments (e.g., optimizing nozzle positioning and drug-release parameters) to achieve precise, uniform aerosol deposition within the small airways, thereby enhancing therapeutic efficacy in pulmonary treatment.

### 2.5 Numerical Setup

#### 2.5.1 CFPD Simulation

Ansys Fluent 2023R1 (Ansys Inc., Canonsburg, PA) was used to predict the transport and deposition of aerosolized drug particles and to generate particle release maps for TDD to the small airways. The numerical simulations used a time step of 0.01 s to ensure stability, and convergence was confirmed when all residuals dropped below 1E-5. The governing equations were solved using a finite-volume method with a second-order upwind momentum scheme, employing the Semi-Implicit Method for Pressure-Linked Equations (SIMPLE) for pressure-velocity coupling in both time and space. The numerical simulations were run on a Dell Precision T7910 workstation equipped with dual Intel® Xeon® Processor E5-2643 v4 processors, providing 24 threads and 128 GB of RAM. Each simulation, which involved a 1-second inhalation, took approximately 11 hours and used 12 threads. In-house user-defined functions (UDFs) and MATLAB scripts were utilized for various purposes, including:

- Generating monodispersed particle injection files,
- Defining the transient sinusoidal inhalation-exhalation waveforms at the mouth opening,
- Customizing the drag force on the particles,
- Specifying the Euler-Lagrange discrete phase model (DPM) time step, and
- Post-processing particle deposition data and generating the particle release maps.

#### 2.5.2 ML Model Training and Testing

The 16 ML models were trained on a location Dell workstation equipped with an AMD Ryzen^TM^ 7950X (16 cores and 32GB of RAM) and an NVIDIA 5070 (6144 CUDA cores and 12GB of video RAM). All ML model training was conducted using the Adam optimizer with a learning rate of 1E-4 and a batch size of 16. The training duration was dynamically controlled based on the composition of the training data. For models trained based on CFPD data listed in **Appendix A**, 15,000 epochs were executed to maximize feature learning. Gradient descent was performed using mini batches with randomized shuffling at each epoch to ensure stable convergence. The CFPD dataset was divided into training and testing sets (i.e., testing dataset) with an 80:20 split, i.e., 86:22 samples. 2 additional CFPD simulations were conducted with distinct feature combinations to generate external validation samples, supplementing the 22 testing cases described in **Appendix A**. Testing samples are marked in bold in **Appendix A**. Model selection is guided by MSE on testing set, with a focus on generalizability to real-world data. During training, the model’s performance on the testing set was used to identify the best-performing epoch based on MSE, which served as the early-stopping criterion. The training times for all ML models are summarized in **Table 2**.

**Table 2:** ML Model training time and index of epoch for achieving the best performance.

| ML Model Name | Training Time [s] | Best Epoch |
| --- | --- | --- |
| RM | 38 | 1323 |
| RM_S | 45 | 3620 |
| RM_M | 49 | 398 |
| RM_D | 60 | 147 |
| RMD | 43 | 1021 |
| RMD_S | 69 | 14885 |
| RMD_M | 69 | 861 |
| RMD_D | 94 | 830 |
| CNN_B | 44 | 1265 |
| CNN_S | 45 | 2723 |
| CNN_M | 51 | 632 |
| CNN_D | 67 | 407 |
| TF | 87 | 20 |
| TF_S | 142 | 72 |
| TF_M | 143 | 43 |
| TF_D | 258 | 78 |

### 2.6 CFPD Model Validation

Comparisons with benchmark experimental data on pulmonary airflow field and particle deposition have validated the CFPD governing model employed in this study. Details of the validations can be found in existing publications (Feng et al., 2016; Feng et al., 2015; Feng et al., 2017; Haghnegahdar et al., 2019).

## 3. Results and Discussion

### 3.1 Pulmonary Airflow Fields

**Figures 6-8** present respiratory airflow patterns at three selected cross-sections (i.e., Planes A, B, and C) in the mouth-to-G10 airway domain, including contours of airflow velocity magnitude 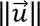, in-plane secondary flow velocity vectors, and turbulence kinetic energy ( 𝑇𝐾𝐸). To provide a detailed visualization of pulmonary airflow dynamics during inhalation with the three inhalation-exhalation waveforms, i.e., 𝑇𝑉= 300, 500, and 750 ml (see **Fig. 4**), three time stations were selected, i.e., t = 0.3 s in **Fig. 6** (i.e., the acceleration phase), 0.5 s in **Fig. 7** (i.e., the peak inhalation), and 0.7 s in **Fig. 8** (i.e., the deceleration phase).

**Figure 6:**
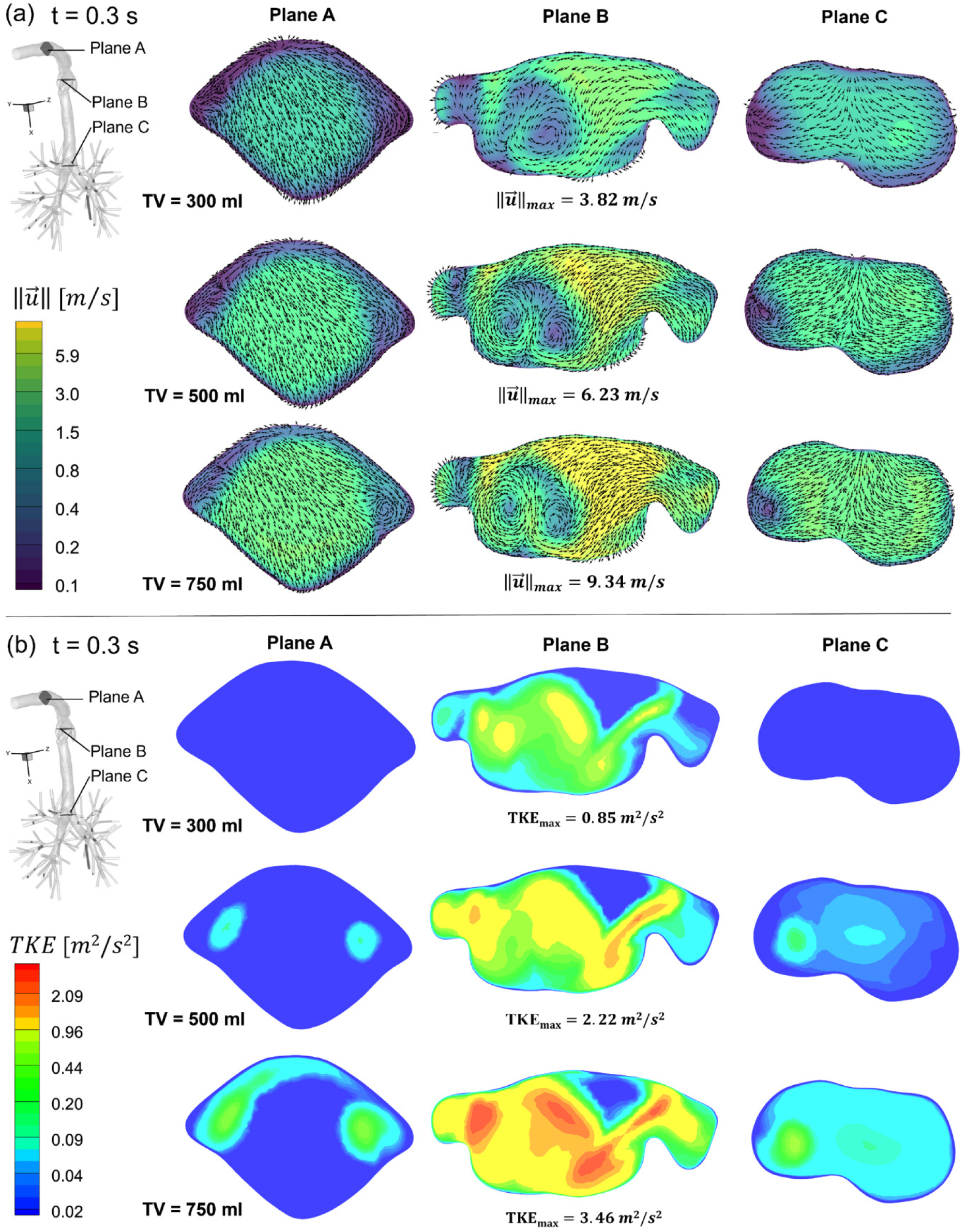
Airflow velocity magnitude contours, in-plane velocity vector distributions and TKE distributions at multiple cross sections at time t= 0.3 s under three investigated TVs: (a) Airflow velocity magnitude contours and in-plane velocity vector distributions, and (b) TKE distributions.

**Figure 7:**
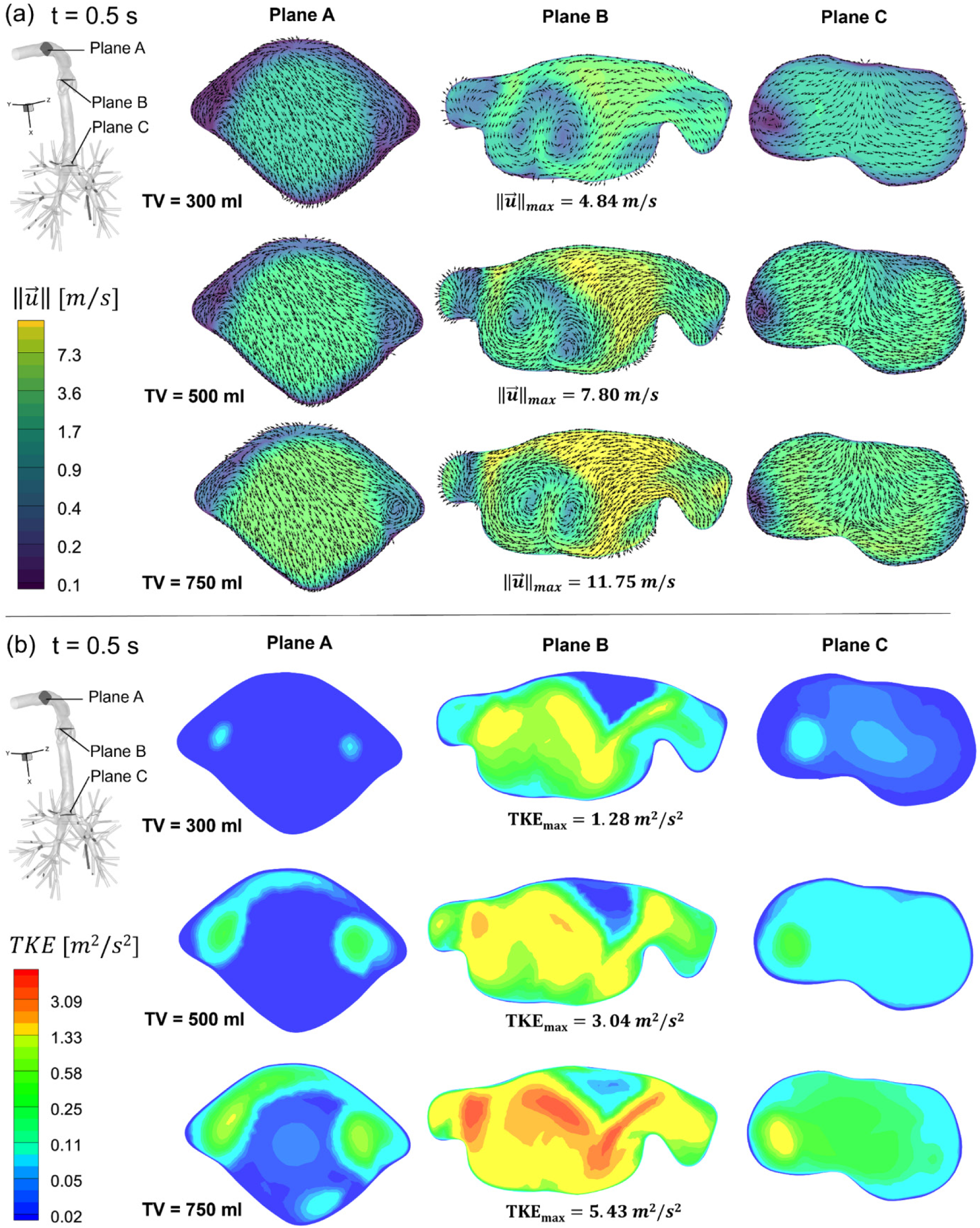
Airflow velocity magnitude contours, in-plane velocity vector distributions and TKE distributions at multiple cross sections at time t= 0.5 s under three investigated TVs: (a) Airflow velocity magnitude contours and in-plane velocity vector distributions, and (b) TKE distributions.

**Figure 8:**
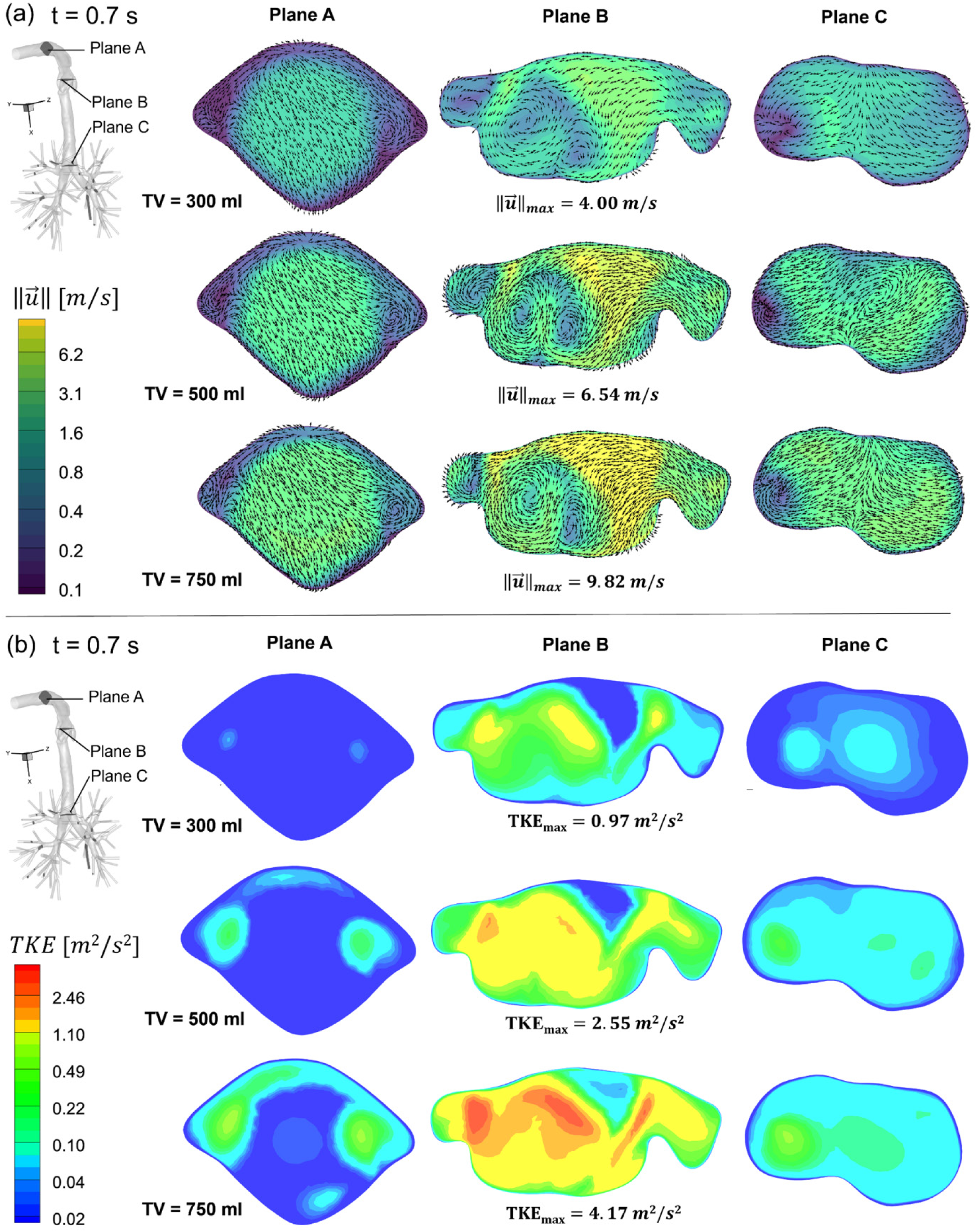
Airflow velocity magnitude contours, in-plane velocity vector distributions and TKE distributions at multiple cross sections at time t= 0.7 s under three investigated TVs: (a) Airflow velocity magnitude contours and in-plane velocity vector distributions, and (b) TKE distributions.

It can be observed that the complex anatomy of the upper airway significantly perturbs inspiratory flow, which is highly dependent on breathing patterns (see **Figs. 6 (a)**, **7 (a), & 8 (a)**). Specifically, in the anterior oral cavity (see Plane A), the velocity distribution is relatively more even than in downstream regions (i.e., Planes B and C). This behavior is attributed to the extended tube connected to the oral cavity and the relatively small variations in cross-sectional shape and area. Comparisons of in-plane velocity vector distributions indicate that secondary flow is relatively weaker at lower 𝑇𝑉 (i.e., 300 ml), while it becomes stronger with paired counter-rotating vortices at higher volumes (𝑇𝑉 = 750 ml) due to the stronger mouth inlet jet and the curvature from the oral cavity to the oropharynx. In the glottis region (i.e., Plane B), the contraction in lumen at the vocal cords generates a laryngeal jet, exhibiting the highest velocity magnitude 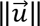 across all time points, peaking at 𝑡 = 0.5 s with a maximum 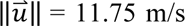 with 𝑇𝑉 = 750 ml. The sudden expansion downstream of the glottis creates intense shear layers and complex, strong secondary flows, characterized by massive recirculation zones and vortices laterally adjacent to the main jet core in the trachea. Near the G0-G1 bifurcation (i.e., Plane C), the laryngeal jet extends into the trachea. Still, it begins to break down, with a highly asymmetric flow pattern pushed toward specific walls by upstream curvature and the impending geometric split. Secondary flows here resemble classic Dean’s flow, driven by the pressure gradients of the curving airway and the flow splitting into the main bronchi.

Furthermore, it can be found that TKE scales non-linearly with both inhalation process and TV (see **Figs. 6 (b)**, **7 (b), & 8 (b)**), and turbulence is maximized at the peak inspiration instant (i.e., t = 0.5 s) under the highest inhalation flow rate (i.e., 𝑇𝑉 = 750 ml), reaching 5.43 m^2^/s^2^. Similarly to the velocity distributions, TKE remains nearly equal to zero in the anterior oral cavity (i.e., Plane A), indicating that the flow remains largely laminar or transitional, even at high flow rates. Then, TKE increases significantly in the glottis region (i.e., Plane B), where the intense shear layers between the high-speed laryngeal jet core and the recirculating air near the tracheal walls generate stronger turbulence. Near the G0-to-G1 bifurcation (i.e., Plane C), TKE remains higher than Plane A, but is more evenly distributed than the glottis region, indicating turbulent dissipation as the airflow travels down the trachea, though the flow remains highly disturbed as it enters G1.

Based on the comparisons of flow patterns shown in **Figs. 6-8**, the strong secondary flow and high TKE near Plane B can potentially lead to particle deposition induced by turbulence dispersion from the posterior oral cavity to the oropharynx. The laryngeal jet (i.e., Plane B and downstream) can induce particle deposition induced by inertial impaction to the G0-to-G1 bifurcation. The laryngeal jet induced strong secondary flow and turbulence in the trachea, which can result in high localized particle deposition on the posterior or lateral walls of the upper trachea immediately downstream, which is aligned with the previous findings (Longest et al., 2019; Yi & Feng, 2026). More detailed airflow influences on inhaled particles transport and deposition can be found in Section 3.2.

### 3.2 Localized Particle Deposition Patterns

To investigate the delivery efficiency of inhaled particles to small airways and the evenness of the particle delivery across five lobes (i.e., LUL, LLL, RUL, RML, and RLL), both local particle deposition patterns from the mouth to the smaller airways and regional DFs in the five lobes were predicted and visualized (see **Figs. 9-11**). Specifically, this study selected four different particle diameters of 𝑑_𝑝_= 0.5, 1, 2, and 5 μm under three different 𝑇𝑉s (i.e., 300, 500, and 750 ml), corresponding to three inhalation peak flow rates 𝑄_𝑚𝑎𝑥_ (i.e., 28.27, 47.12, and 70.69 L/min), respectively. **Figure 9** illustrates the influence of particle size on particle delivery under the three 𝑇𝑉s. For demonstration purposes, the particle release time and release location were fixed at 𝑡_𝑟_ = 0 s and 𝑧_𝑐_ = 1 mm. **Figure 10** visualizes the impact of particle release location ( 𝑧_𝑐_) on particle delivery under three different inhalation-exhalation waveforms, using 1 μm particles as an example. **Figure 11** further shows particle release time (𝑡_𝑟_) effect on particle delivery under three different breathing patterns for 1 μm particles released at 𝑧_𝑐_ = 1 mm.

**Figure 9:**
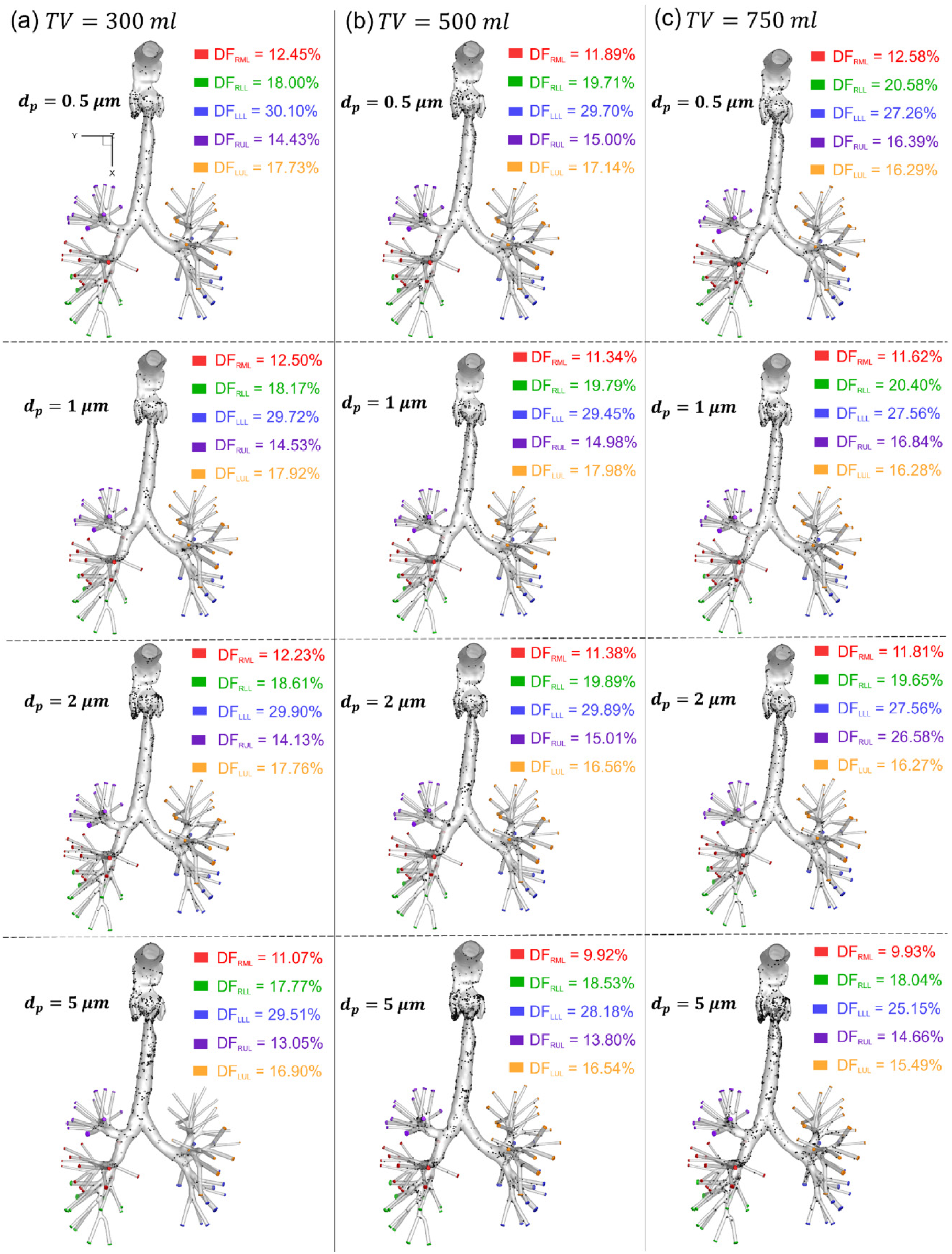
Regional deposition fraction (DF) in each lobe and local deposition patterns of inhaled drug particles at release time 𝑡_𝑟_= 0 s and release location 𝑧_𝑐_= 1 mm with three different tidal volumes (TV): (a) TV = 300 ml, (b) TV =500 ml, and (c) TV = 750 ml.

**Figure 10:**
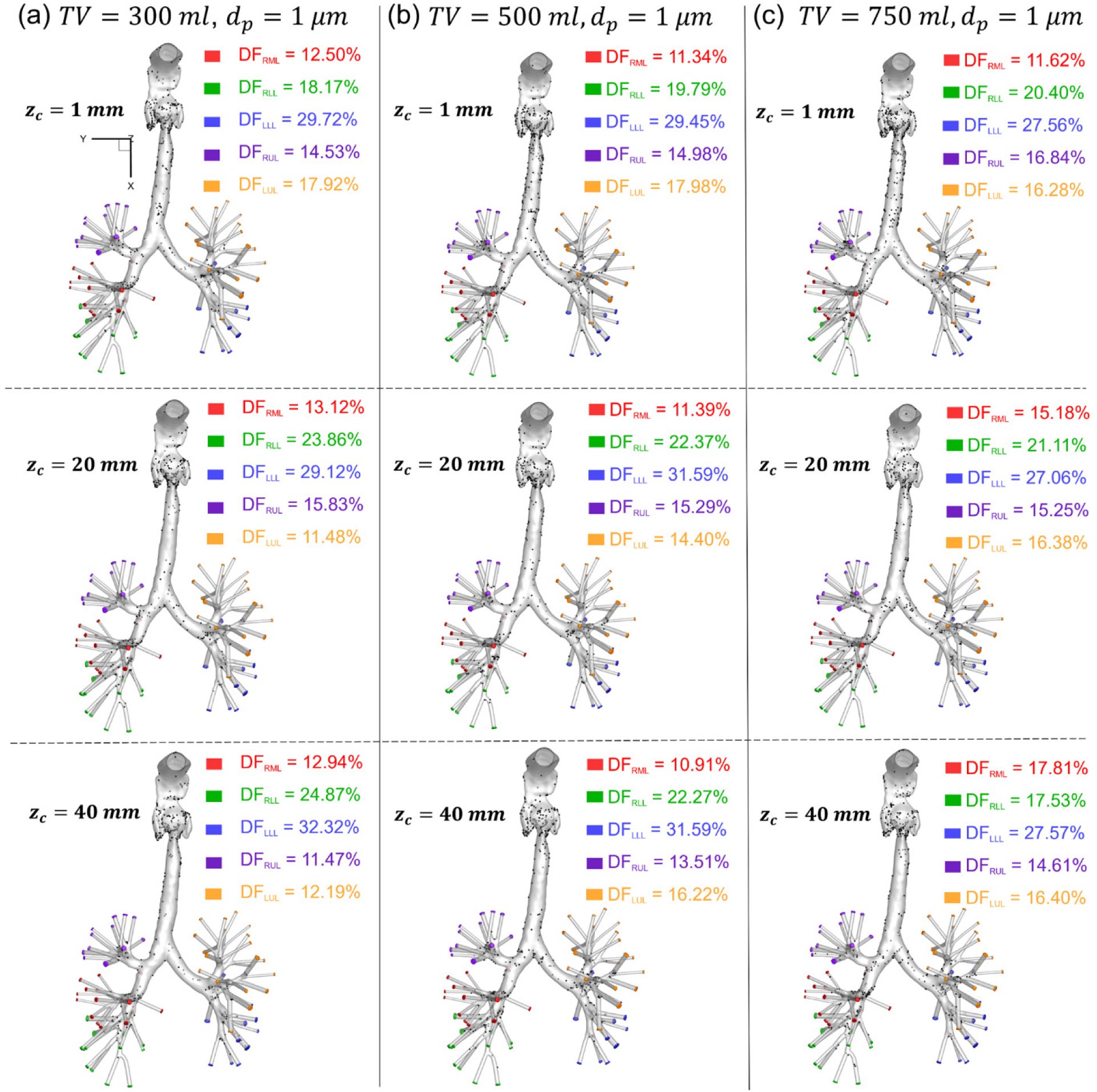
Regional deposition fraction (DF) in each lobe and local deposition patterns of inhaled drug particles at release time 𝑡_𝑟_= 0 s and with drug particle size 𝑑_𝑝_ = 1 µm at three different release locations (𝑧_𝑐_= 1 mm, 20 mm, and 40 mm): (a) TV=300 ml, (b) TV=500 ml, and (c) TV=750 ml.

**Figure 11:**
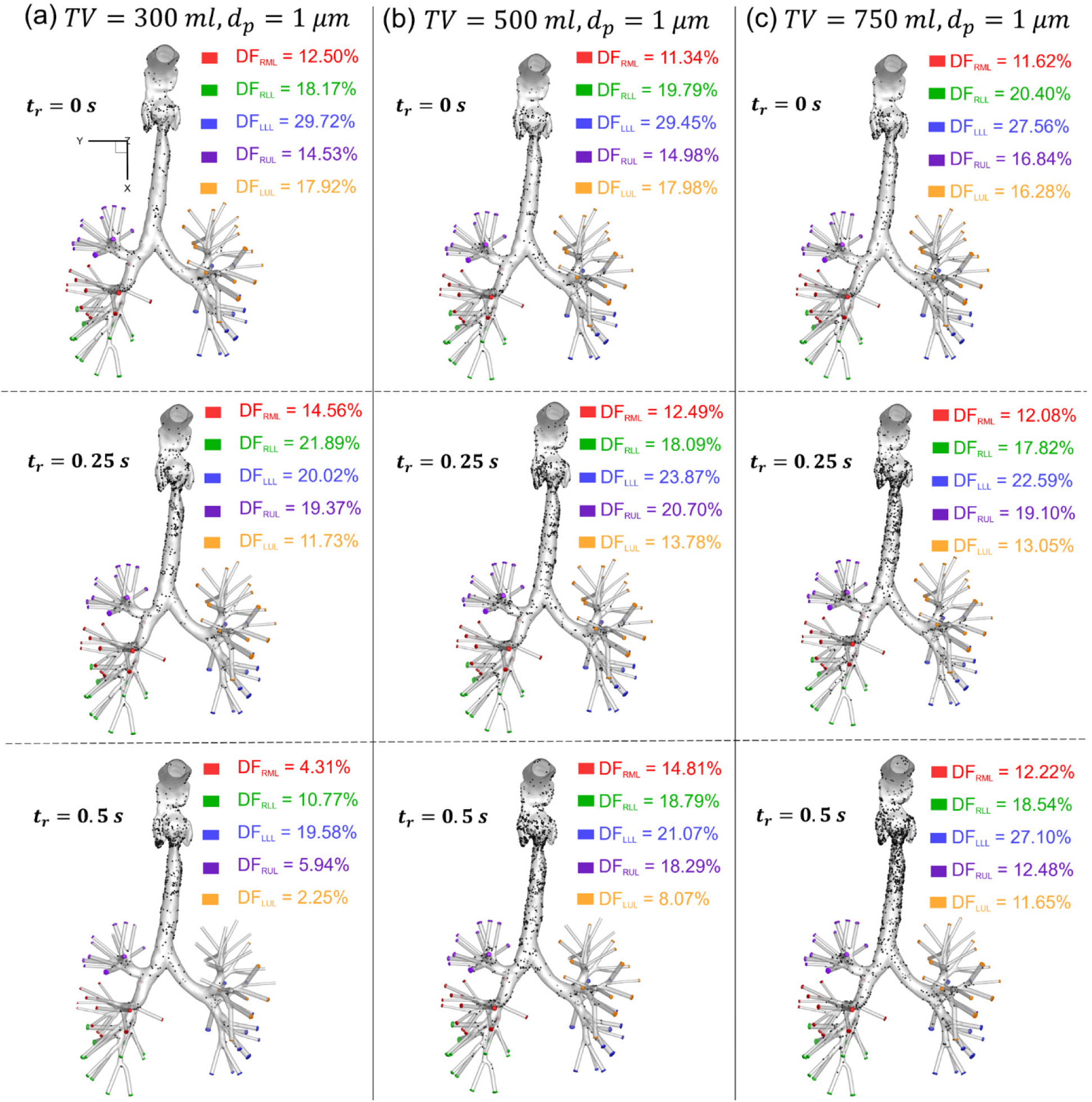
Regional deposition fraction (DF) in each lobe and local deposition patterns of inhaled drug particles (𝑑_𝑝_ = 1 µm) released at 𝑧_𝑐_= 1 mm with three different release times (𝑡_𝑟_= 0 s, 0.25 s and 0.5 s): (a) TV=300 ml, (b) TV=500 ml, and (c) TV=750 ml.

As shown in **Fig. 9**, regardless of particle size or inspiratory flow rate, the distribution of particles reaching the lobar outlets is highly asymmetric. LLL consistently receives the highest DF of particles, ranging from approximately 25% to 30%, followed by RLL at approximately 18% to 20%. This anatomical preferential deposition in the lower lobes is primarily driven by the relatively straight and gravity aligned trajectories of the main bronchi as they transport into the peripheral lung. The airflow pathways to the upper lobes (i.e., LUL and RUL) require sharper directional changes, reducing convective particle delivery to those regions.

With respect to particle size (𝑑_𝑝_), the results demonstrate a clear shift in transport mechanisms as particle size increases from 0.5 µm to 5 µm. Smaller particles (0.5 to 2 µm) exhibit lower inertia and Stokes numbers, allowing them to closely follow airflow streamlines. As a result, the lobar DFs remain relatively stable across all particle sizes within this range, indicating minimal deposition in the upper airway. However, when 𝑑_𝑝_ = 5 µm, a noticeable increase in deposited particles can be observed in the upper airway, i.e., from the oropharynx to the upper trachea (see **Fig. 9**). The increased deposition in the upper airway is due to stronger inertial impaction and gravitational sedimentation of larger particles. Therefore, more particles deposit upstream, and fewer reach the TB tree, resulting in a noticeable decrease in lobar DFs (e.g., DF in LLL decreases to 25.15% at 𝑇𝑉 = 750 ml).

Additionally, it can be found that increasing 𝑇𝑉 directly enhances inertial impaction and turbulence dispersion effects, as observed in **Figs. 6-8**, thereby leading to higher DF in the upper airway. With the lowest 𝑇𝑉 = 300 ml, even 5 µm particles can navigate the complex extrathoracic and upper tracheobronchial airway with relatively low impaction losses, maintaining high DFs to the deep lung. In contrast, with the highest 𝑇𝑉 (i.e., 750 ml), lower total DFs are observed. Especially, the highest *TV* (i.e., 750 ml) combined with the largest particles (i.e., 𝑑_𝑝_=5 µm), results in the lowest overall delivery to small airways.

Using 𝑑_𝑝_ = 1 µm and 𝑡_𝑟_ = 0 s as an example, **Fig. 10** highlights how particle release location (i.e., axial Z coordinate (𝑧_𝑐_)) impacts particle deposition under three 𝑇𝑉s. It needs to be noted that the ultimate lobar destination of inhaled 1 µm particles depends critically on the airflow streamline they originate from, since these particles have extremely low Stokes numbers and act like flow tracers. Releasing particles at 𝑧_𝑐_ = 1 mm forces particles to traverse the entire oral cavity, where the minor secondary flows and geometric expansions in the mouth perturb the particle cloud before it reaches the throat. While releasing particles at 𝑧_𝑐_ = 40 mm can bypass early oral cavity perturbations, resulting in a much more localized, concentrated particle inlet profile entering the laryngeal jet. This streamline dependence is highly visible at the lowest inhalation rate, i.e., 𝑇𝑉= 300 ml, and shifting the release from z sub c = 1 mm to z sub c = 40 mm significantly shifts DFs in the lobar lobe outlets at G10. For example, as shown in **Fig. 10 (a)**, delivery to the RLL outlets jumps from 18.17% to 24.87%, while delivery to the LUL drops from 17.92% to 12.19%. In addition, increasing 𝑇𝑉 can effectively eliminate this spatial dependence on 𝑧_𝑐_, which is more pronounced at lower 𝑇𝑉 with corresponding lower inhalation flow rates. As shown in **Figs. 6-8**, a higher 𝑇𝑉 (i.e., 750 ml) induces higher TKE with stronger secondary flows immediately downstream of the glottis. Stronger turbulence mixing effects disperse the particles rapidly, regardless of whether the release location at the mouthpiece opening (𝑧_𝑐_ = 1 mm) or near the throat (𝑧_𝑐_ = 40 mm). Consequently, as shown in **Fig. 10 (c)**, at 𝑇𝑉 = 750 ml, the lobar DFs remain relatively static across all lobes (especially the left lung), where LLL stabilizes around 27%, LUL stays around 16%, RUL, RML, RLL remain in the range of 14.6%-15.8%, 11.6%-17.8%, 17.5%-21.1%, respectively. The variation with 𝑧_𝑐_ at 𝑇𝑉 = 750 ml is much less than at 𝑇𝑉 = 300 ml.

**Figure 11** further illustrates the influence of particle release time (𝑡_𝑟_) on regional DFs under the three 𝑇𝑉s using 𝑑_𝑝_ = 1 µm and 𝑧_𝑐_ = 1 mm as a representative case. In general, particles released at 𝑡_𝑟_ = 0 s benefit from the entire 1-second inhalation duration (see **Fig. 4**) to transport into the bronchial tree. Delaying particle release significantly alters particle transport dynamics. Especially at the lowest tidal volume (i.e., 𝑇𝑉 = 300 ml), the delayed-release effect is evident (see **Fig. 11(a)**). Indeed, when particles are released at 𝑡_𝑟_= 0.5 s, lobar DFs decrease compared with 𝑡_𝑟_= 0 s. For example, it can be observed from **Fig. 11 (a)** that the DF in LUL drops from 17.92% at 𝑡_𝑟_= 0 s to only 2.25% at 𝑡_𝑟_= 0.5 s. Under such a low inhalation flow rate associated with 𝑇𝑉 = 300 ml, particles injected at 𝑡_𝑟_ = 0.5 s through inhalation experience insufficient momentum and time for convection to reach peripheral airways within the remaining 0.5 s during inhalation. As a result, instead of depositing in the peripheral lung, a large portion of the particle bolus remains suspended in the upper airway and central lung when the 1-second inhalation ceases. Indeed, using Case 007 and Case 008 as examples (see **Appendix A**) of which 𝑡_𝑟_ = 0.5 s and 𝑇𝑉 = 300 ml, 51.25% (Case 007) and 45.85% (Case 008) of released particles remain suspended after the inhalation. In practical inhalation therapy, these suspended particles are likely to be either exhaled during the subsequent exhalation phase or deposited in proximal airways rather than reaching the intended small-airway targets. Consequently, delayed particle release during the inhalation, combined with mild inhalation (e.g., 𝑇𝑉 = 300 ml) is not favorable for efficient drug delivery to the peripheral lung. In contrast, under the highest tidal volume (𝑇𝑉 = 750 ml) (see **Fig. 11 (c)**), the strong laryngeal jet with peak velocities exceeding 11 m/s at the peak inhalation provides sufficient convective momentum to carry particles to the peripheral lung, even with 𝑡_𝑟_ = 0.5 s. Nevertheless, as inhalation progresses from 𝑡_𝑟_ = 0 s to 𝑡_𝑟_ = 0.5 s, the underlying flow field evolves into a highly turbulent, jet-dominated regime, altering the streamline topology that governs particle transport. For example, at *TV* = 500 ml (see **Fig. 11 (b)**, the DF in LLL decreases from 29.45% at 𝑡_𝑟_ = 0s to 21.07% at 𝑡_𝑟_ = 0.5 s, while the DF in RUL increases correspondingly as turbulent mixing redistributes particles across different airflow pathways. These results suggest that changes in particle release timing do not produce a monotonic trend in DF variations among the lobes. Instead, particle delivery is redistributed among lobar outlets depending on the evolving flow structures during the inhalation phase.

The analyses above demonstrate that particle size, release location, and release timing can significantly influence particle transport and deposition in the human respiratory system. However, these results also indicate that particle delivery across the five lobes remains inherently asymmetric under conventional CFPD-FMD inhalation conditions (i.e., full-mouth particle inhalation). Since the ultimate objective of this study is to develop a CFPD-ML-driven TDD strategy capable of achieving more uniform drug delivery among the five lobes, it is necessary to further quantify how each parameter individually affects delivery uniformity. Therefore, a one-factor analysis was conducted by varying tidal volume (𝑇𝑉), particle diameter (𝑑_𝑝_), release location (𝑧_𝑐_), and release time (𝑡_𝑟_) independently, while evaluating the resulting uniformity of lobar deposition fractions (DFs) using the coefficient of variation (𝐶𝑜𝑉). The statistical indicators of 𝐶𝑜𝑉, including the mean, maximum, median, and standard deviation (SD), are summarized in **Table 3**. When each parameter is examined individually, the results suggest that 𝑇𝑉 = 750 ml, 𝑑_𝑝_ = 2.0 µm, 𝑧_𝑐_ = 20 mm, and 𝑡_𝑟_ = 0.25 s provide the most favorable conditions for improving delivery uniformity. Nevertheless, these values should be interpreted only as reference indicators. In practice, achieving uniform drug delivery across the five lobes results from the combined interaction of multiple parameters, rather than from any single parameter acting alone.

**Table 3.**
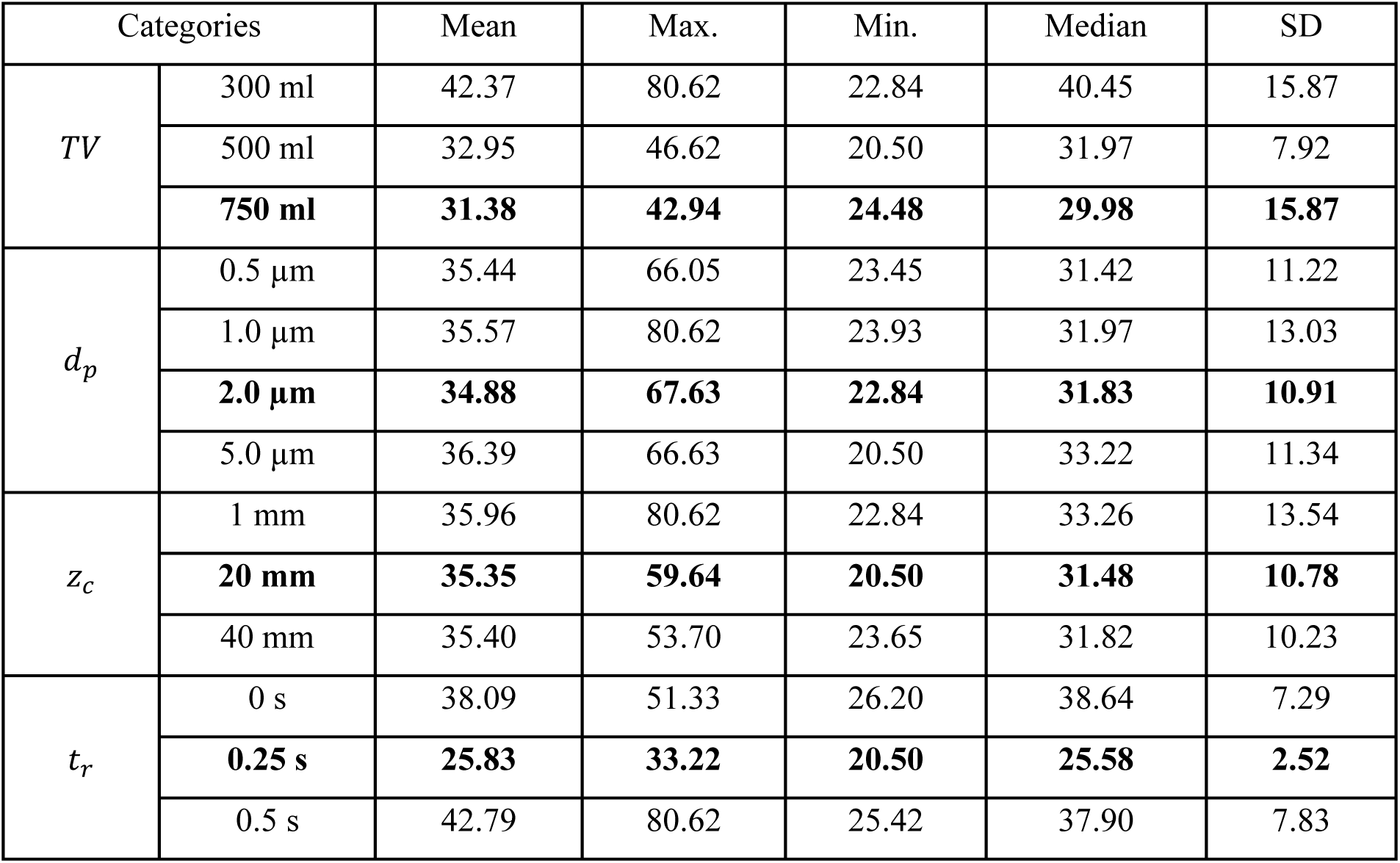
CFPD-FMD results from universal statistical analysis for all parameters examined in 𝐶𝑜𝑉 [%].

| Categories |  | Mean | Max. | Min. | Median | SD |
| --- | --- | --- | --- | --- | --- | --- |
| $TV$ | 300 ml | 42.37 | 80.62 | 22.84 | 40.45 | 15.87 |
|  | 500 ml | 32.95 | 46.62 | 20.50 | 31.97 | 7.92 |
|  | **750 ml** | **31.38** | **42.94** | **24.48** | **29.98** | **15.87** |
| $d_p$ | 0.5 $\mu\text{m}$ | 35.44 | 66.05 | 23.45 | 31.42 | 11.22 |
| | 1.0 $\mu\text{m}$ | 35.57 | 80.62 | 23.93 | 31.97 | 13.03 |
|  | **2.0 $\mu\text{m}$** | **34.88** | **67.63** | **22.84** | **31.83** | **10.91** |
| | 5.0 $\mu\text{m}$ | 36.39 | 66.63 | 20.50 | 33.22 | 11.34 |
| $z_c$ | 1 mm | 35.96 | 80.62 | 22.84 | 33.26 | 13.54 |
|  | **20 mm** | **35.35** | **59.64** | **20.50** | **31.48** | **10.78** |
|  | 40 mm | 35.40 | 53.70 | 23.65 | 31.82 | 10.23 |
| $t_r$ | 0 s | 38.09 | 51.33 | 26.20 | 38.64 | 7.29 |
|  | **0.25 s** | **25.83** | **33.22** | **20.50** | **25.58** | **2.52** |
|  | 0.5 s | 42.79 | 80.62 | 25.42 | 37.90 | 7.83 |

### 3.3 Particle Release Maps and Optimal Nozzle Diameter and Locations

To investigate the effect of tidal volume (𝑇𝑉) and particle sizes (𝑑_𝑝_) on the optimal nozzle diameter (𝑑_𝑛_) for achieving uniform TDD to the small airways, 12 representative particle release maps are visualized in **Fig. 12** as examples, with multiple 𝑇𝑉s (i.e., 300 ml, 500 ml, and 750 ml) and particle sizes (i.e., 𝑑_𝑝_=0.5, 1, 2, and 5 μm). The data in **Fig. 12** were extracted from Cases 001, 010, 019, 028, 037, 046, 055, 064, 073, 082, 091, 100, which can be accessed in **Appendix A**. It is worth mentioning that higher 𝑇𝑉 indicates a higher average inhalation flow rate and peak flow rate 𝑄_𝑚𝑎𝑥_.

**Figure 12:**
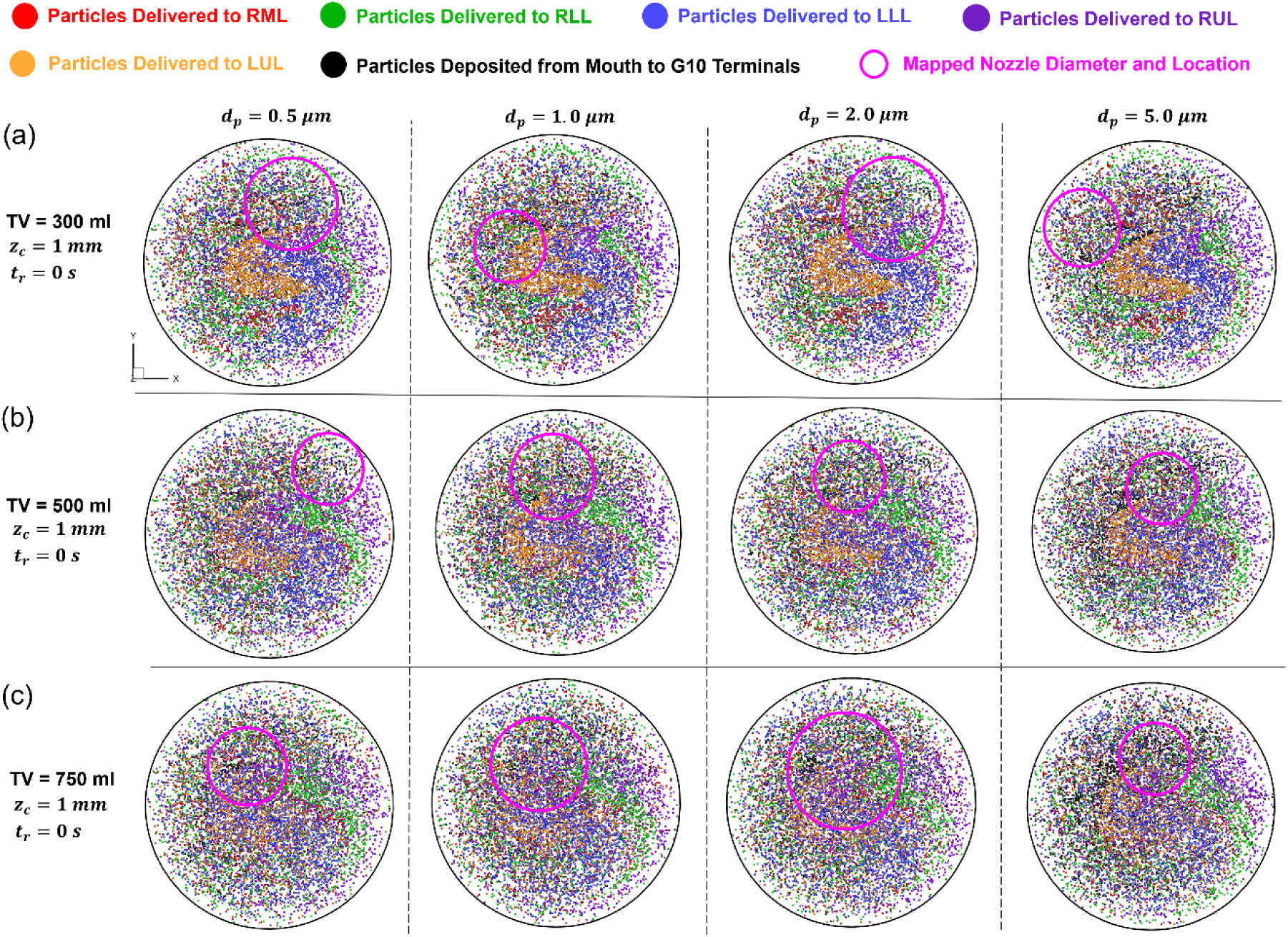
Particle release maps for four different particle sizes (𝑑_𝑝_=0.5, 1, 2, and 5 μm) with constant particle release position 𝑧_𝑐_= 1 mm and particle release time 𝑡_𝑟_ = 0 s under three different tidal volumes: (a) TV = 300 ml, (b) TV = 500 ml, and (c) TV = 750 ml.

Based on CFPD-FMD simulation results representing conventional inhalation therapy, the particle release maps in **Fig. 12** were generated using the methodology presented in **Fig. 5** and Section 2.3. In these maps, the particles are at their release locations and color-coded by their deposition sites, i.e., one of the five lobes. Particles that are deposited somewhere other than the five lobes are shown in black. Connecting particle release positions and their deposition locations, particle release maps can help determine the available regions for particle release to effectively deliver medication to targeted areas, such as small airways in corresponding lobes. Using the mapping method developed in Section 2.3, the optimal nozzle position and diameter can be determined by selecting the area that maximizes the coverage of the five-colored regions. The optimal nozzle parameter selection is based on two principles: (1) the nozzle diameter 𝑑_𝑛_ is from 5 to 20 mm, and (2) the particle numbers with different colors in the circle should be as even as possible. **Figures 13 (a) – 13 (c)** show the relationship between the determined nozzle diameter and the corresponding evaluation rubric (i.e., *CoV* of numbers of particles in different colors). TDD inhaler diameter and location were obtained by choosing the circle with the smallest 𝐶𝑜𝑉, representing the optimal uniformity of DFs in the five lobes. The solid-pink circles in the release maps shown in **Fig. 12** are the nozzles with the optimal nozzle diameter 𝑑_𝑛_ and nozzle center location (𝑥_𝑐_, 𝑦_𝑐_, 𝑧_𝑐_) determined based on CFPD simulations. Furthermore, another 12 representative particle release maps are visualized in **Fig. 14** to demonstrate the impacts of particle release location Z coordinate (𝑧_𝑐_) and particle size (𝑑_𝑝_) on the optimal nozzle diameter 𝑑_𝑛_ and nozzle center location (𝑥_𝑐_, 𝑦_𝑐_) at *TV* = 300 ml. The data in **Fig. 14** were extracted from Cases 001, 002, 003, 010, 011, 012, 019, 020, 021, 028, 029, and 030, which can be accessed in **Appendix A**.

**Figure 13:**
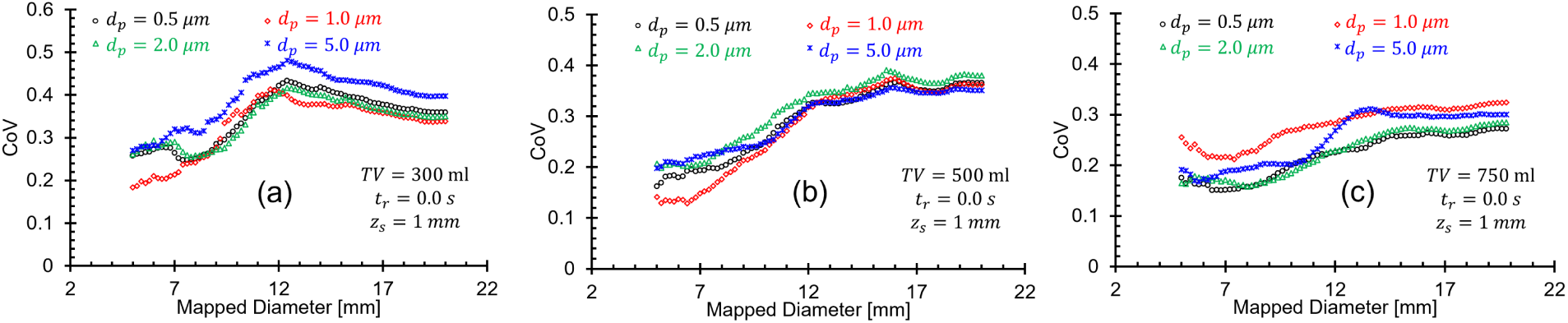
The relationship between the secured TDD nozzle diameter and the corresponding CoV (evaluating uniformity of DFs at five regional lobe outlets) under various settings: (a) TV = 300 ml, 𝑡_𝑟_= 0 s, 𝑧_𝑐_= 1 mm, (b) TV = 500 ml, 𝑡_𝑟_= 0 s, 𝑧_𝑐_= 1 mm, and (c) TV = 750 ml, 𝑡_𝑟_= 0 s, 𝑧_𝑐_= 1 mm.

**Figure 14:**
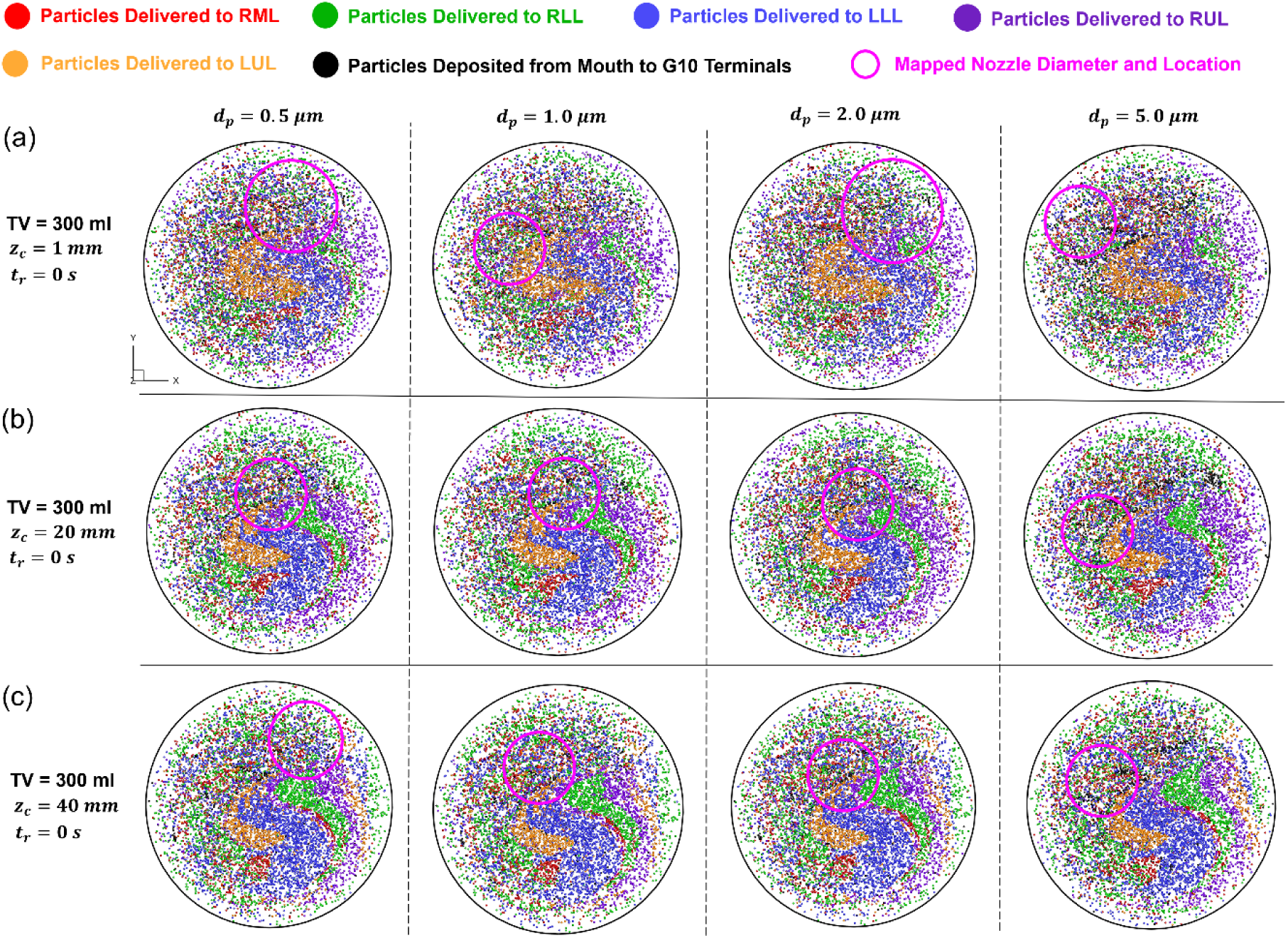
Particle release maps for four different particle sizes (𝑑_𝑝_=0.5, 1, 2, and 5 μm) with constant TV = 300 ml and 𝑡_𝑟_= 0 s for three different particle release locations at Z coordinate: (a) 𝑧_𝑐_= 1 mm, (b) 𝑧_𝑐_= 20 mm, and (c) 𝑧_𝑐_= 40 mm.

𝐶𝑜𝑉s of numbers of particles in different colors vs. the tested mapped nozzle trial diameters are presented in **Fig. 15**. Additionally, to investigate how the particle release time (𝑡_𝑟_) influences 𝑑_𝑛_ and (𝑥_𝑐_, 𝑦_𝑐_), another 12 particle release maps and the corresponding 𝐶𝑜𝑉s of numbers of particles in different colors vs. the tested mapped nozzle trial diameters under TV = 300 ml and 𝑧_𝑐_ = 1 𝑚𝑚 are illustrated in **Figs. 16 & 17**, respectively. The data in **Fig. 16** were extracted from Cases 001, 004, 007, 010, 013, 016, 019, 022, 025, 028, 031, and 034 which can be accessed in **Appendix A**. To summarize the effects of the parameters mentioned above on 𝑑_𝑛_, **Figs. 18 (a)-(d)** are generated to accompany **Figs. 12, 14, and 16**.

**Figure 15:**
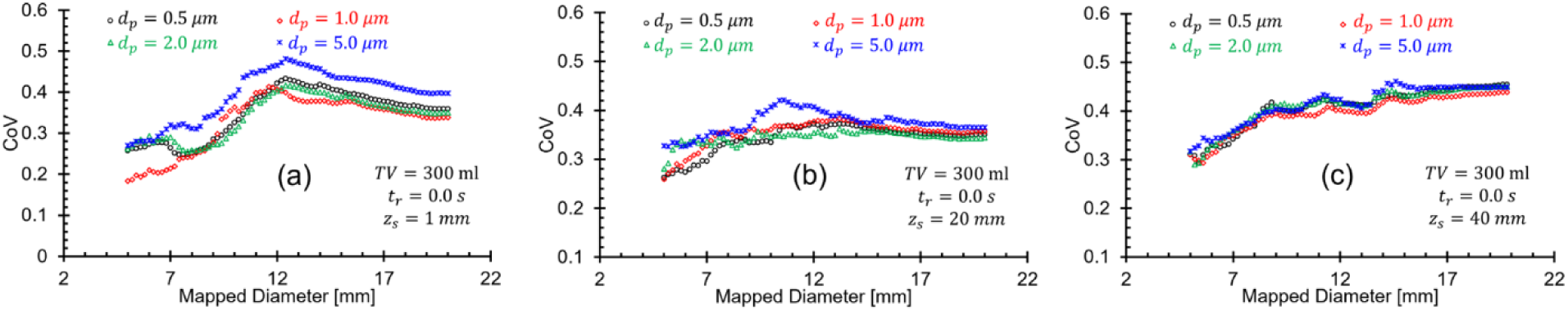
The relationship between the secured TDD nozzle diameter and the corresponding CoV (evaluating uniformity of DFs at five regional lobe outlets) under various settings: (a) TV = 300 ml, 𝑡_𝑟_= 0 s, 𝑧_𝑐_= 1 mm, (b) TV = 300 ml, 𝑡_𝑟_= 0 s, 𝑧_𝑐_= 20 mm, and (c) TV = 300 ml, 𝑡_𝑟_= 0 s, 𝑧_𝑐_= 40 mm.

**Figure 16:**
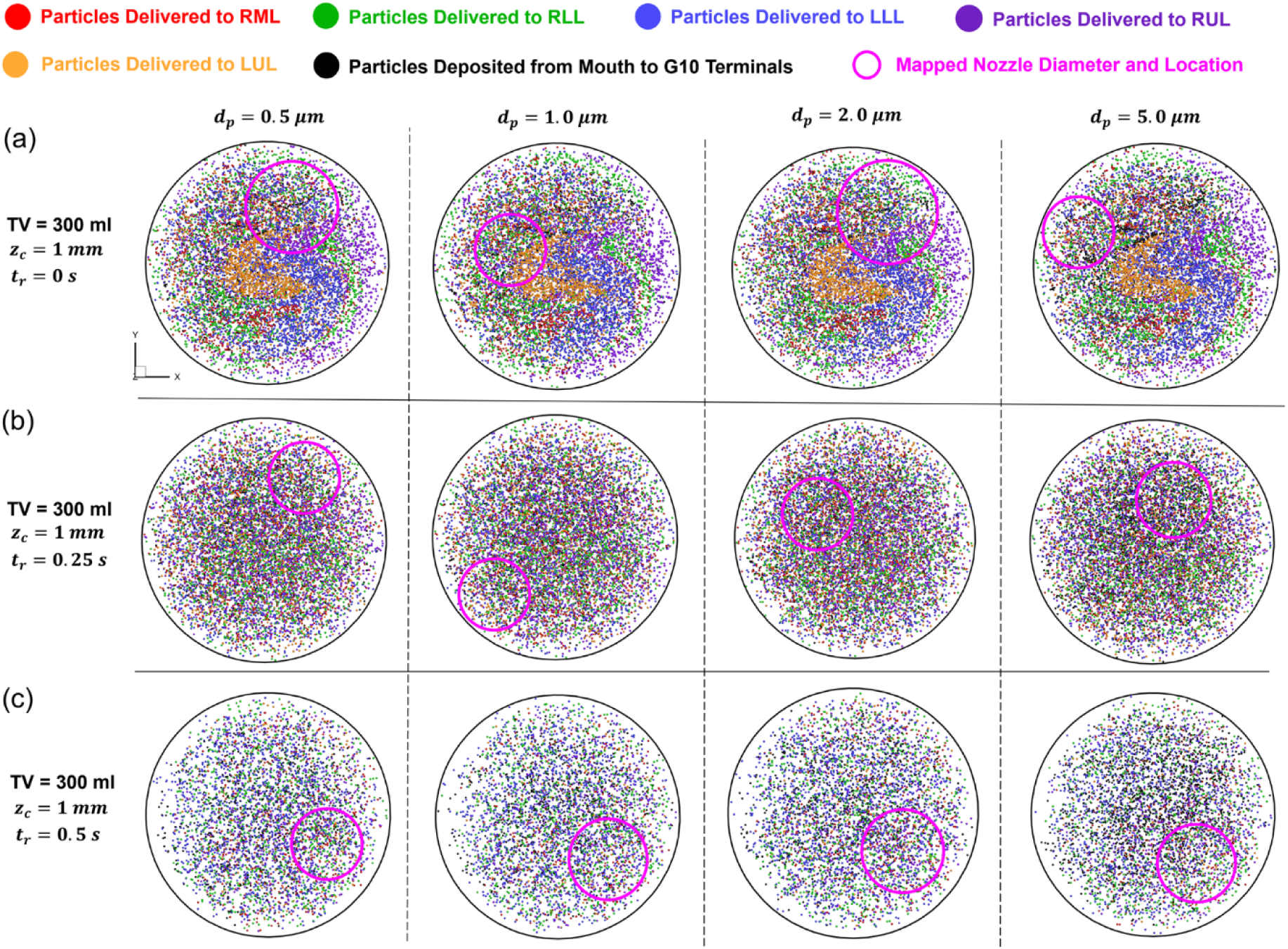
Particle release maps for four different particle sizes (𝑑_𝑝_=0.5, 1, 2, and 5 μm) with constant TV = 300 ml and 𝑧_𝑐_= 1 mm for three different particle release time instants: (a) 𝑡_𝑟_= 0 s, (b) 𝑡_𝑟_= 0.25 s, and (c) 𝑡_𝑟_= 0.5 s.

**Figure 17:**
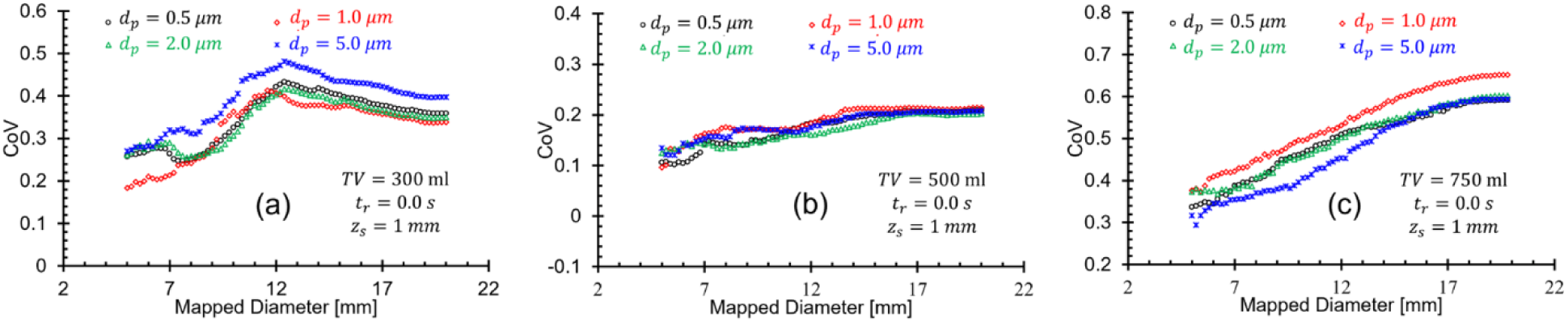
The relationship between the secured TDD nozzle diameter and the corresponding CoV (evaluating uniformity of DFs at five regional lobe outlets) under various settings: (a) TV = 300 ml, 𝑧_𝑐_= 1 mm, 𝑡_𝑟_= 0 s, (b) TV = 300 ml, 𝑧_𝑐_= 1 mm, 𝑡_𝑟_= 0.25 s,, and (c) TV = 300 ml, 𝑧_𝑐_= 1 mm, 𝑡_𝑟_= 0.5 s.

**Figure 18:**
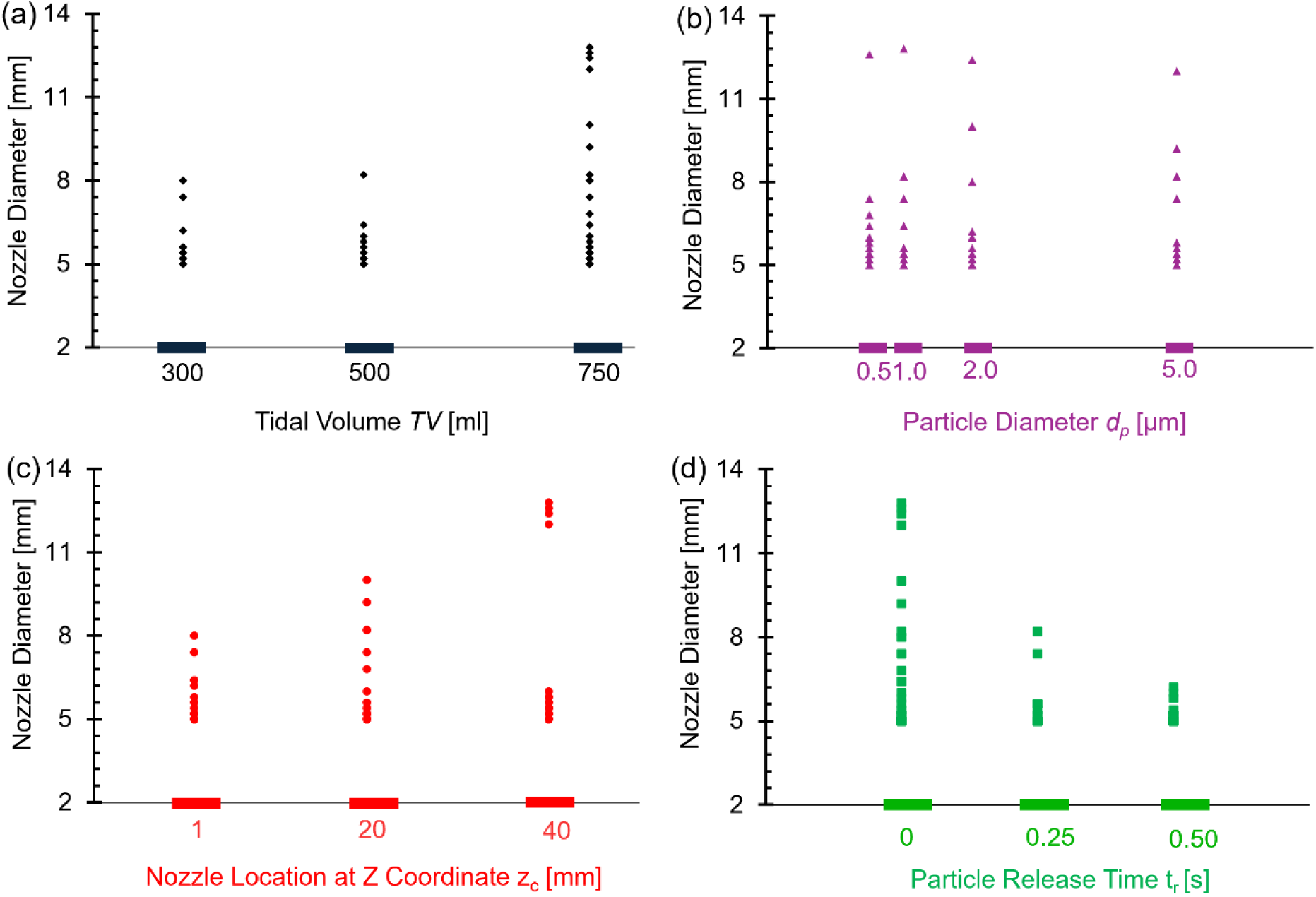
The CFPD-TDD nozzle diameter distributions under four different categories: (a) tidal volume (TV), (b) particle diameter 𝑑_𝑝_, (c) nozzle location at Z coordinates 𝑧_𝑐_, and (d) Particle release time instant.

Overall, the results indicate that in most scenarios, the optimal CFPD-TDD nozzle diameter 𝑑_𝑛_ is close to 5 mm, which corresponds to the lowest 𝐶𝑜𝑉 values in the uniformity comparisons (see **Figs. 13, 15, and 17**). Further analysis of the relationships between the optimal nozzle diameter and key parameters (𝑇𝑉, 𝑑_𝑝_, 𝑧_𝑐_, and 𝑡_𝑟_) reveals several notable observations. First, no significant variation in the optimal nozzle diameter is observed across the four investigated particle sizes (see **Figs. 12, 14, 16, and 18(b)**). Second, the range of viable nozzle diameters becomes wider under the highest tidal volume condition (𝑇𝑉 = 750 ml), extending from approximately 5 mm to 12.8 mm, compared with the lower TV conditions (see **Fig. 18(a)**). Third, as the particle release axial location 𝑧_𝑐_ increases, the optimal nozzle diameter tends to increase, indicating that larger injection regions may become feasible when particle release occurs closer to the throat (see **Fig. 18(c)**). Fourth, increasing the particle release time 𝑡_𝑟_ tends to narrow the feasible range of nozzle diameters, with most viable diameters falling below approximately 7.4 mm (see **Fig. 18(d)**).

These results highlight the complex and nonlinear relationships between the TDD nozzle configuration (i.e., 𝑑_𝑛_, 𝑥_𝑐_, 𝑦_𝑐_) and the governing inhalation and particle parameters (i.e., 𝑇𝑉, 𝑑_𝑝_, 𝑧_𝑐_, and 𝑡_𝑟_). Accurately identifying optimal nozzle configurations for targeted drug delivery, therefore, requires a systematic approach capable of capturing these multidimensional dependencies. This motivates the use of ML techniques to efficiently analyze and predict such complex relationships. The complete dataset of CFPD-determined optimal TDD nozzle locations ( 𝑥_𝑐_, 𝑦_𝑐_) and diameters ( 𝑑_𝑛_) under all investigated conditions is provided in **Appendix A**.

### 3.4 Enhanced Lobar Particle Deposition Uniformity with Machine Learning

#### 3.4.1 ML Model Performance

To comprehensively evaluate and compare the performance of all developed ML models (see Section 2.3.2), both MSE and MAE were calculated for the overall predictions as well as for each target variable (i.e., nozzle center’s X-coordinate 𝑥_𝑐_, nozzle center’s Y-coordinate 𝑦_𝑐_, and nozzle diameter 𝑑_𝑛_). The evaluation metrics were assessed on both the testing set and the independent validation set, as summarized in **Appendix B**. Based on the results shown in **Appendix B**, 3 models with the lowest MSE on the test set were identified, i.e., RM_D, RMD_D, and TF_D. The overall performance based on MSE and MAE can be intuitively compared across 16 ML models in **Fig. 19**.

**Figure 19:**
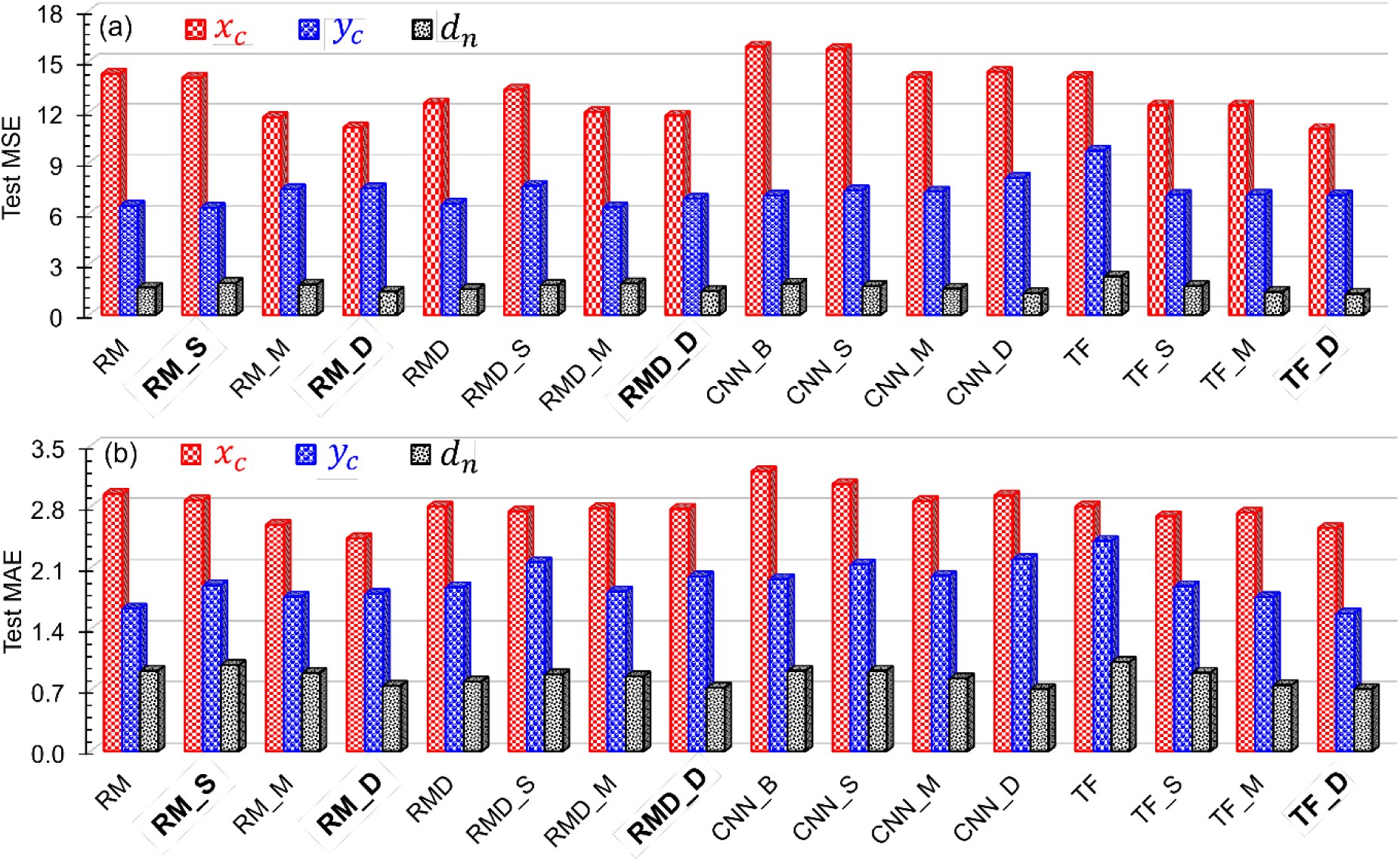
Performance comparisons of the 16 ML models developed in this study: (a) MSEs of target features on the testing set, and (b) MAEs of target features on the testing set.

In addition, TF_D, RM_S and TF_D have the lowest MSE on 𝑥_𝑐_, 𝑦_𝑐_ and 𝑑_𝑛_, respectively. Therefore, a hybrid ensemble model was created by combining the three models that performed best on each target feature, as determined by feature-wise MSE. These four top-performing models are marked in bold in **Appendix B**. Their corresponding MSE values are marked in bold and red for easier distinction.

#### 3.4.2 Cross Validation: CFPD-Driven vs. ML-Driven Targeted Drug Delivery

The raw data of the predicted nozzle parameters for two additional validation cases beyond the training dataset, obtained using the selected machine learning (ML) models (i.e., MixModel, RM_D, RMD_D, and TF_D), are provided in **Appendix C**. Two independent cases (i.e., Case I and Case II) were designed to evaluate whether ML-predicted nozzle configurations can improve the uniformity of drug particle deposition across the five lung lobes. To account for inter-subject variability in breathing patterns, the peak inspiratory flow rates 𝑄_max_ were set to 33 L/min and 46 L/min for Case I and Case II, respectively. A 2-second sinusoidal breathing waveform with an inhalation-to-exhalation ratio of 1:1 was applied in both cases. The four ML-driven models (i.e., MixModel, RM_D, RMD_D, and TF_D) were used to predict nozzle diameter and nozzle center (location), which were then directly implemented in subsequent CFD simulations.

For each case (i.e., Case I and Case II), baseline CFPD simulations were first performed using both CFPD-FMD and CFPD-TDD frameworks following the methodology described in Section 2.3. These simulations established reference deposition performance and optimal nozzle configurations based on first-principles simulation results. Subsequently, additional CFPD simulations were conducted by injecting particles with nozzle parameters (i.e., nozzle diameter and center locations) predicted by each of the 4 selected ML models, under the same boundary conditions. The DFs across five lobes from the ML-driven TDD simulations were then compared against the CFPD-FMD and CFPD-TDD results to assess ML model performance. The coefficient of variation (𝐶𝑜𝑉) of drug deposition across the five lobes was used as the primary metric to quantify deposition uniformity. Detailed cross-validation results for all cases and models are summarized in **Appendix C**.

**Figures 20 (a) & (b)** compare DFs among the traditional CFPD-FMD strategy, CFPD-TDD strategy, and 4 ML-driven TDD strategies across the five lobes (i.e., RML, RLL, LLL, RUL, and LUL) for Cases I and II, respectively. **Figure 20** also evaluates the effectiveness of each set of predicted nozzle parameters in achieving uniform drug distribution across all lobes. As the ground-truth data, the CFPD-TDD strategies exhibit the best performance in uneven deposition patterns (i.e., the highest 𝐶𝑜𝑉). Followed by one of ML-driven nozzle settings based on the RM_D model, it also obtains decent uniform drug distributions across all lobes in both Case I and Case II, with the calculated 𝐶𝑜𝑉 values of 44.19% and 38.21%, respectively. The other three ML models, i.e., MixModel, RMD_D, and TF_D, perform better than the traditional CFPD-FMD strategy in Case-I, but underperform in Case-II, indicating their poor robustness. Thus, based on cross-validation of the four tested ML models, the RM_D model could be a prime choice for the user-centered smart inhaler prototype to enhance uniformity of TDD in small airways compared with other ML models.

**Figure 20:**
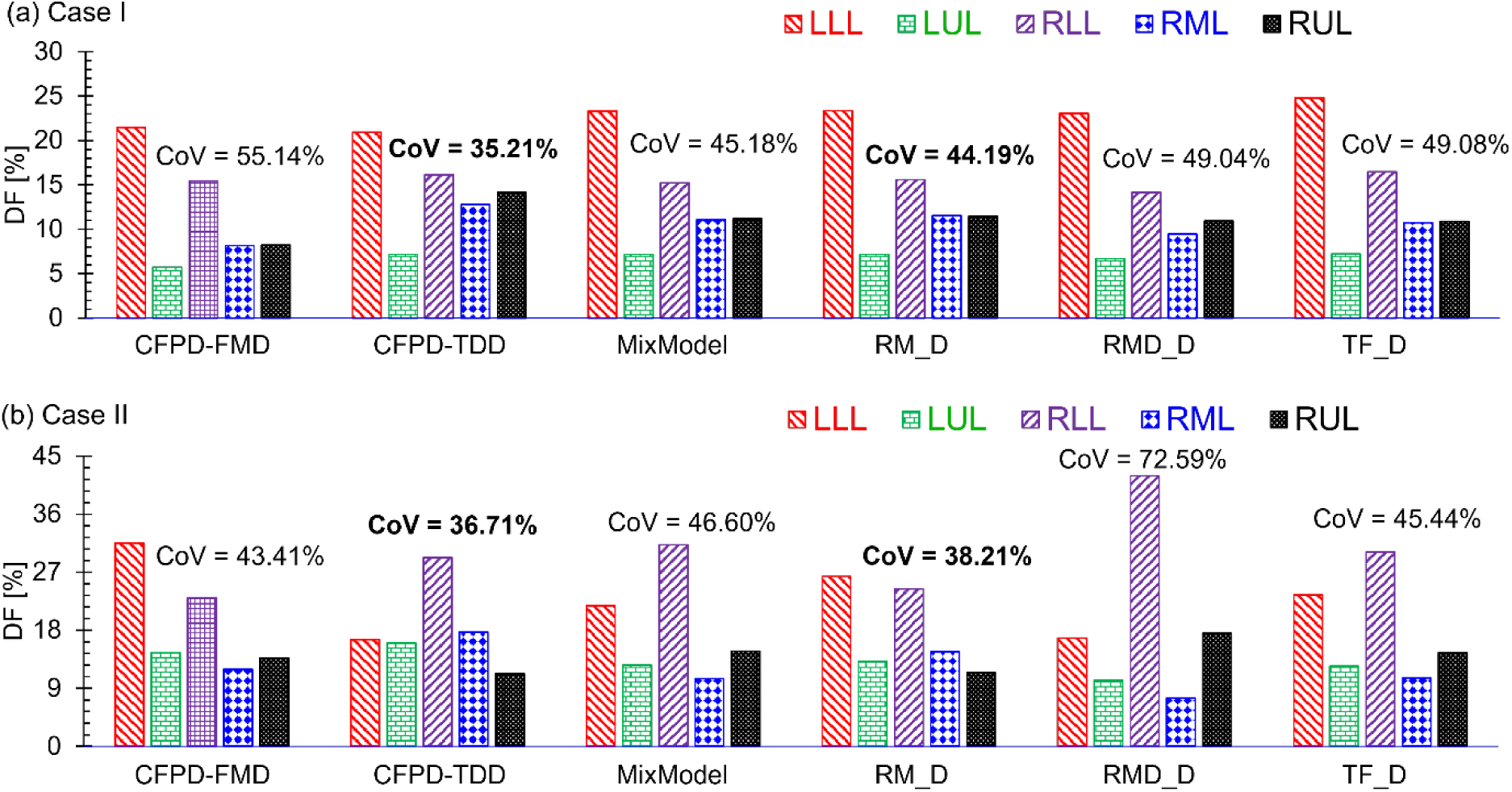
Comparison of DFs and CoV for CFPD-TDD vs. ML-TDD strategies in five lobar lobes: (a) Case I, and (b) Case II.

## 4. Conclusions

This study presented a CFPD-informed, ML-empowered, user-centered smart inhaler framework to establish a personalized TDD strategy via inhalation therapy that efficiently and uniformly deposits aerosolized drug particles in the small airways across five lobes. To identify the most effective predictive ML algorithm for embedding and deployment in the smart inhaler, 16 ML models were trained on 108 CFPD datasets and systematically compared. The four best-performing ML models (i.e., MixModel, RM_D, RMD_D, and TF_D) were carried forward for cross-validation. CFPD-informed ML models can predict the optimal nozzle position, the maximum feasible nozzle diameter, and the particle release timing from patient- and drug-specific inputs (e.g., inhalation flow rate and particle size), enabling control of drug delivery in small airways beyond G10 across all lobes. Across two external validation cases, ML-driven TDD strategies reduced inter-lobar variability relative to CFPD-TDD strategies, with the RM_D model achieving the lowest average CoV while maintaining competitive total drug particle deposition in small airways. These findings support the feasibility and potential of integrating the trained ML models into the user-centered smart inhaler prototype to enhance uniformity and efficiency of pulmonary TDD. The key findings of this study are summarized below. Specifically, i.e.,

1. Most of the inhaled smaller particles (i.e., 𝑑_𝑝_=0.5, 1, and 2 μm) can pass through the respiratory tract and reach the deeper lung airways beyond G10, while a few drug particles deposited in the mouth-to-trachea region. On the other hand, larger particles (i.e., 𝑑_𝑝_=5 μm) show more particle deposition in the mouth-to-trachea region than smaller particles. This study confirms that particles with 𝑑_𝑝_ between 0.5 and 2 μm have the potential to be delivered into small airways without high deposition in the upper airway (i.e., the mouth-to-trachea region).
2. The tidal volume (𝑇𝑉), particle release position (𝑧_𝑐_), and particle release time (𝑡_𝑟_) has a significant impact on the optimal nozzle diameter for CFPD-TDD, while particle size near has no impact on the selection of nozzle diameter ranges in CFPD-TDD strategies.
3. Although the conventional CFPD-FMD strategies achieve some degree of even distribution across the lung lobes, CFPD-TDD offers a key advantage by significantly reducing non-targeted deposition. This advantage of the CFPD-TDD strategy allows a greater proportion of the drug to reach the intended small airway, ultimately leading to a more efficient and safer therapeutic outcome.
4. Compared with the conventional CFPD-FMD strategy, the ML-based approaches, such as the RMD_M model, notably improve the consistency of particle deposition across lung lobes, although it still underperforms when compared with the ground-truth CFPD-TDD strategies. These results demonstrate that the ML-driven models can reproduce the deposition trends captured by high-fidelity CFPD and even enhance evenness across five lobes. Therefore, integrating ML predictions into the TDD framework provides a computationally efficient alternative to CFPD-TDD planning for small airway disease treatment innovation.

## 5. Limitations of This Study and Future Work

The simplification and assumptions of this study are as follows:

1. An idealized sinusoidal waveform was used for the CFPD simulation instead of a realistic breathing profile (Colby et al., 2016).
2. This study employed a one-way air-particle coupling approach, neglecting particle effects on the airflow field and particle-particle interactions, such as agglomeration or de-agglomeration (Elghobashi & Truesdell, 1993).
3. Monodisperse drug particle diameters were used in CFPD simulations instead of more realistic polydisperse particle size distributions (Cui et al., 2018).
4. This study assumed that the glottis is static during the breathing cycle by neglecting the influence of glottal motion on airflow dynamics (Zhao et al., 2020).
5. This study employed a single subject-specific airway geometry, which leads to the negligence of inter-subject variability impacts, including airway anatomical heterogeneity, which can significantly affect pulmonary air-particle transport dynamics.
6. This study is purely computational and serves as a proof-of-concept investigation based on CFPD simulations and machine learning predictions. No in vitro, benchtop, or in vivo experiments were conducted to validate the manufacturability, real-time controllability, or clinical feasibility of a smart inhaler capable of dynamically adjusting nozzle diameter and location.

To address those above-mentioned simplifications and assumptions, the following mechanisms will be considered in future studies, i.e.,

1. Include a more realistic breathing profile in the CFPD simulations to provide more accurate insights into particle behavior in real-life respiratory conditions.
2. Utilize a two-way coupled Euler–Lagrange approach or discrete element method (DEM) to simulate realistic particle-airflow interactions and particle-particle interactions.
3. Incorporate more realistic polydisperse particle size distributions in future CFPD simulations.
4. Develop a CFPD model with glottis motion to evaluate its impact on pulmonary airflow and particle transport.
5. Expand the training and testing datasets with CFPD simulations using multiple subject-specific and disease-specific airway geometries, enabling improved generalizability for the ML model to enhance inhalation therapy outcomes.
6. Fabricate and experimentally evaluate a smart inhaler prototype through in vitro aerosol characterization (e.g., cascade impaction or laser diffraction), benchtop deposition testing, and ultimately preclinical or clinical studies to validate the feasibility, robustness, and translational performance of the ML-guided nozzle design framework.

## Supporting information

Appendix A: Complete Dataset for ML Training and Testing (108 CFPD Simulations)

Appendix B: Machine learning performance and model selection

Appendix C: Comparisons of Performances between ML-Driven TDD and CFPD-TDD Strategies on Cross-Validation Cases

## Acknowledgments

This material is based upon work supported by the National Science Foundation under Award Number 2429582. The authors gratefully acknowledge the support previously provided by Canopy Healthtech (Dr. Rachel Lane and Dr. Stacie Sober). The use of Ansys software (Ansys Inc., Canonsburg, PA) as part of the Ansys–CBBL academic partnership is also gratefully acknowledged (Thierry Marchal and Vishal Ganore).

## Notes

### Competing Interest Statement

The authors have declared no competing interest.

