## Appendix A: Complete Dataset for ML Training and Testing (108 CFPD Simulations) for "Computational Fluid Particle Dynamics-Informed Machine Learning Prototype for a User-Centered Smart Inhaler Enabling Uniform Drug Delivery to Small Airways"

| Lobe Name and Its Zone ID Obtained from CFPD Simulatio | | | | | | | |
|---|---|---|---|---|---|---|---|
| | PFR [L/min] | VM [m/s] | PD [mm] | Z [mm] | PRT [s] | RT | X-C [mm] |
| **Case 001** | 28.27 | 1.5 | 0.0005 | 1 | 0 | FMD | 0 |
| | 28.27 | 1.5 | 0.0005 | 1 | 0 | TDD | 2.2 |
| Case 002 | 28.27 | 1.5 | 0.0005 | 20 | 0 | FMD | 0 |
| | 28.27 | 1.5 | 0.0005 | 20 | 0 | TDD | 0.5 |
| Case 003 | 28.27 | 1.5 | 0.0005 | 40 | 0 | FMD | 0 |
| | 28.27 | 1.5 | 0.0005 | 40 | 0 | TDD | 4.45 |
| Case 004 | 28.27 | 1.5 | 0.0005 | 1 | 0.25 | FMD | 0 |
| | 28.27 | 1.5 | 0.0005 | 1 | 0.25 | TDD | 3.05 |
| **Case 005** | 28.27 | 1.5 | 0.0005 | 20 | 0.25 | FMD | 0 |
| | 28.27 | 1.5 | 0.0005 | 20 | 0.25 | TDD | 2.25 |
| Case 006 | 28.27 | 1.5 | 0.0005 | 40 | 0.25 | FMD | 0 |
| | 28.27 | 1.5 | 0.0005 | 40 | 0.25 | TDD | 3.25 |
| Case 007 | 28.27 | 1.5 | 0.0005 | 1 | 0.5 | FMD | 0 |
| | 28.27 | 1.5 | 0.0005 | 1 | 0.5 | TDD | 6.1 |
| Case 008 | 28.27 | 1.5 | 0.0005 | 20 | 0.5 | FMD | 0 |
| | 28.27 | 1.5 | 0.0005 | 20 | 0.5 | TDD | 4.25 |
| Case 009 | 28.27 | 1.5 | 0.0005 | 40 | 0.5 | FMD | 0 |
| | 28.27 | 1.5 | 0.0005 | 40 | 0.5 | TDD | 1.1 |
| Case 010 | 28.27 | 1.5 | 0.001 | 1 | 0 | FMD | 0 |
| | 28.27 | 1.5 | 0.001 | 1 | 0 | TDD | -4.9 |
| **Case 011** | 28.27 | 1.5 | 0.001 | 20 | 0 | FMD | 0 |
| | 28.27 | 1.5 | 0.001 | 20 | 0 | TDD | 0.5 |
| **Case 012** | 28.27 | 1.5 | 0.001 | 40 | 0 | FMD | 0 |
| | 28.27 | 1.5 | 0.001 | 40 | 0 | TDD | -0.7 |
| **Case 013** | 28.27 | 1.5 | 0.001 | 1 | 0.25 | FMD | 0 |
| | 28.27 | 1.5 | 0.001 | 1 | 0.25 | TDD | -6 |
| Case 014 | 28.27 | 1.5 | 0.001 | 20 | 0.25 | FMD | 0 |
| | 28.27 | 1.5 | 0.001 | 20 | 0.25 | TDD | 2.4 |
| Case 015 | 28.27 | 1.5 | 0.001 | 40 | 0.25 | FMD | 0 |
| | 28.27 | 1.5 | 0.001 | 40 | 0.25 | TDD | 1.6 |
| Case 016 | 28.27 | 1.5 | 0.001 | 1 | 0.5 | FMD | 0 |
| | 28.27 | 1.5 | 0.001 | 1 | 0.5 | TDD | 5.2 |
| Case 017 | 28.27 | 1.5 | 0.001 | 20 | 0.5 | FMD | 0 |
| | 28.27 | 1.5 | 0.001 | 20 | 0.5 | TDD | 5 |
| Case 018 | 28.27 | 1.5 | 0.001 | 40 | 0.5 | FMD | 0 |
| | 28.27 | 1.5 | 0.001 | 40 | 0.5 | TDD | 6 |
| **Case 019** | 28.27 | 1.5 | 0.002 | 1 | 0 | FMD | 0 |
| | 28.27 | 1.5 | 0.002 | 1 | 0 | TDD | 2 |
| Case 020 | 28.27 | 1.5 | 0.002 | 20 | 0 | FMD | 0 |
| | 28.27 | 1.5 | 0.002 | 20 | 0 | TDD | 1 |
| Case 021 | 28.27 | 1.5 | 0.002 | 40 | 0 | FMD | 0 |
| | 28.27 | 1.5 | 0.002 | 40 | 0 | TDD | -0.9 |
| Case 022 | 28.27 | 1.5 | 0.002 | 1 | 0.25 | FMD | 0 |

| | | | | | | | |
|---|---|---|---|---|---|---|---|
| Case 022 | 28.27 | 1.5 | 0.002 | 1 | 0.25 | TDD | -3 |
| Case 023 | 28.27 | 1.5 | 0.002 | 20 | 0.25 | FMD | 0 |
| | 28.27 | 1.5 | 0.002 | 20 | 0.25 | TDD | 2 |
| Case 024 | 28.27 | 1.5 | 0.002 | 40 | 0.25 | FMD | 0 |
| | 28.27 | 1.5 | 0.002 | 40 | 0.25 | TDD | 2.5 |
| Case 025 | 28.27 | 1.5 | 0.002 | 1 | 0.5 | FMD | 0 |
| | 28.27 | 1.5 | 0.002 | 1 | 0.5 | TDD | 5.1 |
| Case 026 | 28.27 | 1.5 | 0.002 | 20 | 0.5 | FMD | 0 |
| | 28.27 | 1.5 | 0.002 | 20 | 0.5 | TDD | 3.1 |
| Case 027 | 28.27 | 1.5 | 0.002 | 40 | 0.5 | FMD | 0 |
| | 28.27 | 1.5 | 0.002 | 40 | 0.5 | TDD | 0.1 |
| Case 028 | 28.27 | 1.5 | 0.005 | 1 | 0 | FMD | 0 |
| | 28.27 | 1.5 | 0.005 | 1 | 0 | TDD | -7 |
| Case 029 | 28.27 | 1.5 | 0.005 | 20 | 0 | FMD | 0 |
| | 28.27 | 1.5 | 0.005 | 20 | 0 | TDD | -4.9 |
| Case 030 | 28.27 | 1.5 | 0.005 | 40 | 0 | FMD | 0 |
| | 28.27 | 1.5 | 0.005 | 40 | 0 | TDD | -4 |
| **Case 031** | 28.27 | 1.5 | 0.005 | 1 | 0.25 | FMD | 0 |
| | 28.27 | 1.5 | 0.005 | 1 | 0.25 | TDD | 2.3 |
| Case 032 | 28.27 | 1.5 | 0.005 | 20 | 0.25 | FMD | 0 |
| | 28.27 | 1.5 | 0.005 | 20 | 0.25 | TDD | 0.2 |
| Case 033 | 28.27 | 1.5 | 0.005 | 40 | 0.25 | FMD | 0 |
| | 28.27 | 1.5 | 0.005 | 40 | 0.25 | TDD | 6.5 |
| Case 034 | 28.27 | 1.5 | 0.005 | 1 | 0.5 | FMD | 0 |
| | 28.27 | 1.5 | 0.005 | 1 | 0.5 | TDD | 4.1 |
| Case 035 | 28.27 | 1.5 | 0.005 | 20 | 0.5 | FMD | 0 |
| | 28.27 | 1.5 | 0.005 | 20 | 0.5 | TDD | 1.1 |
| Case 036 | 28.27 | 1.5 | 0.005 | 40 | 0.5 | FMD | 0 |
| | 28.27 | 1.5 | 0.005 | 40 | 0.5 | TDD | -1.5 |
| **Case 037** | 47.12 | 2.5 | 0.0005 | 1 | 0 | FMD | 0 |
| | 47.12 | 2.5 | 0.0005 | 1 | 0 | TDD | 5 |
| Case 038 | 47.12 | 2.5 | 0.0005 | 20 | 0 | FMD | 0 |
| | 47.12 | 2.5 | 0.0005 | 20 | 0 | TDD | 5.5 |
| Case 039 | 47.12 | 2.5 | 0.0005 | 40 | 0 | FMD | 0 |
| | 47.12 | 2.5 | 0.0005 | 40 | 0 | TDD | -1.5 |
| Case 040 | 47.12 | 2.5 | 0.0005 | 1 | 0.25 | FMD | 0 |
| | 47.12 | 2.5 | 0.0005 | 1 | 0.25 | TDD | 5.3 |
| Case 041 | 47.12 | 2.5 | 0.0005 | 20 | 0.25 | FMD | 0 |
| | 47.12 | 2.5 | 0.0005 | 20 | 0.25 | TDD | -4.7 |
| Case 042 | 47.12 | 2.5 | 0.0005 | 40 | 0.25 | FMD | 0 |
| | 47.12 | 2.5 | 0.0005 | 40 | 0.25 | TDD | -4.3 |
| **Case 043** | 47.12 | 2.5 | 0.0005 | 1 | 0.5 | FMD | 0 |
| | 47.12 | 2.5 | 0.0005 | 1 | 0.5 | TDD | 3.4 |
| Case 044 | 47.12 | 2.5 | 0.0005 | 20 | 0.5 | FMD | 0 |
| | 47.12 | 2.5 | 0.0005 | 20 | 0.5 | TDD | 6.7 |

| | | | | | | | |
|---|---|---|---|---|---|---|---|
| Case 045 | 47.12 | 2.5 | 0.0005 | 40 | 0.5 | FMD | 0 |
| | 47.12 | 2.5 | 0.0005 | 40 | 0.5 | TDD | 6.9 |
| **Case 046** | 47.12 | 2.5 | 0.001 | 1 | 0 | FMD | 0 |
| | 47.12 | 2.5 | 0.001 | 1 | 0 | TDD | -0.2 |
| Case 047 | 47.12 | 2.5 | 0.001 | 20 | 0 | FMD | 0 |
| | 47.12 | 2.5 | 0.001 | 20 | 0 | TDD | 4.8 |
| Case 048 | 47.12 | 2.5 | 0.001 | 40 | 0 | FMD | 0 |
| | 47.12 | 2.5 | 0.001 | 40 | 0 | TDD | -1.6 |
| Case 049 | 47.12 | 2.5 | 0.001 | 1 | 0.25 | FMD | 0 |
| | 47.12 | 2.5 | 0.001 | 1 | 0.25 | TDD | -4.2 |
| Case 050 | 47.12 | 2.5 | 0.001 | 20 | 0.25 | FMD | 0 |
| | 47.12 | 2.5 | 0.001 | 20 | 0.25 | TDD | -3.9 |
| Case 051 | 47.12 | 2.5 | 0.001 | 40 | 0.25 | FMD | 0 |
| | 47.12 | 2.5 | 0.001 | 40 | 0.25 | TDD | -3.3 |
| Case 052 | 47.12 | 2.5 | 0.001 | 1 | 0.5 | FMD | 0 |
| | 47.12 | 2.5 | 0.001 | 1 | 0.5 | TDD | -3.5 |
| Case 053 | 47.12 | 2.5 | 0.001 | 20 | 0.5 | FMD | 0 |
| | 47.12 | 2.5 | 0.001 | 20 | 0.5 | TDD | 6.9 |
| Case 054 | 47.12 | 2.5 | 0.001 | 40 | 0.5 | FMD | 0 |
| | 47.12 | 2.5 | 0.001 | 40 | 0.5 | TDD | 1.6 |
| Case 055 | 47.12 | 2.5 | 0.002 | 1 | 0 | FMD | 0 |
| | 47.12 | 2.5 | 0.002 | 1 | 0 | TDD | 0 |
| Case 056 | 47.12 | 2.5 | 0.002 | 20 | 0 | FMD | 0 |
| | 47.12 | 2.5 | 0.002 | 20 | 0 | TDD | 4.9 |
| **Case 057** | 47.12 | 2.5 | 0.002 | 40 | 0 | FMD | 0 |
| | 47.12 | 2.5 | 0.002 | 40 | 0 | TDD | -1.5 |
| Case 058 | 47.12 | 2.5 | 0.002 | 1 | 0.25 | FMD | 0 |
| | 47.12 | 2.5 | 0.002 | 1 | 0.25 | TDD | 0.6 |
| Case 059 | 47.12 | 2.5 | 0.002 | 20 | 0.25 | FMD | 0 |
| | 47.12 | 2.5 | 0.002 | 20 | 0.25 | TDD | 5 |
| Case 060 | 47.12 | 2.5 | 0.002 | 40 | 0.25 | FMD | 0 |
| | 47.12 | 2.5 | 0.002 | 40 | 0.25 | TDD | -1 |
| Case 061 | 47.12 | 2.5 | 0.002 | 1 | 0.5 | FMD | 0 |
| | 47.12 | 2.5 | 0.002 | 1 | 0.5 | TDD | 2 |
| Case 062 | 47.12 | 2.5 | 0.002 | 20 | 0.5 | FMD | 0 |
| | 47.12 | 2.5 | 0.002 | 20 | 0.5 | TDD | 0 |
| **Case 063** | 47.12 | 2.5 | 0.002 | 40 | 0.5 | FMD | 0 |
| | 47.12 | 2.5 | 0.002 | 40 | 0.5 | TDD | 7.2 |
| Case 064 | 47.12 | 2.5 | 0.005 | 1 | 0 | FMD | 0 |
| | 47.12 | 2.5 | 0.005 | 1 | 0 | TDD | 1.3 |
| Case 065 | 47.12 | 2.5 | 0.005 | 20 | 0 | FMD | 0 |
| | 47.12 | 2.5 | 0.005 | 20 | 0 | TDD | 0.1 |
| Case 066 | 47.12 | 2.5 | 0.005 | 40 | 0 | FMD | 0 |
| | 47.12 | 2.5 | 0.005 | 40 | 0 | TDD | -0.5 |
| Case 067 | 47.12 | 2.5 | 0.005 | 1 | 0.25 | FMD | 0 |

| | | | | | | | |
|---|---|---|---|---|---|---|---|
| Case 067 | 47.12 | 2.5 | 0.005 | 1 | 0.25 | TDD | 6.6 |
| **Case 068** | 47.12 | 2.5 | 0.005 | 20 | 0.25 | FMD | 0 |
| | 47.12 | 2.5 | 0.005 | 20 | 0.25 | TDD | 1 |
| Case 069 | 47.12 | 2.5 | 0.005 | 40 | 0.25 | FMD | 0 |
| | 47.12 | 2.5 | 0.005 | 40 | 0.25 | TDD | -3 |
| **Case 070** | 47.12 | 2.5 | 0.005 | 1 | 0.5 | FMD | 0 |
| | 47.12 | 2.5 | 0.005 | 1 | 0.5 | TDD | 2.5 |
| Case 071 | 47.12 | 2.5 | 0.005 | 20 | 0.5 | FMD | 0 |
| | 47.12 | 2.5 | 0.005 | 20 | 0.5 | TDD | 0 |
| Case 072 | 47.12 | 2.5 | 0.005 | 40 | 0.5 | FMD | 0 |
| | 47.12 | 2.5 | 0.005 | 40 | 0.5 | TDD | 1.1 |
| Case 073 | 70.69 | 3.75 | 0.0005 | 1 | 0 | FMD | 0 |
| | 70.69 | 3.75 | 0.0005 | 1 | 0 | TDD | -1 |
| Case 074 | 70.69 | 3.75 | 0.0005 | 20 | 0 | FMD | 0 |
| | 70.69 | 3.75 | 0.0005 | 20 | 0 | TDD | -0.6 |
| Case 075 | 70.69 | 3.75 | 0.0005 | 40 | 0 | FMD | 0 |
| | 70.69 | 3.75 | 0.0005 | 40 | 0 | TDD | 2.9 |
| Case 076 | 70.69 | 3.75 | 0.0005 | 1 | 0.25 | FMD | 0 |
| | 70.69 | 3.75 | 0.0005 | 1 | 0.25 | TDD | 2.1 |
| **Case 077** | 70.69 | 3.75 | 0.0005 | 20 | 0.25 | FMD | 0 |
| | 70.69 | 3.75 | 0.0005 | 20 | 0.25 | TDD | -4.6 |
| **Case 078** | 70.69 | 3.75 | 0.0005 | 40 | 0.25 | FMD | 0 |
| | 70.69 | 3.75 | 0.0005 | 40 | 0.25 | TDD | -3.7 |
| Case 079 | 70.69 | 3.75 | 0.0005 | 1 | 0.5 | FMD | 0 |
| | 70.69 | 3.75 | 0.0005 | 1 | 0.5 | TDD | -5.9 |
| **Case 080** | 70.69 | 3.75 | 0.0005 | 20 | 0.5 | FMD | 0 |
| | 70.69 | 3.75 | 0.0005 | 20 | 0.5 | TDD | -7.1 |
| **Case 081** | 70.69 | 3.75 | 0.0005 | 40 | 0.5 | FMD | 0 |
| | 70.69 | 3.75 | 0.0005 | 40 | 0.5 | TDD | 3.2 |
| **Case 082** | 70.69 | 3.75 | 0.001 | 1 | 0 | FMD | 0 |
| | 70.69 | 3.75 | 0.001 | 1 | 0 | TDD | -0.9 |
| Case 083 | 70.69 | 3.75 | 0.001 | 20 | 0 | FMD | 0 |
| | 70.69 | 3.75 | 0.001 | 20 | 0 | TDD | -1.1 |
| **Case 084** | 70.69 | 3.75 | 0.001 | 40 | 0 | FMD | 0 |
| | 70.69 | 3.75 | 0.001 | 40 | 0 | TDD | 3 |
| Case 085 | 70.69 | 3.75 | 0.001 | 1 | 0.25 | FMD | 0 |
| | 70.69 | 3.75 | 0.001 | 1 | 0.25 | TDD | -3.4 |
| Case 086 | 70.69 | 3.75 | 0.001 | 20 | 0.25 | FMD | 0 |
| | 70.69 | 3.75 | 0.001 | 20 | 0.25 | TDD | 3.4 |
| Case 087 | 70.69 | 3.75 | 0.001 | 40 | 0.25 | FMD | 0 |
| | 70.69 | 3.75 | 0.001 | 40 | 0.25 | TDD | -5.2 |
| Case 088 | 70.69 | 3.75 | 0.001 | 1 | 0.5 | FMD | 0 |
| | 70.69 | 3.75 | 0.001 | 1 | 0.5 | TDD | -7.4 |
| Case 089 | 70.69 | 3.75 | 0.001 | 20 | 0.5 | FMD | 0 |
| | 70.69 | 3.75 | 0.001 | 20 | 0.5 | TDD | 3.8 |

| | | | | | | | |
|---|---|---|---|---|---|---|---|
| Case 090 | 70.69 | 3.75 | 0.001 | 40 | 0.5 | FMD | 0 |
| | 70.69 | 3.75 | 0.001 | 40 | 0.5 | TDD | 3 |
| Case 091 | 70.69 | 3.75 | 0.002 | 1 | 0 | FMD | 0 |
| | 70.69 | 3.75 | 0.002 | 1 | 0 | TDD | -0.8 |
| Case 092 | 70.69 | 3.75 | 0.002 | 20 | 0 | FMD | 0 |
| | 70.69 | 3.75 | 0.002 | 20 | 0 | TDD | -0.2 |
| Case 093 | 70.69 | 3.75 | 0.002 | 40 | 0 | FMD | 0 |
| | 70.69 | 3.75 | 0.002 | 40 | 0 | TDD | 2.8 |
| Case 094 | 70.69 | 3.75 | 0.002 | 1 | 0.25 | FMD | 0 |
| | 70.69 | 3.75 | 0.002 | 1 | 0.25 | TDD | 0.8 |
| Case 095 | 70.69 | 3.75 | 0.002 | 20 | 0.25 | FMD | 0 |
| | 70.69 | 3.75 | 0.002 | 20 | 0.25 | TDD | 2.5 |
| **Case 096** | 70.69 | 3.75 | 0.002 | 40 | 0.25 | FMD | 0 |
| | 70.69 | 3.75 | 0.002 | 40 | 0.25 | TDD | -4.4 |
| Case 097 | 70.69 | 3.75 | 0.002 | 1 | 0.5 | FMD | 0 |
| | 70.69 | 3.75 | 0.002 | 1 | 0.5 | TDD | 0.9 |
| Case 098 | 70.69 | 3.75 | 0.002 | 20 | 0.5 | FMD | 0 |
| | 70.69 | 3.75 | 0.002 | 20 | 0.5 | TDD | 6.8 |
| Case 099 | 70.69 | 3.75 | 0.002 | 40 | 0.5 | FMD | 0 |
| | 70.69 | 3.75 | 0.002 | 40 | 0.5 | TDD | 1.5 |
| Case 100 | 70.69 | 3.75 | 0.005 | 1 | 0 | FMD | 0 |
| | 70.69 | 3.75 | 0.005 | 1 | 0 | TDD | 0.9 |
| Case 101 | 70.69 | 3.75 | 0.005 | 20 | 0 | FMD | 0 |
| | 70.69 | 3.75 | 0.005 | 20 | 0 | TDD | 0.85 |
| Case 102 | 70.69 | 3.75 | 0.005 | 40 | 0 | FMD | 0 |
| | 70.69 | 3.75 | 0.005 | 40 | 0 | TDD | 2.6 |
| Case 103 | 70.69 | 3.75 | 0.005 | 1 | 0.25 | FMD | 0 |
| | 70.69 | 3.75 | 0.005 | 1 | 0.25 | TDD | 2.5 |
| Case 104 | 70.69 | 3.75 | 0.005 | 20 | 0.25 | FMD | 0 |
| | 70.69 | 3.75 | 0.005 | 20 | 0.25 | TDD | 5.1 |
| Case 105 | 70.69 | 3.75 | 0.005 | 40 | 0.25 | FMD | 0 |
| | 70.69 | 3.75 | 0.005 | 40 | 0.25 | TDD | 2.85 |
| Case 106 | 70.69 | 3.75 | 0.005 | 1 | 0.5 | FMD | 0 |
| | 70.69 | 3.75 | 0.005 | 1 | 0.5 | TDD | -4.4 |
| **Case 107** | 70.69 | 3.75 | 0.005 | 20 | 0.5 | FMD | 0 |
| | 70.69 | 3.75 | 0.005 | 20 | 0.5 | TDD | -1.3 |
| Case 108 | 70.69 | 3.75 | 0.005 | 40 | 0.5 | FMD | 0 |
| | 70.69 | 3.75 | 0.005 | 40 | 0.5 | TDD | 5.25 |

| |
|---|
| **Note:** |
| **PFR:** Flow Rate at the Inhalation Peak Instant |
| **VM:** Velocity of Magnitude at the Inhalation Peak Instant |

| |
|---|
| **PD:** Particle Diameter |
| **Z:** Z-Coordinate |
| **PRT:** Particle Release Time |
| **X-C:** Optimal Nozzle Center X-Coordinate |
| **Y-C**: Optimal Nozzle Center Y-Coordinate |
| **ND:** Optimal Nozzle Diameter |
| **DPN:** Deposition Particle Number |
| **TDD:** Target Drug Delivery |
| **TRPN:** Total Release Particle Number |
| **DF:** Deposition Fraction |
| **LL:** Left Lower Outlets |
| **LU:** Left Upper Outlets |
| **RL:** Right Lower Outlets |
| **RM:** Right Middle Outlets |
| **RT:** Release Type |
| **RU:** Right Upper Outlets |
| **CoV:** Coefficient of Variation |
| **FMR:** Full mouth Release |

| ...n | | | LLL (ID=35) | | LUL (ID=40) | | RLL (ID=32) | |
|---|---|---|---|---|---|---|---|---|
| Y-C [mm] | ND [mm] | TRPN | DPN-LL | DF-LL | DPN-LU | DF-LU | DPN-RL | DF-RL |
| 0 | 20 | 10036 | 3021 | 0.3010 | 1779 | 0.1773 | 1806 | 0.1800 |
| 4.9 | 7.4 | | | | | | | |
| 0 | 20 | 10036 | 2908 | 0.2898 | 1183 | 0.1179 | 2373 | 0.2364 |
| 3 | 5 | | | | | | | |
| 0 | 20 | 10036 | 3272 | 0.3260 | 1159 | 0.1155 | 2514 | 0.2505 |
| 4.7 | 5.4 | | | | | | | |
| 0 | 20 | 10036 | 1986 | 0.1979 | 1242 | 0.1238 | 2193 | 0.2185 |
| 5.8 | 5.6 | | | | | | | |
| 0 | 20 | 10036 | 2060 | 0.2053 | 1120 | 0.1116 | 2233 | 0.2225 |
| 6.75 | 5 | | | | | | | |
| 0 | 20 | 10036 | 2070 | 0.2063 | 1245 | 0.1241 | 2238 | 0.2230 |
| 4.5 | 5 | | | | | | | |
| 0 | 20 | 10036 | 1691 | 0.1685 | 268 | 0.0267 | 1115 | 0.1111 |
| -1.9 | 5 | | | | | | | |
| 0 | 20 | 10036 | 1724 | 0.1718 | 267 | 0.0266 | 1354 | 0.1349 |
| -0.25 | 5 | | | | | | | |
| 0 | 20 | 10036 | 1684 | 0.1678 | 345 | 0.0344 | 1477 | 0.1472 |
| -5.4 | 5.2 | | | | | | | |
| 0 | 20 | 10036 | 2983 | 0.2972 | 1798 | 0.1792 | 1823 | 0.1816 |
| 1.4 | 5 | | | | | | | |
| 0 | 20 | 10036 | 2922 | 0.2912 | 1152 | 0.1148 | 2395 | 0.2386 |
| 3.5 | 5 | | | | | | | |
| 0 | 20 | 10036 | 3244 | 0.3232 | 1223 | 0.1219 | 2496 | 0.2487 |
| 3.3 | 5.6 | | | | | | | |
| 0 | 20 | 10036 | 2009 | 0.2002 | 1177 | 0.1173 | 2197 | 0.2189 |
| -4.5 | 5 | | | | | | | |
| 0 | 20 | 10036 | 2078 | 0.2071 | 1200 | 0.1196 | 2200 | 0.2192 |
| 5.4 | 5.6 | | | | | | | |
| 0 | 20 | 10036 | 2008 | 0.2001 | 1198 | 0.1194 | 2241 | 0.2233 |
| 3 | 5 | | | | | | | |
| 0 | 20 | 10036 | 1965 | 0.1958 | 226 | 0.0225 | 1081 | 0.1077 |
| -4.8 | 5.4 | | | | | | | |
| 0 | 20 | 10036 | 1670 | 0.1664 | 279 | 0.0278 | 1356 | 0.1351 |
| -1 | 5 | | | | | | | |
| 0 | 20 | 10036 | 1639 | 0.1633 | 390 | 0.0389 | 1450 | 0.1445 |
| 0.5 | 5 | | | | | | | |
| 0 | 20 | 10036 | 3001 | 0.2990 | 1782 | 0.1776 | 1868 | 0.1861 |
| 5 | 8 | | | | | | | |
| 0 | 20 | 10036 | 2976 | 0.2965 | 1151 | 0.1147 | 2324 | 0.2316 |
| 3 | 5 | | | | | | | |
| 0 | 20 | 10036 | 3267 | 0.3255 | 1176 | 0.1172 | 2523 | 0.2514 |
| 3.1 | 5.2 | | | | | | | |
| 0 | 20 | 10036 | 1984 | 0.1977 | 1143 | 0.1139 | 2124 | 0.2116 |

| | | | | | | | | |
|---|---|---|---|---|---|---|---|---|
| 1.5 | 5 | | | | | | | |
| 0 | 20 | 10036 | 2040 | 0.2033 | 1161 | 0.1157 | 2111 | 0.2103 |
| 4 | 5 | | | | | | | |
| 0 | 20 | 10036 | 2145 | 0.2137 | 1213 | 0.1209 | 2164 | 0.2156 |
| 3 | 5 | | | | | | | |
| 0 | 20 | 10036 | 1672 | 0.1666 | 250 | 0.0249 | 1074 | 0.1070 |
| -3.4 | 6.2 | | | | | | | |
| 0 | 20 | 10036 | 1593 | 0.1587 | 309 | 0.0308 | 1277 | 0.1272 |
| -5.9 | 5.2 | | | | | | | |
| 0 | 20 | 10036 | 1659 | 0.1653 | 388 | 0.0387 | 1462 | 0.1457 |
| 7.1 | 5.2 | | | | | | | |
| 0 | 20 | 10036 | 2962 | 0.2951 | 1696 | 0.1690 | 1783 | 0.1777 |
| 2.5 | 5 | | | | | | | |
| 0 | 20 | 10036 | 2895 | 0.2885 | 1106 | 0.1102 | 2212 | 0.2204 |
| 0.6 | 5.2 | | | | | | | |
| 0 | 20 | 10036 | 3162 | 0.3151 | 1167 | 0.1163 | 2390 | 0.2381 |
| 1.5 | 5 | | | | | | | |
| 0 | 20 | 10036 | 1793 | 0.1787 | 1061 | 0.1057 | 1968 | 0.1961 |
| 3.8 | 5.6 | | | | | | | |
| 0 | 20 | 10036 | 1830 | 0.1823 | 1101 | 0.1097 | 2057 | 0.2050 |
| 4.7 | 7.4 | | | | | | | |
| 0 | 20 | 10036 | 1929 | 0.1922 | 1084 | 0.1080 | 2079 | 0.2072 |
| 3.5 | 5 | | | | | | | |
| 0 | 20 | 10036 | 1393 | 0.1388 | 187 | 0.0186 | 918 | 0.0915 |
| -4.9 | 5.2 | | | | | | | |
| 0 | 20 | 10036 | 1415 | 0.1410 | 213 | 0.0212 | 1057 | 0.1053 |
| -3.4 | 5.2 | | | | | | | |
| 0 | 20 | 10036 | 1423 | 0.1418 | 260 | 0.0259 | 1276 | 0.1271 |
| -6.5 | 5 | | | | | | | |
| 0 | 20 | 10036 | 2981 | 0.2970 | 1720 | 0.1714 | 1978 | 0.1971 |
| 5.5 | 5 | | | | | | | |
| 0 | 20 | 10036 | 3157 | 0.3146 | 1432 | 0.1427 | 2339 | 0.2331 |
| 4.9 | 5 | | | | | | | |
| 0 | 20 | 10036 | 3153 | 0.3142 | 1598 | 0.1592 | 2252 | 0.2244 |
| 2.6 | 6 | | | | | | | |
| 0 | 20 | 10036 | 2385 | 0.2376 | 1371 | 0.1366 | 1812 | 0.1806 |
| 3.9 | 5 | | | | | | | |
| 0 | 20 | 10036 | 2288 | 0.2280 | 1391 | 0.1386 | 1814 | 0.1807 |
| -3.6 | 5 | | | | | | | |
| 0 | 20 | 10036 | 2288 | 0.2280 | 1317 | 0.1312 | 1946 | 0.1939 |
| -2.7 | 5.2 | | | | | | | |
| 0 | 20 | 10036 | 2134 | 0.2126 | 760 | 0.0757 | 1997 | 0.1990 |
| 0.1 | 5.2 | | | | | | | |
| 0 | 20 | 10036 | 2108 | 0.2100 | 881 | 0.0878 | 1997 | 0.1990 |
| -1.4 | 5 | | | | | | | |

| | | | | | | | | |
|---|---|---|---|---|---|---|---|---|
| 0 | 20 | 10036 | 2138 | 0.2130 | 905 | 0.0902 | 2006 | 0.1999 |
| -1.6 | 5.8 | | | | | | | |
| 0 | 20 | 10036 | 2935 | 0.2924 | 1786 | 0.1780 | 1986 | 0.1979 |
| 4.7 | 6.4 | | | | | | | |
| 0 | 20 | 10036 | 3170 | 0.3159 | 1445 | 0.1440 | 2245 | 0.2237 |
| 5.1 | 5.2 | | | | | | | |
| 0 | 20 | 10036 | 3149 | 0.3138 | 1628 | 0.1622 | 2235 | 0.2227 |
| 2.3 | 5.2 | | | | | | | |
| 0 | 20 | 10036 | 2396 | 0.2387 | 1383 | 0.1378 | 1815 | 0.1808 |
| -3.9 | 5 | | | | | | | |
| 0 | 20 | 10036 | 2259 | 0.2251 | 1382 | 0.1377 | 1915 | 0.1908 |
| -1 | 5 | | | | | | | |
| 0 | 20 | 10036 | 2320 | 0.2312 | 1412 | 0.1407 | 1937 | 0.1930 |
| -5.7 | 5.6 | | | | | | | |
| 0 | 20 | 10036 | 2115 | 0.2107 | 810 | 0.0807 | 1886 | 0.1879 |
| -6.6 | 5 | | | | | | | |
| 0 | 20 | 10036 | 2151 | 0.2143 | 847 | 0.0844 | 1869 | 0.1862 |
| -1.6 | 5 | | | | | | | |
| 0 | 20 | 10036 | 2157 | 0.2149 | 927 | 0.0924 | 1960 | 0.1953 |
| -0.3 | 5 | | | | | | | |
| 0 | 20 | 10036 | 3000 | 0.2989 | 1662 | 0.1656 | 1996 | 0.1989 |
| 3.9 | 5.2 | | | | | | | |
| 0 | 20 | 10036 | 3169 | 0.3158 | 1513 | 0.1508 | 2293 | 0.2285 |
| 5 | 5 | | | | | | | |
| 0 | 20 | 10036 | 3110 | 0.3099 | 1628 | 0.1622 | 2238 | 0.2230 |
| 2.6 | 5.4 | | | | | | | |
| 0 | 20 | 10036 | 2381 | 0.2372 | 1284 | 0.1279 | 1803 | 0.1797 |
| -7.2 | 5 | | | | | | | |
| 0 | 20 | 10036 | 2353 | 0.2345 | 1366 | 0.1361 | 1790 | 0.1784 |
| -6.7 | 5 | | | | | | | |
| 0 | 20 | 10036 | 2278 | 0.2270 | 1363 | 0.1358 | 1916 | 0.1909 |
| -6 | 5.2 | | | | | | | |
| 0 | 20 | 10036 | 2095 | 0.2087 | 761 | 0.0758 | 1879 | 0.1872 |
| -0.5 | 5 | | | | | | | |
| 0 | 20 | 10036 | 2060 | 0.2053 | 812 | 0.0809 | 1930 | 0.1923 |
| 0.5 | 5 | | | | | | | |
| 0 | 20 | 10036 | 2208 | 0.2200 | 856 | 0.0853 | 1860 | 0.1853 |
| -1 | 5.2 | | | | | | | |
| 0 | 20 | 10036 | 2828 | 0.2818 | 1660 | 0.1654 | 1860 | 0.1853 |
| 3.9 | 5 | | | | | | | |
| 0 | 20 | 10036 | 3109 | 0.3098 | 1269 | 0.1264 | 2155 | 0.2147 |
| 3.9 | 5 | | | | | | | |
| 0 | 20 | 10036 | 3118 | 0.3107 | 1473 | 0.1468 | 2053 | 0.2046 |
| 2.3 | 5 | | | | | | | |
| 0 | 20 | 10036 | 1763 | 0.1757 | 1048 | 0.1044 | 1430 | 0.1425 |

| | | | | | | | | |
|---|---|---|---|---|---|---|---|---|
| 3.4 | 5 | | | | | | | |
| 0 | 20 | 10036 | 2095 | 0.2087 | 1625 | 0.1619 | 1545 | 0.1539 |
| -3 | 8.2 | | | | | | | |
| 0 | 20 | 10036 | 1835 | 0.1828 | 1098 | 0.1094 | 1520 | 0.1515 |
| -4.5 | 5 | | | | | | | |
| 0 | 20 | 10036 | 1370 | 0.1365 | 500 | 0.0498 | 1341 | 0.1336 |
| -0.5 | 5.2 | | | | | | | |
| 0 | 20 | 10036 | 1432 | 0.1427 | 542 | 0.0540 | 1248 | 0.1244 |
| 0.4 | 5 | | | | | | | |
| 0 | 20 | 10036 | 1366 | 0.1361 | 581 | 0.0579 | 1315 | 0.1310 |
| 0.7 | 5.8 | | | | | | | |
| 0 | 20 | 10036 | 2736 | 0.2726 | 1635 | 0.1629 | 2065 | 0.2058 |
| 3.8 | 6.4 | | | | | | | |
| 0 | 20 | 10036 | 2857 | 0.2847 | 1539 | 0.1533 | 2096 | 0.2088 |
| 1.9 | 6.8 | | | | | | | |
| 0 | 20 | 10036 | 2739 | 0.2729 | 1546 | 0.1540 | 1893 | 0.1886 |
| 1.9 | 12.6 | | | | | | | |
| 0 | 20 | 10036 | 2279 | 0.2271 | 1270 | 0.1265 | 1824 | 0.1817 |
| -5.5 | 5 | | | | | | | |
| 0 | 20 | 10036 | 2261 | 0.2253 | 1251 | 0.1247 | 1858 | 0.1851 |
| -5.2 | 5.6 | | | | | | | |
| 0 | 20 | 10036 | 2338 | 0.2330 | 1308 | 0.1303 | 1828 | 0.1821 |
| -6.1 | 5.4 | | | | | | | |
| 0 | 20 | 10036 | 2723 | 0.2713 | 1170 | 0.1166 | 1869 | 0.1862 |
| 0.3 | 5 | | | | | | | |
| 0 | 20 | 10036 | 2732 | 0.2722 | 1156 | 0.1152 | 1850 | 0.1843 |
| -2.3 | 5 | | | | | | | |
| 0 | 20 | 10036 | 2650 | 0.2640 | 1131 | 0.1127 | 1890 | 0.1883 |
| -1.8 | 5.2 | | | | | | | |
| 0 | 20 | 10036 | 2766 | 0.2756 | 1634 | 0.1628 | 2047 | 0.2040 |
| 3.1 | 7.4 | | | | | | | |
| 0 | 20 | 10036 | 2716 | 0.2706 | 1644 | 0.1638 | 2119 | 0.2111 |
| 2.9 | 8.2 | | | | | | | |
| 0 | 20 | 10036 | 2767 | 0.2757 | 1646 | 0.1640 | 1759 | 0.1753 |
| 1.8 | 12.8 | | | | | | | |
| 0 | 20 | 10036 | 2267 | 0.2259 | 1310 | 0.1305 | 1788 | 0.1782 |
| -5.6 | 5 | | | | | | | |
| 0 | 20 | 10036 | 2266 | 0.2258 | 1339 | 0.1334 | 1855 | 0.1848 |
| -6.2 | 5.2 | | | | | | | |
| 0 | 20 | 10036 | 2250 | 0.2242 | 1380 | 0.1375 | 1769 | 0.1763 |
| -5 | 5.4 | | | | | | | |
| 0 | 20 | 10036 | 2720 | 0.2710 | 1169 | 0.1165 | 1861 | 0.1854 |
| 1.2 | 5 | | | | | | | |
| 0 | 20 | 10036 | 2831 | 0.2821 | 1150 | 0.1146 | 1853 | 0.1846 |
| -6.4 | 5 | | | | | | | |

| | | | | | | | | |
|---|---|---|---|---|---|---|---|---|
| 0 | 20 | 10036 | 2605 | 0.2596 | 1095 | 0.1091 | 1888 | 0.1881 |
| -1.8 | 5 | | | | | | | |
| 0 | 20 | 10036 | 2766 | 0.2756 | 1633 | 0.1627 | 1972 | 0.1965 |
| 4 | 8 | | | | | | | |
| 0 | 20 | 10036 | 2768 | 0.2758 | 1553 | 0.1547 | 2025 | 0.2018 |
| 3.6 | 10 | | | | | | | |
| 0 | 20 | 10036 | 2824 | 0.2814 | 1581 | 0.1575 | 1785 | 0.1779 |
| 2 | 12.4 | | | | | | | |
| 0 | 20 | 10036 | 2206 | 0.2198 | 1308 | 0.1303 | 1686 | 0.1680 |
| -5.4 | 5 | | | | | | | |
| 0 | 20 | 10036 | 2311 | 0.2303 | 1355 | 0.1350 | 1828 | 0.1821 |
| -5.9 | 5 | | | | | | | |
| 0 | 20 | 10036 | 2244 | 0.2236 | 1313 | 0.1308 | 1780 | 0.1774 |
| -4.4 | 5.6 | | | | | | | |
| 0 | 20 | 10036 | 2194 | 0.2186 | 1272 | 0.1267 | 1705 | 0.1699 |
| -6.9 | 5 | | | | | | | |
| 0 | 20 | 10036 | 2602 | 0.2593 | 1161 | 0.1157 | 1745 | 0.1739 |
| -1.6 | 6 | | | | | | | |
| 0 | 20 | 10036 | 2494 | 0.2485 | 1072 | 0.1068 | 1808 | 0.1802 |
| -1.9 | 5 | | | | | | | |
| 0 | 20 | 10036 | 2524 | 0.2515 | 1555 | 0.1549 | 1810 | 0.1804 |
| 3.7 | 5.8 | | | | | | | |
| 0 | 20 | 10036 | 2625 | 0.2616 | 1468 | 0.1463 | 1916 | 0.1909 |
| 3.1 | 9.2 | | | | | | | |
| 0 | 20 | 10036 | 2634 | 0.2625 | 1531 | 0.1526 | 1712 | 0.1706 |
| 2.8 | 12 | | | | | | | |
| 0 | 20 | 10036 | 1358 | 0.1353 | 776 | 0.0773 | 1037 | 0.1033 |
| -5.5 | 5 | | | | | | | |
| 0 | 20 | 10036 | 1321 | 0.1316 | 749 | 0.0746 | 982 | 0.0978 |
| 3.6 | 5.2 | | | | | | | |
| 0 | 20 | 10036 | 1391 | 0.1386 | 721 | 0.0718 | 1067 | 0.1063 |
| 6.1 | 5.2 | | | | | | | |
| 0 | 20 | 10036 | 1155 | 0.1151 | 623 | 0.0621 | 710 | 0.0707 |
| -1.9 | 5.2 | | | | | | | |
| 0 | 20 | 10036 | 1082 | 0.1078 | 566 | 0.0564 | 729 | 0.0726 |
| 4.2 | 5.4 | | | | | | | |
| 0 | 20 | 10036 | 1253 | 0.1249 | 674 | 0.0672 | 736 | 0.0733 |
| -4.25 | 5 | | | | | | | |

| RML (ID=30) | | RUL (ID=38) | | **CoV** | **Total DF** |
|---|---|---|---|---|---|
| **DPN-RM** | **DF-RM** | **DPN-RU** | **DF-RU** | | |
| 1249 | 0.1245 | 1448 | 0.1443 | **0.370464** | **0.9270** |
| 1306 | 0.1301 | 1603 | 0.1597 | **0.394868** | **0.9339** |
| 1305 | 0.1300 | 1149 | 0.1145 | **0.513321** | **0.9365** |
| 1406 | 0.1401 | 1986 | 0.1979 | **0.23448** | **0.8781** |
| 1313 | 0.1308 | 2060 | 0.2053 | **0.286401** | **0.8754** |
| 1424 | 0.1419 | 1951 | 0.1944 | **0.240214** | **0.8896** |
| 510 | 0.0508 | 671 | 0.0669 | **0.660528** | **0.4240** |
| 622 | 0.0620 | 883 | 0.0880 | **0.596437** | **0.4833** |
| 753 | 0.0750 | 989 | 0.0985 | **0.515988** | **0.5229** |
| 1254 | 0.1250 | 1458 | 0.1453 | **0.359549** | **0.9283** |
| 1317 | 0.1312 | 1589 | 0.1583 | **0.402878** | **0.9341** |
| 1299 | 0.1294 | 1151 | 0.1147 | **0.499745** | **0.9379** |
| 1461 | 0.1456 | 1944 | 0.1937 | **0.240731** | **0.8756** |
| 1388 | 0.1383 | 1962 | 0.1955 | **0.251283** | **0.8796** |
| 1416 | 0.1411 | 1982 | 0.1975 | **0.248973** | **0.8813** |
| 433 | 0.0431 | 596 | 0.0594 | **0.806224** | **0.4286** |
| 606 | 0.0604 | 818 | 0.0815 | **0.595562** | **0.4712** |
| 785 | 0.0782 | 993 | 0.0989 | **0.479342** | **0.5238** |
| 1227 | 0.1223 | 1418 | 0.1413 | **0.371096** | **0.9263** |
| 1315 | 0.1310 | 1660 | 0.1654 | **0.401994** | **0.9392** |
| 1285 | 0.1280 | 1175 | 0.1171 | **0.509047** | **0.9392** |
| 1563 | 0.1557 | 1961 | 0.1954 | **0.228355** | **0.8744** |

| | | | | | |
|---|---|---|---|---|---|
| 1419 | 0.1414 | 1978 | 0.1971 | **0.244009** | **0.8678** |
| 1474 | 0.1469 | 1854 | 0.1847 | **0.236477** | **0.8818** |
| 470 | 0.0468 | 679 | 0.0677 | **0.676292** | **0.4130** |
| 629 | 0.0627 | 861 | 0.0858 | **0.546289** | **0.4652** |
| 779 | 0.0776 | 937 | 0.0934 | **0.493764** | **0.5206** |
| 1111 | 0.1107 | 1310 | 0.1305 | **0.406107** | **0.8830** |
| 1240 | 0.1236 | 1610 | 0.1604 | **0.409083** | **0.9030** |
| 1246 | 0.1242 | 1122 | 0.1118 | **0.504969** | **0.9054** |
| 1347 | 0.1342 | 1754 | 0.1748 | **0.233952** | **0.7895** |
| 1313 | 0.1308 | 1752 | 0.1746 | **0.243586** | **0.8024** |
| 1321 | 0.1316 | 1708 | 0.1702 | **0.25581** | **0.8092** |
| 427 | 0.0425 | 590 | 0.0588 | **0.666261** | **0.3502** |
| 527 | 0.0525 | 713 | 0.0710 | **0.594038** | **0.3911** |
| 664 | 0.0662 | 777 | 0.0774 | **0.537024** | **0.4384** |
| 1193 | 0.1189 | 1505 | 0.1500 | **0.363597** | **0.9343** |
| 1174 | 0.1170 | 1422 | 0.1417 | **0.43508** | **0.9490** |
| 1086 | 0.1082 | 1433 | 0.1428 | **0.428665** | **0.9488** |
| 1243 | 0.1239 | 2032 | 0.2025 | **0.26604** | **0.8811** |
| 1327 | 0.1322 | 2108 | 0.2100 | **0.238127** | **0.8896** |
| 1312 | 0.1307 | 2069 | 0.2062 | **0.250708** | **0.8900** |
| 1468 | 0.1463 | 1879 | 0.1872 | **0.336881** | **0.8208** |
| 1491 | 0.1486 | 1781 | 0.1775 | **0.297232** | **0.8228** |

| | | | | | |
|---|---|---|---|---|---|
| 1379 | 0.1374 | 1819 | 0.1812 | **0.306443** | **0.8217** |
| 1138 | 0.1134 | 1503 | 0.1498 | **0.361312** | **0.9314** |
| 1143 | 0.1139 | 1534 | 0.1528 | **0.42638** | **0.9503** |
| 1095 | 0.1091 | 1356 | 0.1351 | **0.43329** | **0.9429** |
| 1253 | 0.1249 | 2077 | 0.2070 | **0.266374** | **0.8892** |
| 1300 | 0.1295 | 2143 | 0.2135 | **0.243186** | **0.8967** |
| 1303 | 0.1298 | 2039 | 0.2032 | **0.239277** | **0.8979** |
| 1486 | 0.1481 | 1836 | 0.1829 | **0.312915** | **0.8104** |
| 1397 | 0.1392 | 1966 | 0.1959 | **0.319686** | **0.8200** |
| 1430 | 0.1425 | 1785 | 0.1779 | **0.293867** | **0.8229** |
| 1142 | 0.1138 | 1506 | 0.1501 | **0.379742** | **0.9273** |
| 1102 | 0.1098 | 1399 | 0.1394 | **0.441641** | **0.9442** |
| 1139 | 0.1135 | 1361 | 0.1356 | **0.418962** | **0.9442** |
| 1234 | 0.1230 | 2027 | 0.2020 | **0.280765** | **0.8698** |
| 1199 | 0.1195 | 2087 | 0.2080 | **0.274039** | **0.8763** |
| 1246 | 0.1242 | 2025 | 0.2018 | **0.250832** | **0.8796** |
| 1415 | 0.1410 | 1806 | 0.1800 | **0.330024** | **0.7927** |
| 1397 | 0.1392 | 1768 | 0.1762 | **0.315462** | **0.7938** |
| 1362 | 0.1357 | 1778 | 0.1772 | **0.32193** | **0.8035** |
| 996 | 0.0992 | 1385 | 0.1380 | **0.393105** | **0.8698** |
| 1019 | 0.1015 | 1477 | 0.1472 | **0.466253** | **0.8997** |
| 1049 | 0.1045 | 1308 | 0.1303 | **0.457577** | **0.8969** |
| 906 | 0.0903 | 1627 | 0.1621 | **0.271684** | **0.6750** |

| | | | | | |
|---|---|---|---|---|---|
| 1216 | 0.1212 | 1436 | 0.1431 | **0.205048** | **0.7889** |
| 994 | 0.0990 | 1650 | 0.1644 | **0.254089** | **0.7072** |
| 901 | 0.0898 | 1176 | 0.1172 | **0.343325** | **0.5269** |
| 915 | 0.0912 | 1178 | 0.1174 | **0.324845** | **0.5296** |
| 870 | 0.0867 | 1246 | 0.1242 | **0.314451** | **0.5359** |
| 1262 | 0.1257 | 1645 | 0.1639 | **0.300773** | **0.9309** |
| 1482 | 0.1477 | 1501 | 0.1496 | **0.314237** | **0.9441** |
| 1801 | 0.1795 | 1512 | 0.1507 | **0.262036** | **0.9457** |
| 1239 | 0.1235 | 2017 | 0.2010 | **0.266354** | **0.8598** |
| 1264 | 0.1259 | 2034 | 0.2027 | **0.263912** | **0.8637** |
| 1210 | 0.1206 | 1922 | 0.1915 | **0.270056** | **0.8575** |
| 1308 | 0.1303 | 1219 | 0.1215 | **0.396941** | **0.8259** |
| 1330 | 0.1325 | 1267 | 0.1262 | **0.39135** | **0.8305** |
| 1358 | 0.1353 | 1327 | 0.1322 | **0.36817** | **0.8326** |
| 1166 | 0.1162 | 1690 | 0.1684 | **0.319938** | **0.9270** |
| 1523 | 0.1518 | 1530 | 0.1525 | **0.269796** | **0.9498** |
| 1787 | 0.1781 | 1466 | 0.1461 | **0.269996** | **0.9391** |
| 1212 | 0.1208 | 1917 | 0.1910 | **0.257692** | **0.8464** |
| 1191 | 0.1187 | 1959 | 0.1952 | **0.259453** | **0.8579** |
| 1215 | 0.1211 | 1939 | 0.1932 | **0.244804** | **0.8522** |
| 1226 | 0.1222 | 1252 | 0.1248 | **0.40296** | **0.8198** |
| 1211 | 0.1207 | 1256 | 0.1251 | **0.429459** | **0.8271** |

| | | | | | |
|---|---|---|---|---|---|
| 1276 | 0.1271 | 1400 | 0.1395 | **0.367907** | **0.8234** |
| 1185 | 0.1181 | 1664 | 0.1658 | **0.31826** | **0.9187** |
| 1580 | 0.1574 | 1529 | 0.1524 | **0.280959** | **0.9421** |
| 1754 | 0.1748 | 1480 | 0.1475 | **0.28637** | **0.9390** |
| 1155 | 0.1151 | 1888 | 0.1881 | **0.258961** | **0.8213** |
| 1105 | 0.1101 | 1918 | 0.1911 | **0.280044** | **0.8486** |
| 1133 | 0.1129 | 1939 | 0.1932 | **0.2707** | **0.8379** |
| 1191 | 0.1187 | 1869 | 0.1862 | **0.254194** | **0.8201** |
| 1198 | 0.1194 | 1280 | 0.1275 | **0.38106** | **0.7957** |
| 1236 | 0.1232 | 1301 | 0.1296 | **0.366032** | **0.7883** |
| 997 | 0.0993 | 1471 | 0.1466 | **0.335083** | **0.8327** |
| 1336 | 0.1331 | 1483 | 0.1478 | **0.298919** | **0.8796** |
| 1686 | 0.1680 | 1177 | 0.1173 | **0.308509** | **0.8709** |
| 615 | 0.0613 | 1096 | 0.1092 | **0.29611** | **0.4864** |
| 604 | 0.0602 | 1229 | 0.1225 | **0.312548** | **0.4867** |
| 608 | 0.0606 | 1240 | 0.1236 | **0.332245** | **0.5009** |
| 440 | 0.0438 | 621 | 0.0619 | **0.376992** | **0.3536** |
| 556 | 0.0554 | 857 | 0.0854 | **0.290002** | **0.3776** |
| 502 | 0.0500 | 585 | 0.0583 | **0.393113** | **0.3737** |
