## Appendix B: Machine learning performance and model selection for "Computational Fluid Particle Dynamics-Informed Machine Learning Prototype for a User-Centered Smart Inhaler Enabling Uniform Drug Delivery to Small Airways"

| Model | Model Abbreviation | BestEpoch | MSE_Test | MAE_Test | MSE_Test_$x_c$ | MSE_Test_$y_c$ | MSE_Test_$d_n$ | MAE_Test_$x_c$ | MAE_Test_$y_c$ | MAE_Test_$d_n$ | MSE_Val | MAE_Val | MSE_Val_$x_c$ | MSE_Val_$y_c$ | MSE_Val_$d_n$ | MAE_Val_$x_c$ | MAE_Val_$y_c$ | MAE_Val_$d_n$ |
|---|---|---|---|---|---|---|---|---|---|---|---|---|---|---|---|---|---|---|
| RegressionModel | RM | 1323 | 7.453 | 1.831 | 14.267 | 6.464 | 1.627 | 2.944 | 1.632 | 0.919 | 3.036 | 1.422 | 6.001 | 2.810 | 0.298 | 2.415 | 1.369 | 0.483 |
| **RegressionMode_Small** | **RM_S** | 3620 | 7.425 | 1.918 | 14.031 | **6.345** | 1.900 | 2.871 | 1.898 | 0.986 | 4.778 | 1.756 | 11.680 | 2.527 | 0.126 | 3.391 | 1.547 | 0.328 |
| RegressionModel_Medium | RM_M | 398 | 6.975 | 1.751 | 11.693 | 7.438 | 1.793 | 2.591 | 1.768 | 0.894 | 4.719 | 1.705 | 12.020 | 1.904 | 0.233 | 3.463 | 1.173 | 0.477 |
| **RegressionModel_Deep** | **RM_D** | 147 | **6.656** | 1.661 | 11.088 | 7.495 | 1.385 | 2.436 | 1.800 | 0.748 | 5.987 | 1.909 | 15.284 | 2.399 | 0.277 | 3.896 | 1.378 | 0.453 |
| RegressionModelWithDropout | RMD | 1021 | 6.878 | 1.824 | 12.521 | 6.581 | 1.531 | 2.800 | 1.876 | 0.797 | 4.713 | 1.796 | 10.918 | 3.031 | 0.189 | 3.304 | 1.683 | 0.401 |
| RegressionModelWithDropout_Small | RMD_S | 14885 | 7.583 | 1.928 | 13.329 | 7.641 | 1.779 | 2.745 | 2.162 | 0.878 | 5.920 | 1.902 | 14.753 | 2.830 | 0.178 | 3.841 | 1.444 | 0.421 |
| RegressionModelWithDropout_Medium | RMD_M | 861 | 6.748 | 1.821 | 11.997 | 6.373 | 1.874 | 2.779 | 1.827 | 0.856 | 5.690 | 1.852 | 14.493 | 2.403 | 0.173 | 3.795 | 1.380 | 0.381 |
| **RegressionModelWithDropout_Deep** | **RMD_D** | 830 | **6.697** | 1.833 | 11.792 | 6.877 | 1.422 | 2.773 | 1.999 | 0.726 | 5.150 | 1.750 | 13.269 | 2.064 | 0.116 | 3.606 | 1.331 | 0.314 |
| CNNRegressor | CNN_B | 1265 | 8.263 | 2.031 | 15.869 | 7.067 | 1.854 | 3.203 | 1.969 | 0.920 | 5.772 | 1.760 | 16.508 | 0.623 | 0.184 | 4.062 | 0.788 | 0.429 |
| CNNRegressor_Small | CNN_S | 2723 | 8.272 | 2.038 | 15.720 | 7.378 | 1.717 | 3.057 | 2.138 | 0.920 | 5.060 | 1.479 | 14.533 | 0.375 | 0.272 | 3.591 | 0.441 | 0.406 |
| CNNRegressor_Medium | CNN_M | 632 | 7.643 | 1.900 | 14.070 | 7.298 | 1.559 | 2.863 | 2.007 | 0.832 | 5.096 | 1.716 | 14.070 | 0.953 | 0.265 | 3.703 | 0.935 | 0.510 |
| CNNRegressor_Deep | CNN_D | 407 | 7.929 | 1.945 | 14.375 | 8.112 | 1.298 | 2.926 | 2.198 | 0.710 | 7.248 | 2.094 | 18.403 | 3.153 | 0.187 | 4.290 | 1.560 | 0.433 |
| TransformerModel | TF | 20 | 8.693 | 2.076 | 14.074 | 9.718 | 2.286 | 2.800 | 2.406 | 1.020 | 6.508 | 2.050 | 16.600 | 2.531 | 0.393 | 4.071 | 1.571 | 0.509 |
| TransformerRegressor_Small | TF_S | 72 | 7.077 | 1.824 | 12.398 | 7.110 | 1.724 | 2.688 | 1.887 | 0.897 | 5.052 | 1.687 | 13.720 | 1.335 | 0.099 | 3.683 | 1.116 | 0.263 |
| TransformerRegressor_Medium | TF_M | 43 | 6.960 | 1.749 | 12.393 | 7.117 | 1.370 | 2.730 | 1.763 | 0.755 | 2.788 | 1.239 | 7.851 | 0.248 | 0.264 | 2.771 | 0.465 | 0.481 |
| **TransformerRegressor_Deep** | **TF_D** | 78 | **6.445** | 1.613 | **10.998** | 7.083 | **1.255** | 2.552 | 1.576 | 0.711 | 5.243 | 1.903 | 11.372 | 4.136 | 0.222 | 3.365 | 1.872 | 0.471 |
