## Appendix C: Comparisons of Performances between ML-Driven TDD and CFPD-TDD Strategies on Cross-Validation Cases for "Computational Fluid Particle Dynamics-Informed Machine Learning Prototype for a User-Centered Smart Inhaler Enabling Uniform Drug Delivery to Small Airways"

| | Models | PFR [L/min] | VM [m/s] | PD [mm] | Z [mm] | PRT [s] | X-C [mm] | Y-C [mm] | ND [mm] | TRPN | DPN-LL | DF-LL | DPN-LU | DF-LU | DPN-RL | DF-RL | DPN-RM | DF-RM | DPN-RU | DF-RU | CoV | DF |
|---|---|---|---|---|---|---|---|---|---|---|---|---|---|---|---|---|---|---|---|---|---|---|
| **Case I** | CFPD -FMD | 33 | 1.7554 | 0.001 | 1 | 0.5 | 0.0000 | 0.0000 | 20.0000 | 10036 | 2154 | 0.2146 | 576 | 0.0574 | 1546 | 0.1540 | 820 | 0.0817 | 823 | 0.0820 | 0.55142 | 0.5898 |
| | **CFPD-TDD** | | | | | | 7.0000 | -1.4000 | 5.6000 | 10036 | 2097 | 0.2089 | 718 | 0.0715 | 1620 | 0.1614 | 1282 | 0.1277 | 1419 | 0.1414 | **0.35207** | 0.7110 |
| | MixModel | | | | | | 3.4127 | -3.3118 | 5.1432 | 10036 | 2339 | 0.2331 | 718 | 0.0715 | 1529 | 0.1524 | 1112 | 0.1108 | 1120 | 0.1116 | 0.4518 | 0.6794 |
| | RM_D | | | | | | 3.4345 | -3.4848 | 4.8790 | 10036 | 2342 | 0.2334 | 717 | 0.0714 | 1564 | 0.1558 | 1159 | 0.1155 | 1149 | 0.1145 | **0.44189** | 0.6906 |
| | RMD_D | | | | | | 2.8781 | -2.1906 | 5.1529 | 10036 | 2311 | 0.2303 | 669 | 0.0667 | 1419 | 0.1414 | 949 | 0.0946 | 1097 | 0.1093 | 0.49039 | 0.6422 |
| | TF_D | | | | | | 3.4127 | -4.0668 | 5.1432 | 10036 | 2490 | 0.2481 | 723 | 0.0720 | 1653 | 0.1647 | 1078 | 0.1074 | 1093 | 0.1089 | 0.49074 | 0.7012 |
| **Case II** | CFPD -FMD | 46 | 2.4470 | 0.0005 | 20 | 0 | 0.0000 | 0.0000 | 20.0000 | 10036 | 3164 | 0.3153 | 1460 | 0.1455 | 2311 | 0.2303 | 1198 | 0.1194 | 1377 | 0.1372 | 0.43407 | 0.9476 |
| | **CFPD-TDD** | | | | | | 5.0000 | 5.0000 | 5.0000 | 10036 | 1660 | 0.1654 | 1607 | 0.1601 | 2935 | 0.2924 | 1780 | 0.1774 | 1131 | 0.1127 | **0.36708** | 0.9080 |
| | MixModel | | | | | | 1.8576 | 3.8175 | 5.4851 | 10036 | 2189 | 0.2181 | 1262 | 0.1257 | 3136 | 0.3125 | 1051 | 0.1047 | 1478 | 0.1473 | 0.46602 | 0.9083 |
| | RM_D | | | | | | 0.7744 | 4.3281 | 5.1844 | 10036 | 2644 | 0.2635 | 1319 | 0.1314 | 2449 | 0.2440 | 1468 | 0.1463 | 1146 | 0.1142 | **0.38206** | 0.8994 |
| | RMD_D | | | | | | 1.9100 | 3.1282 | 5.1805 | 10036 | 1686 | 0.1680 | 1028 | 0.1024 | 4212 | 0.4197 | 746 | 0.0743 | 1761 | 0.1755 | 0.72589 | 0.9399 |
| | TF_D | | | | | | 1.8576 | 3.9225 | 5.4851 | 10036 | 2354 | 0.2346 | 1248 | 0.1244 | 3027 | 0.3016 | 1067 | 0.1063 | 1457 | 0.1452 | 0.45437 | 0.9120 |

| **Note:** |
|---|
| **PFR:** Peak Flow Rate |
| **VM:** Velocity of Magnitude |
| **PD:** Particle Diameter |
| **Z:** Z-Coordinate |
| **PRT:** Particle Release Time |
| **X-C:** Optimal Nozzle Center X-Coordinate |
| **Y-C:** Optimal Nozzle Y-Coordinate |
| **ND:** Optimal Nozzle Diameter |
| **DPN:** Deposition Particle Number |
| **TRPN:** Total Release Particle Number |
| **DF:** Deposition Fraction |
| **LL:**Left Lower Outlets |
| **LU:** Left Upper Outlets |
| **RL:** Right Lower Outlets |
| **RM:** Right Middle Outlets |
| **RU:** Right Upper Outlets |
| **CoV:** Coefficient of Variation |
